# Golden Mutagenesis: An efficient multi-site-saturation mutagenesis approach by Golden Gate cloning with automated primer design

**DOI:** 10.1101/453621

**Authors:** Pascal Püllmann, Chris Ulpinnis, Sylvestre Marillonnet, Ramona Gruetzner, Steffen Neumann, Martin J. Weissenborn

## Abstract

Site-directed methods for the generation of genetic diversity are essential tools in the field of directed enzyme evolution. The Golden Gate cloning technique has been proven to be an efficient tool for a variety of cloning setups. The utilization of restriction enzymes which cut outside of their recognition domain allows the assembly of multiple gene fragments obtained by PCR amplification without altering the open reading frame of the reconstituted gene. We have developed a protocol, termed Golden Mutagenesis that allows the rapid, straightforward, reliable and inexpensive construction of mutagenesis libraries. One to five amino acid positions within a coding sequence could be altered simultaneously using a protocol which can be performed within one day. To facilitate the implementation of this technique, a software library and web application for automated primer design and for the graphical evaluation of the randomization success based on the sequencing results was developed. This allows facile primer design and application of Golden Mutagenesis also for laboratories, which are not specialized in molecular biology.

---

Directed evolution endeavors require highly efficient molecular cloning techniques to simultaneously alter multiple residues in a rapid, reproducible and cost-effective manner. It is moreover desirable to avoid nucleobase distribution bias in the created library and ideally produce statistical distributions depending on the chosen degeneracy. In recent years, targeted combinatorial approaches for directed evolution such as CASTing^1,2^ or iterative saturation mutagenesis (ISM)^3,4^ have evolved as successful and widely applied techniques for protein engineering using “smart” directed evolution. A general requirement for these approaches is the ability to efficiently alter specific protein residues in a simultaneous manner.

Commonly employed techniques for single site-saturation mutagenesis include, amongst others, MOD-PCR (Mutagenic Oligonucleotide-Directed PCR Amplification),^5^ Codon Cassette Mutagenesis,^6^ Overlap Extension PCR,^7^ Megaprimer PCR^8^ and the commercial kit QuikChange^®^.^9^ However, the successful generation of high-quality libraries of gene sequences containing multiple randomization sites often requires more sophisticated methods to achieve a high success rate. Recent developments to address this issue include the introduction of Omnichange,^10^ Darwin Assembly,^11^ and ISOR.^12^

To facilitate the design of primers utilizing various mutagenesis techniques, computer tools were developed amongst others for Gibson Assembly (https://nebuilder.neb.com/#!/), gene domestication and assembly for Golden Gate cloning^13^ and primer design for cloning and site-saturation mutagenesis using GeneGenie.^14^ The use of gene synthesis as a mutagenic tool in directed evolution approaches was carefully assessed by Reetz and co-workers.^15^ By using high-fidelity on-chip gene synthesis and cloning by overlap extension PCR followed by classical type II restriction digest and gene ligation into the expression vector, they were able to obtain an excellent 97 % ratio of the overall possible gene diversity. They concluded that the application of gene synthesis is likely to become a powerful alternative to established PCR-based randomization protocols, if the overall costs of this type of library creation continue to drop. Also, the combination of high fidelity DNA synthesis of mutagenized DNA fragments with efficient and seamless cloning techniques such as Golden Gate cloning or Gibson Assembly could represent an interesting next developmental step.

Golden Gate cloning was introduced in 2008^16^ and is based on the use of type IIS restriction enzymes. This subclass of restriction enzymes is defined by their ability to cleave double-stranded DNA templates outside of their recognition site. A remarkable feature of the technique is that it allows to perform restriction and ligation reactions in a time and cost saving one-pot reaction setup with an overall efficiency of correct assembly close to 100 %.^17-23^ Since the type IIS recognition sequence is liberated from the insert upon successful restriction, a correctly integrated insert cannot be digested further—this feature enables an efficient one-pot restriction ligation setup leading to steadily increasing number of recombinant constructs in the course of the setup. In the case of classical type II enzymes, which digest double-stranded DNA within their recognition site, the site is restored upon ligation into the expression plasmid and therefore creating a competition scenario between restriction and ligation, if performed in a one-pot scenario and leading to overall low efficiencies of recombination. An increasingly large number of Golden Gate compatible plasmids is furthermore accessible through the non-profit plasmid repository Addgene.

Arguably the most valuable feature of the Golden Gate cloning technique is that it allows highly specific assembly of several gene fragments, which is mediated through the generation of four base pair overhangs that flank the fragments after restriction. 240 unique overhangs (4^4^ – 16 palindromic sequences) can be employed to join adjacent fragments, therefore enabling broad applicability in the field of multiple site-directed mutagenesis. Correct assembly occurs in a seamless manner, *i.e.* frameshift mutations are avoided and the original open reading frame of the target gene is restored. Furthermore, the conducted blue/orange against white bacterial colony color screening enables the distinction of negative events (no restriction/ligation) and hence save screening effort, since it allows the exclusion of those events for subsequent testing. In contrast to other widespread mutagenesis techniques like QuikChange^®^, amplification of the acceptor plasmid backbone via PCR is not necessary for Golden Gate cloning. This eliminates the risk of introducing unwanted mutations within the plasmid backbone. However, so far only very few examples were reported employing Golden Gate cloning for mutagenesis in the context of directed evolution approaches.^16,24,25^

We have developed and optimized a hands-on protocol for the implementation of Golden Gate-based Mutagenesis (coined Golden Mutagenesis) in any laboratory focusing on rational or random protein engineering. The entire Golden Mutagenesis protocol requires solely three enzymes (BsaI, BbsI and a DNA ligase) and one to two different plasmids (one cloning and one *E. coli* expression plasmid)—a Golden Gate compatible pET28b based expression vector was constructed and deposited at Addgene. The success of the respective Golden Gate digestion-ligation approach can be directly observed and estimated on the plate due to the utilized blue (LacZ; pAGM9121) or orange (CRed; pAGM22082_CRed) selection markers. Furthermore, an open source web tool was developed to facilitate primer design and analysis of the created randomized library, which is accessible at https://msbi.ipb-halle.de/GoldenMutagenesisWeb/. This tool allows the user to upload any gene sequence of interest, select a number of protein residues that shall be mutagenized and select a suitable cloning/expression vector depending on the desired cloning task. All required primers for Golden Gate cloning are automatically designed after specifying the desired codon degeneracies (*e.g.* NDT, NNK) and differences in melting temperatures of the corresponding primer pairs are minimized and set to a default T_m_ value of 60 °C. Following physical construction of the randomization library and sequencing of a pooled library, consisting of > n distinct bacterial colonies (depending on the number of randomization sites and the chosen degeneracy), the generated randomizations are assessed (.ab1 file format) and illustrated as nucleobase distributions in pie charts. Herein, we present a straightforward technique for multiple-site saturations aided by a web tool for primer design and sequencing analysis.

## General concept of Golden Mutagenesis

Extensive information about the conceptual basis of Golden Gate cloning can be found in detail elsewhere.^17,18,26^ Briefly, type IIS restriction enzymes (mostly BsaI or BbsI) are utilized, which cut double-stranded DNA molecules outside of their recognition site. This characteristic feature allows the assembly of DNA fragments in a seamless manner.

Golden Mutagenesis utilizes this above-mentioned cloning and assembly strategy enabling efficient parallel one-pot restriction-ligation procedures. PCR fragments are generated, which carry the terminal type IIS recognition sites and introduce additional specific randomization sites outside of the binding primer sequence. The general structure of the designed oligonucleotides follows this scheme: type IIS recognition site, specified four bp overhang, randomization site and template binding sequence (5’to 3’ direction). The generated PCR products are then reassembled in a target expression vector and the resulting genetic library is directly transformed into an *E. coli* expression strain (Figure 1, left). Direct Golden Gate assembly of multiple PCR fragments, which exhibit several parallel randomization sites may become less efficient with increasing gene fragment numbers and complexity levels. This is due to limitations in ligation efficiency and the generally lower transformation competency of *E. coli* expression strains, if compared to classical cloning strains. Therefore as second strategy was developed were individual gene fragments are subcloned into a cloning vector (pAGM9121) and transformed into an *E. coli* cloning strain. In order to preserve the introduced degeneracy the plasmid DNA preparations of the resulting single gene fragment constructs, are therefore prepared as libraries by direct inoculation of the transformation mixture into a liquid culture. The subcloned gene fragment libraries are then assembled into the final expression vector in the course of a second Golden Gate reaction—no additional PCR step is required (Figure 1, right). This two-step procedure also opens interesting possibilities in directed evolution approaches like iterative saturation mutagenesis (ISM) by fragmenting the gene into subunits.^1-4^ A gene fragment 1 could be subjected to targeted mutagenesis and subcloned in the cloning vector and finally assemble with the unmodified wildtype gene fragments 2 and 3 in the expression vector in a second Golden Gate reaction. In a subsequent round, fragment 2 can be subjected to mutagenesis and combined with the before optimized gene fragment 1 and fragment 3. Within this two-step Golden Gate subcloning approach different gene regions can be easily assigned and resulting libraries screened consecutively as envisioned in CASTing and ISM approaches. This offers an efficient approach for a controlled, hierarchical mutagenesis approach that was employed in previous studies in a similar method.^24^

**Figure 1.**
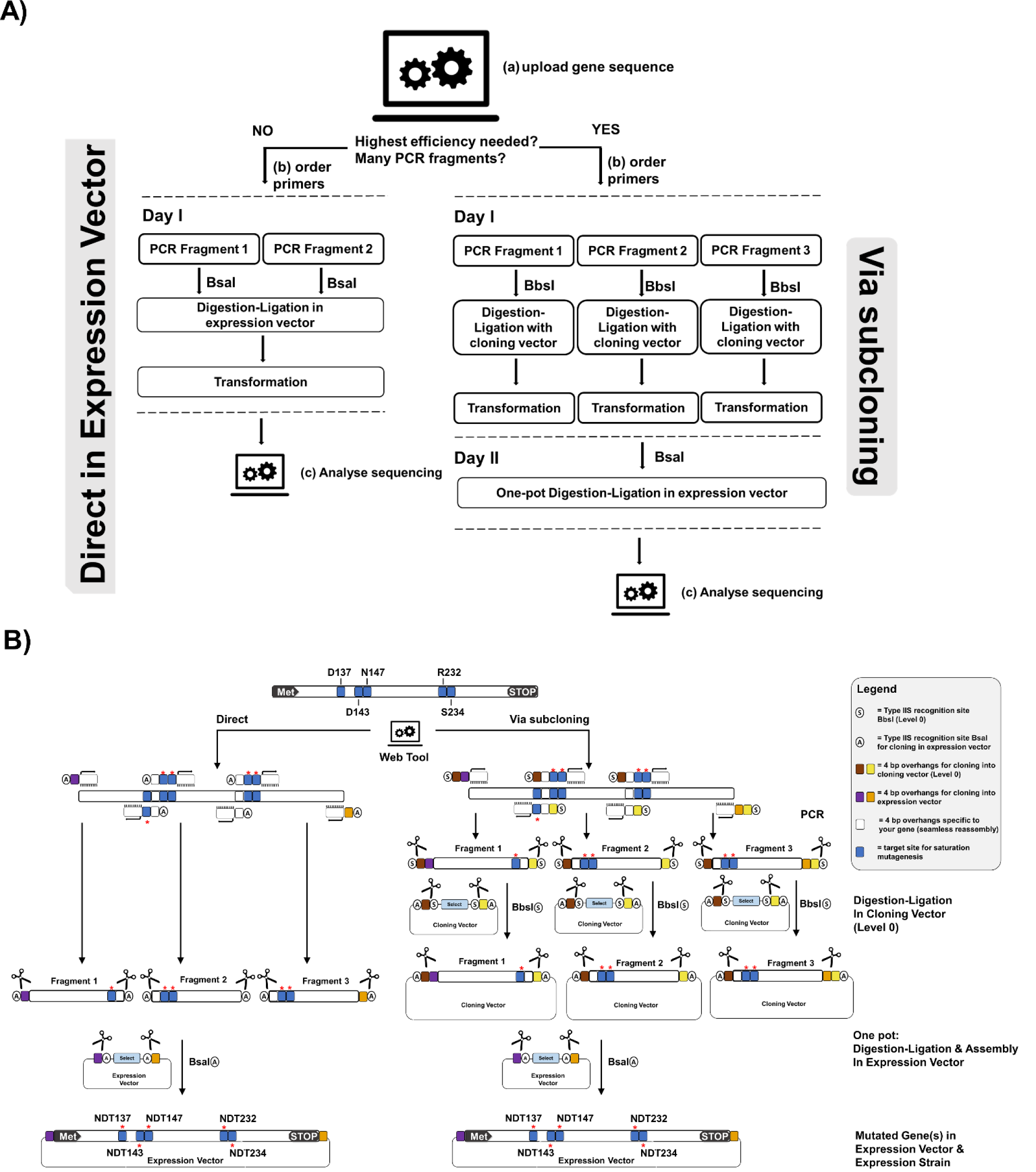
Setup for Golden Mutagenesis. **A)** Schematic overview of the workflow. After uploading a gene of interest into the web tool, the user can decide whether to perform mutagenesis as a one-step or two-step procedure depending on the number of gene fragments and mutations to be introduced. For the one-step procedure, the program designs primers flanked by BsaI or BbsI sites for direct cloning into the expression or cloning vector. For the two-steps procedure, primers are flanked by BbsI sites and the products are first cloned in an intermediate cloning vector before the final assembly in the expression vector. **B)** Schematic representation of the two-step procedure. Primers are designed carrying type IIS recognition sites (circle with S = BbsI and circle with A = BsaI) and 4 bp sequences for compatibility with the cloning vector (level 0, brown [5’] and yellow [3’]). Additionally, the primers at the 5’ and 3’ end include specific expression vector (level 2) overhangs (purple and bordeaux-red) and the “inner” primers carry gene-specific 4 bp overhangs for seamless reassembly of the fragments (white). Flanking the template binding part of the primer, randomization sites (blue with a red asterisk) are introduced. In the subsequent one-pot Golden Gate digestion-ligation reaction, BbsI binds to its recognition sequence and creates specific, pre-designed 4 bp overhangs (brown and beige) flanking the mutated PCR fragments, which are complementary to the exposed overhangs in the acceptor vector (cloning vector).

The final gene assembly step into the compatible *E. coli* expression vector (pAGM22082_CRed)—either directly from the obtained PCR fragments or via intermediate subcloning of individual fragments—is performed using an *E. coli* BL21(DE3) expression strain, enabling the direct assessment of the protein phenotypes of the respective clones. The DE3 genotype is crucial since it includes a T7 RNA polymerase gene thus enabling T7 promoter-dependent target gene expression. A BL21(DE3) strain harboring an additional pLysS plasmid was chosen in our studies. The pLysS plasmid is introduced to suppress basal expression of T7 RNA polymerase by constitutive expression of T7 lysozyme.^27^ This allows for tight control of gene expression and hence can improve the transformation efficiency in the case of potentially toxic proteins. In our studies, some mutagenesis gene targets seemed to have a strongly negative effect on cell growth (on LB agar plates) leading to overall low colony numbers as well as a substantial accumulation of unmodified expression plasmid. By using the pLysS harboring expression strain based on the possible color distinction mutagenesis libraries with high ratios of target colonies (>99 % of recombinant plasmid) could be obtained. Subsequent expression of the cloned gene requires induction of the T7 promoter using IPTG or lactose as an inductor. The screening of colonies containing recombinant constructs without inducing expression of the cloned genes requires a visual selection marker other than the widely spread blue/white LacZ color selection cassette, which requires basal lac promoter regulated expression. However, an IPTG/lactose induction would contradict the beneficial effect of the pLysS system. A novel pET28b based *E. coli* expression vector was therefore constructed, which carries an orange dye forming biosynthesis operon (termed CRed) under control of a constitutive promotor. The plasmid pAGM22082_CRed exhibits a T7 promoter for target gene expression and has been deposited at Addgene (plasmid #117225). The CRed selection cassette, which is released upon BsaI restriction digest consists of an eight kb canthaxanthin biosynthesis operon (Figure 2). This operon consists in total of seven genes—four genes from *Pantoea ananatis* (crtE, crtB, crtI and crtY) converting farnesyl pyrophosphate to β-carotene and an additional gene from *Agrobacterium aurantiacum* (crtW) to oxidize β-carotene to canthaxanthin.^28^ Also two *E. coli* derived genes are included to facilitate the synthesis of the isoprenoid precursor molecules. The more prominent orange color of canthaxanthin in comparison to β-carotene enables an easy detection on LB agar. The CRed marker operon within the pAGM22082 plasmid produces the carotenoid independently of an exogenous inducer and therefore allows orange/white selection in basically all common *E. coli* cloning as well as expression strains due to the intracellular accumulation of canthaxanthin (Figure S5).

**Figure 2.**
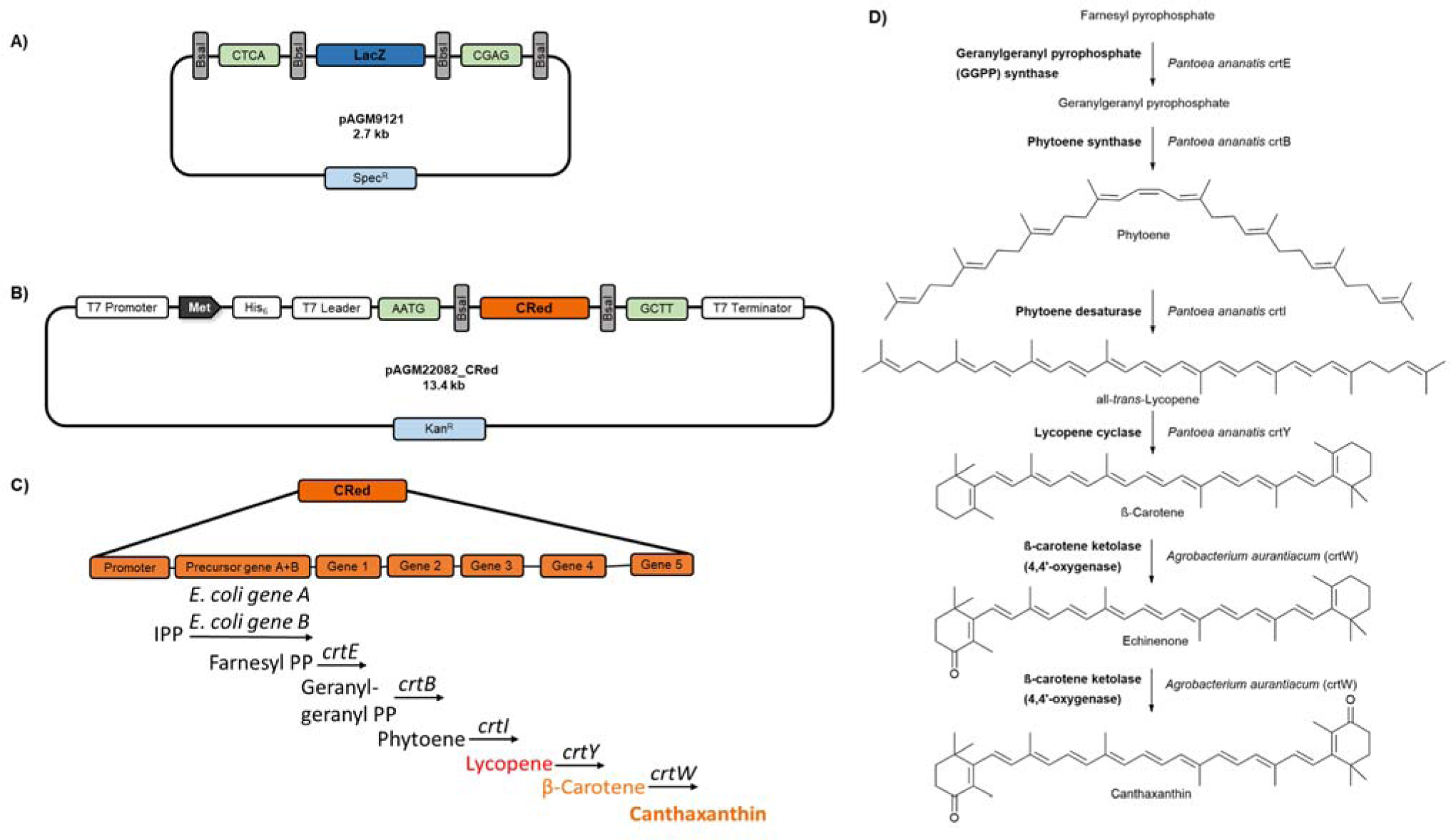
Overview of Golden Gate plasmids used in this study **A)** Schematic overview of the universal cloning plasmid (level 0) pAGM9121 which was used for all level 0 cloning approaches in this study. pAGM9121 harbors a LacZ selection marker which is flanked by internal BbsI sites specifying the 4 bp overhangs CTCA (5’) and CGAG (3’). Upon correct assembly of the insert, the BbsI sites are eliminated. The insert can be released in a subsequent restriction step based on the flanking BsaI sites (principle of cloning into pAGM22082_CRed plasmid). **B)** pAGM22082_CRed plasmid constructed in this study is a backbone derivative of the commercial pET28b plasmid harboring a canthaxanthin biosynthesis selection marker^28^ flanked by internal BsaI sites specifying the 4 bp overhangs AATG (start codon) and GCTT (post stop codon). Cloning of open reading frames into this plasmid is performed utilizing BsaI as restriction enzyme and occurs in frame with a 34 amino acid N-terminal linker, consisting of a hexahistidine-detection and purification tag as well as a T7 leader tag. Target gene expression is regulated by a T7 promoter. **C)** Overview of CRed selection marker. Consisting of a five-gene synthesis operon, reconstituting an artificial canthaxanthin biosynthesis pathway originating from farnesyl pyrophosphate as substrate and ultimately leading to the intracellular accumulation of the orange carotenoid canthaxanthin. Besides the synthesis unit, also two *E. coli* genes (*E. coli* gene A: isopentyl-diphosphate delta-isomerase and *E. coli* gene B: 1-deoxy-D-xylulose-5-phosphate synthase) are included for precursor synthesis within the operon. **D)** A detailed schematic description of the enzymatic reactions leading to canthaxanthin.

## Golden Mutagenesis software for in silico primer design and quick quality control (QQC)

Since the most challenging part of Golden Mutagenesis is the primer design considering suitable types IIs recognition sites, specified 4 bp overlaps, minimal melting temperature differences of corresponding primer pairs and specifying the mutation sites, an R package and a web tool was developed based on the programming language R. This web application is shown in Figure 3 and available at https://msbi.ipb-halle.de/GoldenMutagenesisWeb/. The source code of the package and an alternative interface based on Jupyter (run in binder^29^) notebooks are available at https://github.com/ipb-halle/GoldenMutagenesis/.

**Figure 3.**
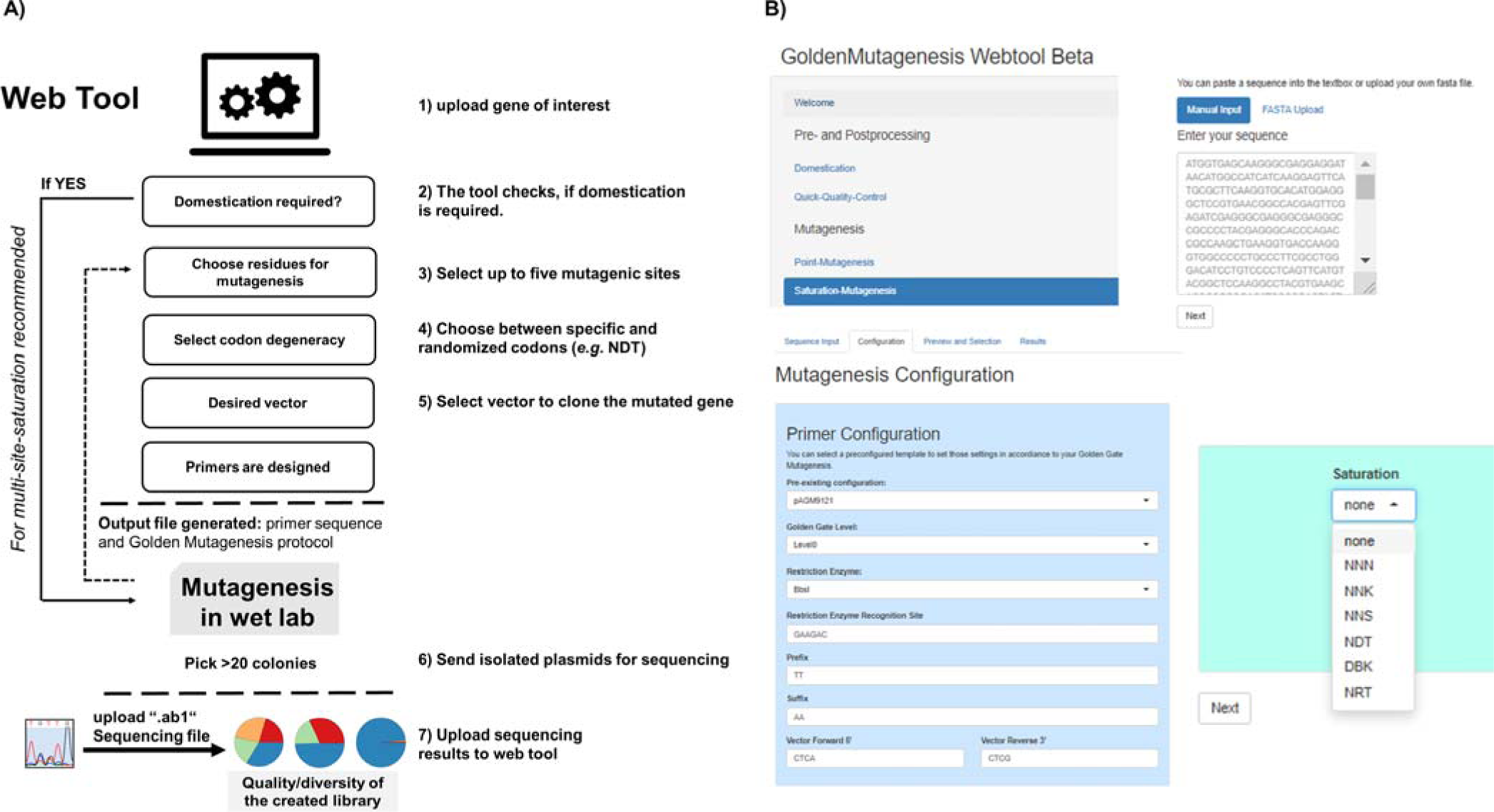
A)The workflow of the web tool for primer design and quick quality control. **B)** Screenshots of the web interface URL: https://msbi.ipb-halle.de/GoldenMutagenesisWeb/

The tool is implemented into the overall workflow as shown in Figure 3A. The user specifies the gene of interest and which residue(s) shall be modified and in which manner (specific vs saturation mutagenesis). Removal of internal type IIS restriction sites (termed domestication) within the gene sequence, which is highly recommended for efficient Golden Gate cloning, can be performed in parallel to the point or saturation mutagenesis step. The requirement for domestication is automatically indicated and suitable silent point mutations for the removal of internal cleavage sites are suggested and implemented by the tool. Primers which are generated by the tool for the purpose of point mutagenesis and domestication additionally take into account the *E. coli* codon usage to avoid the occurrence of rare codons upon mutagenesis which might hamper subsequent protein production. Besides *E. coli* codon usage also eukaryotic (*S. cerevisiae*) and plant (*A. thaliana*) codon usages are selectable options to streamline subsequent gene expression in those organisms using suitable Golden Gate plasmids. For multi-site-saturation mutagenesis, however, it is recommended to perform this domestication step in the “wet lab” before generating the library to provide a clean domesticated gene template prior to mutagenesis. This step will help to maximize the efficiency of the saturation mutagenesis experiment, as fewer PCR products are needed to be amplified and assembled at the same time. This domestication has only to be done once as the starting point of the entire mutagenesis endeavor.

Two different cloning options are available: a) Direct cloning of PCR fragments into the level 2 expression vector pAGM22082_CRed or level 0 cloning vector pAGM9121 (Figure 1A, left) or b) subcloning of individual gene fragments into the cloning vector pAGM9121 followed by an assembly of the subcloned fragments into the expression vector pAGM22082_CRed (Figure 1A, right). For the mutagenesis of a few residues, leading to 3 or fewer PCR fragments option a) is recommended since it is quicker and can be performed within one day.

The tool calculates and displays the full set of required primers, which carry the distinct type IIS restriction sites, mutagenic sites as well as suitable, matching four bp overhangs for specific sequential gene reassembly. First, the number of fragments is determined based on the distance of the selected mutation sites and a minimal fragment length. The tool decides whether a fragment should cover more than one mutation site, based on the user-specified parameters in the “Mutagenesis Configuration”. For option a) primers are flanked by BsaI (expression) or BbsI (cloning) sites, while for option b) primers are generally flanked by BbsI sites to enable cloning into pAGM9121. In the case of saturation mutagenesis approaches, the designed primers conceptually differ from the design of primers for targeted point mutagenesis, since randomization sites are not suitable for targeted reassembly and hence cannot be positioned within overhang regions, whereas in the case of point mutagenesis the altered genetic information is placed within the four bp overhang to facilitate correct gene fragment rejoining based on the altered information. Randomization sites (Figure 1B, blue with an asterisk) are introduced between the primer-template binding sequence (3’ direction) and the respective four base pair reassembly overhangs (5’direction). Matching primer pairs are automatically calculated and optimized to exhibit minimal differences in melting temperature with a default T_m_ value of 60 °C.^30,31^ A comprehensive text protocol enabling the user to easily perform all practical steps of the mutagenesis procedure is supplemented by the tool.

To access the quality of the generated library an analysis feature has been constructed and is available to easily assess the efficiency of the performed mutagenesis, a method that has previously been referred to as Quick Quality Control (QQC).^32^ The sequencing results in a “.ab1” file format containing chromatogram information can be uploaded to the web tool, which is based on sangerseq R package.^33^ The nucleobase distribution within the randomization sites can then be visualized as pie-charts (Figure 4).

**Figure 4.**
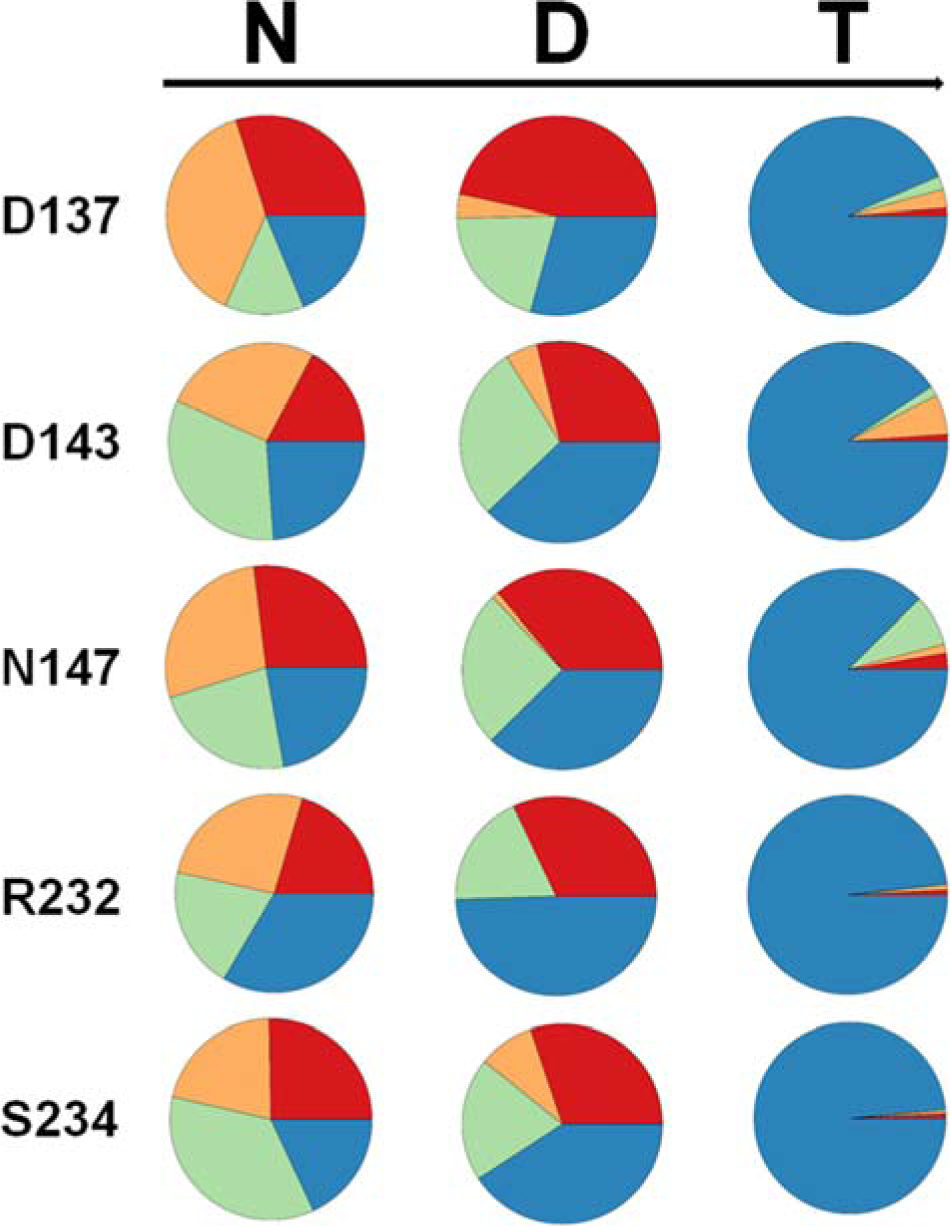
The created randomization libraries can be analyzed using the software tool. Base distribution with the example of a simultaneous NDT codon saturation at 5 residue positions (simultaneous NDT (N = A,T,C,G; D=A,T,G, T=T) randomization of 5 amino acid residues of the enzyme YfeX,). Color code: blue: thymine; orange: cytosine; green: guanine and red: adenine.

## Developed protocols

### Case I: Single site-directed mutagenesis and introduction of the target gene into the Golden Gate System

As a proof-of-concept for the functionality of the software, the widely used fluorescent protein mCherry^34^ was used as gene target, with the aim of introducing a single amino acid change (L69V) using a one nucleotide point mutation. The mCherry gene sequence contains an internal BbsI site which is recommended to be removed for Golden Gate cloning since the generally high efficiency of the digestion-ligation reaction is compromised in the case of the internal cleavage site, thus a second nucleotide exchange was envisioned. Therefore, the mCherry gene was split into three gene fragments using PCR. This workflow required a total amount of six primers flanked by BbsI recognition sites (Table S1). All primers were designed with the aid of the web tool, and the prior requirement to remove the internal BbsI restriction site was automatically detected by the program within the point mutagenesis workflow. New genetic information was automatically introduced at the terminal gene regions to provide a suitable type IIS recognition site as well as four bp overhang compatibility with the acceptor plasmid pAGM9121 (Figure S2). The performed cloning setup led to 29000 bacterial colonies with a 99:1 ratio of white to blue colonies (see Figure S3 for colony PCR). As mentioned above, it is recommended to perform an initial domestication step (only once required in the entire mutagenesis procedure) prior to the randomization of specific sites to obtain maximal efficiency of the Golden Gate cloning. Alternatively, a subcloning step of individual PCR fragments into a cloning vector and subsequent reassembly of the domesticated, mutated gene in a second step in the expression vector can be performed (Figure 1). However, if no randomization library but a specific, defined point mutation is required as above, this can preferably be done in one step.

### Case II: Multi-Site-Saturation Mutagenesis

Saturation mutagenesis was performed using the sequence of the *E. coli* derived YfeX^35,36^ gene as a template to introduce NDT codon degeneracy at either one or five positions simultaneously. The two strategies presented in Figure 1B were tested: direct cloning and assembly of the mutagenic PCR fragments into the final expression vector or subcloning each fragment into an intermediate cloning vector, followed by subsequent gene reassembly in the final expression vector.

Direct cloning into the expression vector introducing one NDT saturation site (two PCR fragments) and transformation of the ligation mixture in *E. coli* BL21(DE3) pLysS led to 860 white, recombined colonies and no orange colony, and a 96 % efficiency rate for correct YfeX reassembly—based on quick testing by colony PCR. In the case of the simultaneous saturation mutagenesis of five residues (three PCR fragments) 1400 white and 55 orange colonies were obtained. The assembly efficiency was 100 % based on 24 quick-tested colonies.

As a consequence of using the subcloning step of the PCR fragments followed by gene reassembly into the expression vector, substantially higher colony numbers were obtained in case of saturation mutagenesis at one position; 5000 white and twelve orange colonies could be obtained (ratio: 99.8:0.2). The efficiency for YfeX gene reassembly proved to be 100 % again. In the case of the saturation mutagenesis at 5 sites (three respective fragments), a higher colony number (1600 in total) and a white to orange colony ratio of 98:2 could be achieved, with a ratio of correct white colonies of 100 % as determined by analogous quick testing of 24 colonies (Figure S4).

In this single- as well as multi-site saturation mutagenesis approach of the YfeX gene nearly perfect efficiencies of correct Golden Gate gene reassembly could be achieved. Overall efficiency and fidelity of correct Golden Gate assembly in general, however, might be substantially depending on the nature of the created 4 bp overhang. Previous extensive studies on the fidelity of 4 bp overhangs revealed that certain AT-rich overhang like TAAA, TCAA, TGAA, TTAA are less efficiently joined by the employed T4 Ligase, thereby leading to the overall low fidelity of the Golden Gate assembly reaction.^23,37^

To access the base distribution within the randomization sites for this five saturation sites (D137, D143, N147, R232 and S234) approach, a QQC was performed and the sequencing results of a pooled library analyzed using the software tool (Figure 4). The expected NDT distribution pattern could be demonstrated at all 5 targeted positions, with a slight overrepresentation of thymine, which may be caused either by bias within the ordered primer mixtures or also possible due to altered template binding behavior within the primers containing a higher GC content due to more pronounced secondary structure occurrence, leading to less efficient annealing to the gene template. To prove that the overall genetic diversity of the library is not compromised by the diversity of the intermediate subcloning step, libraries obtained at the intermediate fragment subcloning step (level 0) were plated as well, leading to colony numbers of up to 58000 colonies and white to blue ratios of up to 99.7:0.3 (Table S2), thereby easily covering the required genetic diversity within the randomization sites.

## Conclusions

A Golden Gate cloning based protocol for the efficient execution of defined site-specific mutations within a gene of interest as well as for the generation of targeted randomization libraries is introduced. Efficient cloning protocols were established–including a novel Golden Gate compatible expression plasmid enabling T7 dependent expression in *E. coli*. The process of primer design was significantly simplified by the development of a freely available open source web tool that moreover facilitates the analysis of the randomization library quality by displaying nucleobase distributions within the envisioned randomization sites. The Golden Mutagenesis technique is in particular suited for streamlining iterative multiple site-saturation techniques like CASTing, ISM and SCSM/DCSM/TCSM protocols.

## Methods

### 1. General PCR Protocol

Reactions were carried out using standardized conditions. In a final reaction volume of 50 µl 100 ng of plasmid DNA (pET28a_mCherry or pCA24N_YfeX) were added as an amplification template. The pCA24N_YfeX plasmid was derived from the ASKA library.^38^ Final reaction mixtures consisted of 3 % (v/v) DMSO; 1x concentrated Phusion Green HF buffer (ThermoFisherScientific, Waltham, US); 0.5 units of Phusion High-Fidelity DNA polymerase (ThermoFisherScientific, Waltham, US); 200 µmol dNTP-Mix (ThermoFisherScientific, Waltham, US) and 200 nmol forward primer and reverse primer (stock solutions dissolved in ddH_2_0). PCR reactions were carried out by default under the following conditions: a) initial denaturation: 98 °C (60 s); b) cycling (35 passes in total): 95 °C (15 s), 60 °C (30 s) and 72 °C (90 s per kb amplification product) c) final elongation: 72 °C (10 min). Following PCR the samples were analyzed by agarose gel electrophoresis (7 V/cm; 45 min) (1 to 2.5 % (w/v) agarose (AppliChem, Darmstadt, DE) in TAE buffer), using a 1 kb DNA ladder (ThermoFisherScientific, Waltham, US) as standard for size determination. Therefore a volume of 5 µl out of the total PCR mixture was loaded onto the agarose gel for analysis. Double-stranded PCR products were visualized under UV light using a Genoplex Imager (VWR, Darmstadt, DE). After confirmation of occurrence and correct band sizes corresponding to PCR products (45 µl) were further purified. PCR products were recovered and purified using a NucleoSpin® Gel and PCR Clean-up Kit (Macherey-Nagel, Düren, DE). Concentrations of purified PCR products were determined by absorbance measurements at a wavelength of 260 nm using an infinite F200 pro device (TECAN, Grödig, AT).

### 2. Golden Gate cloning procedure

All Golden Gate reactions were performed in a total volume of 15 µl. The final reaction volume contained 1-fold concentrated T4 ligase buffer (Promega, Madison, US). Prepared reaction mixtures (ligase buffer, acceptor plasmid, insert(s)) was adjusted to 13.5 µl with ddH_2_O. In a final step, the corresponding enzymes were quickly added. First, a volume of 0.5 µl of the respective restriction enzyme BbsI (5 units; ThermoFisherScientific, Waltham, US) or BsaI-HF®v2 (10 units; New England Biolabs, Ipswich, US) and then 1 µl (1-3 units) of T4 ligase (Promega, Madison, US) was added. Golden Gate reactions were carried out by default under following conditions: a) Enzymatic restriction 37 °C (2 min) [40 passes]; b) Ligation 20 °C (5 min) [40 passes] and c) enzyme inactivation: 80 °C (20 min).

### 3. Procedure A: General Transformation

Following the Golden Gate reaction, the whole reaction volume of 15 µl was used to transform RbCl chemo-competent *E. coli* DH10B cells (aliquot of 50 µl; 7.1 × 10^7^ cfu/µg pUC19 DNA; ThermoFisherScientific, Waltham, US) by heat shock procedure (90 s at 42 °C).). After heat shock, cells were recovered in 500 µl LB medium for 1 h at 37 °C. Transformed cells were plated on LB agar plates (50 µg ml^−1^ X-Gal, 100 µg ml^−1^ spectinomycin, 150 µM IPTG). LB agar plates were then incubated overnight at 37 °C. Based on the presence of a LacZ cassette within the cloning site of pAGM9121 a color distinction (blue/white screening) between the unmodified pAGM9121 (blue colonies) and recombinant plasmid (white colonies) is possible. Colony numbers are given with two significant figures.^39^ For colony numbers > 50 the agar plates were divided into sections and the counted colonies multiplied with the number of sections. **Procedure B: Subcloning into cloning vector pAGM9121**. General transformation procedure as described in procedure A. After recovery, the transformation volume of approx. 550 µl was split into two fractions. A volume of 50 µl was used for plating on LB-agar plates (+ X-Gal,+ spectinomycin, + IPTG) and incubated overnight (37 °C) to evaluate the success of cloning on the following day (desired scenario: low percentage of blue colonies, high percentage and number of white colonies). The remaining volume was used to directly inoculate an overnight liquid culture of 4 ml TB medium (100 µg ml^−1^ spectinomycin) to preserve the prior created genetic diversity within the randomization sites of the saturation mutagenesis approach.

#### Procedure C: Transformation of expression vector pAGM22082_CRed

General transformation procedure as described procedure A but using RbCl chemo-competent BL21(DE3) pLysS (aliquot of 100 µl; 2.4 × 10^7^ cfu µg^−1^ pUC19 DNA, Merck Millipore, Darmstadt, DE). Transformed cells were plated (50 µl final volume) on LB agar plates (50 µg ml^−1^ kanamycin, 50 µg ml^−1^ chloramphenicol). LB agar plates were then incubated overnight at 37 °C. Based on the presence of an artificial bacterial operon responsible for canthaxanthin biosynthesis (termed CRed) within the cloning site of pAGM22082_CRed a color distinction between the unmodified pAGM22082_CRed (orange colonies; intracellular accumulation of canthaxanthin) and recombinant plasmid (white colonies; loss of operon) is possible.

### 4. Golden Gate cloning (cloning vector): point mutagenesis

General reaction conditions as described in section 2 using BbsI as restriction enzyme. The three PCR fragments for Golden Gate reassembly into the cloning vector (level 0) were prepared as described in section 1. The universal level 0 plasmid pAGM9121, which carries a LacZ selection marker, was used as acceptor plasmid for reassembly of mCherry in a final concentration of 20 fmol. Plasmid pAGM9121 can be obtained through Addgene (plasmid #51833). The three PCR fragments were added in equimolar amounts (20 fmol each). The transformation was performed according to procedure A.

### 5. Golden Gate cloning into cloning vector pAGM9121: individual fragment cloning for saturation mutagenesis

General reaction conditions as described in section 2 using BbsI as restriction enzyme. Individual PCR fragments for Golden Gate assembly into cloning vector were prepared as described in section 1. The plasmid pAGM9121 was used as acceptor plasmid in a final concentration of 20 fmol. PCR fragments were added in equimolar amounts relative to the plasmid (20 fmol each) and subsequent Golden Gate reactions were performed in parallel (*e.g.* two fragments in two parallel cloning approaches). The transformation was performed according to procedure B.

### 6. Direct Golden Gate cloning of PCR fragments into expression vector pAGM22082

General reaction conditions as described in section 2 using BsaI-HF®v2 as restriction enzyme. The PCR fragments for Golden Gate reassembly into level 2 were prepared as described in section 1. The level 2 expression plasmid pAGM22082_CRed was used as acceptor plasmid for reassembly of YfeX in a final concentration of 20 fmol. Plasmid pAGM22082_CRed can be obtained through Addgene (plasmid #117225). Respective PCR fragments were added in equimolar amount compared to the plasmid (20 fmol each). The transformation was performed according to procedure C.

### 7. Golden Gate cloning into expression vector pAGM22082 from cloning vector

General reaction conditions as described in section 2 using BsaI-HF®v2 as restriction enzyme. Starting from overnight grown liquid cultures of respective pAGM9121 constructs (prepared as described in section 5) plasmid mixtures were prepared using a NucleoSpin Plasmid Kit (Macherey-Nagel, Düren, DE). Following the preparation, the concentrations of purified plasmid DNA were determined by absorbance measurements at a wavelength of 260 nm using an infinite F200 pro device (TECAN, Grödig, AUT). The level 2 expression plasmid pAGM22082_CRed was used as acceptor plasmid for reassembly of YfeX in a final concentration of 20 fmol. Respective pAGM9121 donor plasmids were added in equimolar amounts relative to pAGM22082_CRed plasmid (20 fmol each). The transformation was performed according to procedure C.

### 8. Efficiency control of gene reassembly via colony PCR

After performing the Golden Gate cloning reactions the success rate of correct gene reassembly was determined by the means of colony PCR. PCR reactions were carried out using standardized conditions in a total reaction volume of 25 µl. Final reaction mixtures consisted of: 1x concentrated GoTaq® Green buffer (Promega, Madison, US); 0.7 units of GoTaq® DNA Polymerase (Promega, Madison, US); 150 µmol dNTP-Mix (ThermoFisherScientific, Waltham, US) and 150 nmol forward primer and reverse primer (stock solutions dissolved in ddH_2_0). As corresponding primer pairs, two options were chosen: a) forward and reverse primer of the corresponding acceptor plasmid or b) forward amplification primer of the insert and reverse primer of the acceptor plasmid. As amplification template, a clearly separated single *E. coli* colony was scratched off the LB agar plate using a sterile toothpick and transferred to the respective PCR tube.

PCR reactions were carried out by default using the following conditions: a) initial denaturation: 98 °C (120 s); b) cycling [35 passes]: 95 °C (15 s), 55 °C (30 s) and 72 °C (60 s per kb amplification product) c) final elongation: 72 °C (6 min). Following PCR the samples were analyzed by agarose gel electrophoresis (7 V/cm; 45 min) (1 % (w/v) agarose (AppliChem, Darmstadt, DE) in TAE buffer), using a 1 kb DNA Ladder (ThermoFisherScientific, Waltham, US) as the standard for size determination. Double-stranded PCR products were visualized under UV light using a Genoplex Imager (VWR, Darmstadt, DE).

### 9. Quick quality control (QQC) of randomization through sequencing

To assess the created genetic diversity in the prior mutagenesis step(s), a sequencing step was implemented.^32^ At least 150 white colonies were pooled and inoculated into 4 ml TB medium (50 µg ml^−1^ kanamycin, 50 µg ml^−1^ chloramphenicol). The liquid culture was incubated overnight (37 °C) for subsequent plasmid preparation. The plasmid mixtures were prepared as described in section 7. After determining the concentration, the elution volume was split into two tubes (approx. 18 µl each), the respective forward (pAGM22082_for) or reverse primer (pAGM22082_rev) at a concentration of 1 µM added and samples sent to Eurofins Genomics (Ebersberg, DE) for sequencing. Sequencing results were received in a .ab1 file format the following day.

## Data availability

Vectors and inserts: The universal level 0 Golden Gate cloning vector pAGM9121 (plasmid #51833) and the newly designed pET-based level 2 *E. coli* expression vector pAGM22082_CRed (#117225) can be obtained through the non-profit plasmid depository Addgene (www.addgene.org). Detailed information on the usage of the R Package can be found in the R vignettes and documentation files provided as supplemental material. They cover three examples (point mutagenesis, site-saturation mutagenesis and graphical illustration of quick quality control (QQC)) including the created data. The web application is available at https://msbi.ipb-halle.de/GoldenMutagenesisWeb/. The source code and an alternative interface based on Jupyter notebooks are available at https://github.com/ipb-halle/GoldenMutagenesis/ under the LGPL Open Source license.

## Supporting information

Experimental details

Computer Tool example Multi-Site Directed Mutagenesis 1

Computer Tool example Multi-Site Directed Mutagenesis 2

Computer Tool example Multi-Site Directed Mutagenesis 3

Computer Tool example point mutagenesis and domestication

Computer Tool example Quick Quality Control

## Acknowledgement

M.J.W thanks the BMBF ("Biotechnologie 2020+ Strukturvorhaben: Leibniz Research Cluster“, 031A360B) for generous funding. P.P. thanks the Landesgraduiertenförderung Sachsen-Anhalt for a PhD scholarship. C.U thanks the Leibniz SAW funding. The authors are moreover grateful to Dr. Martin Dippe for providing the mCherry gene as well as Jonathon Hill who developed the PolyPeakParser Package and provided helpful tips for accessing the sequencing data via R.

## Author Contribution Statement

PP and SM planned and performed the cloning setup as well as plasmid construction. RG supported practical work and data acquisition. CU and SN designed the strategy for the *in silico* primer design tool and CU scripted it. CU and PP developed the *in silico* sequencing analysis for Quick Quality Control (QCC). MJW and PP designed the study. MJW, PP and SM wrote the manuscript, which was proofread and approved by all authors.

## Additional information

### Supplementary information

Experimental details (File Supplemental info), Open Source Computer Tool examples for Multi-Site Directed Mutagenesis (File: Vignette Multi SD, Vignette Multi SD2, Vignette SD3), point mutagenesis and domestication (File: Vignette Point_Mutagenesis and Domestication), and Quick Quality Control (File: Vignette QQC).

## Tables

**Table 1.**
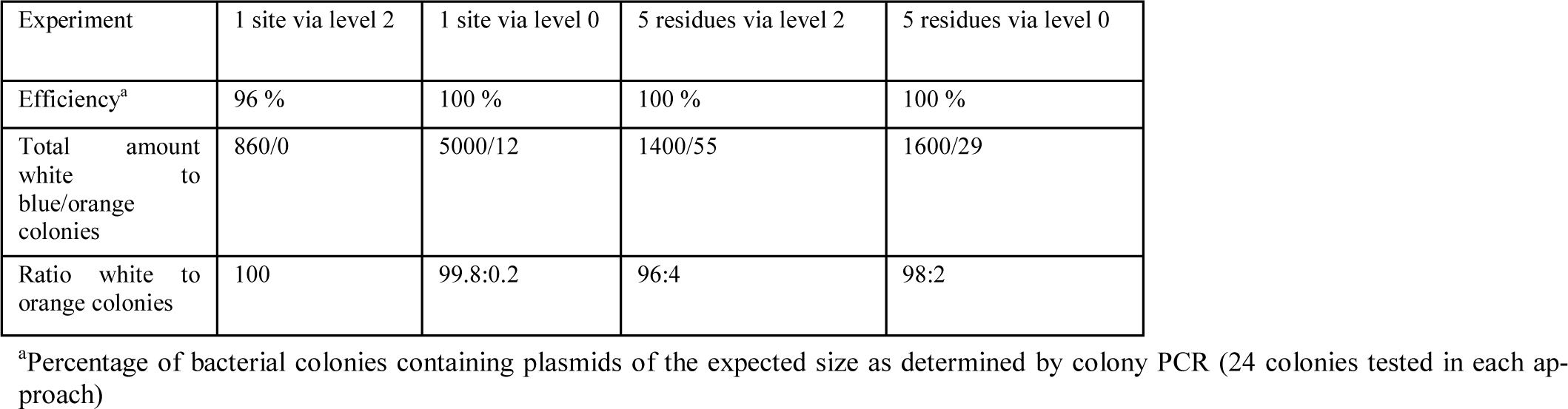
Overview of cloning results for NDT saturation mutagenesis approaches.

| Experiment | 1 site via level 2 | 1 site via level 0 | 5 residues via level 2 | 5 residues via level 0 |
| --- | --- | --- | --- | --- |
| Efficiency<sup>a</sup> | 96 % | 100 % | 100 % | 100 % |
| Total amount white to blue/orange colonies | 860/0 | 5000/12 | 1400/55 | 1600/29 |
| Ratio white to orange colonies | 100 | 99.8:0.2 | 96:4 | 98:2 |

<sup>a</sup>Percentage of bacterial colonies containing plasmids of the expected size as determined by colony PCR (24 colonies tested in each approach)

## References

1 Reetz, M. T., Wang, L. W. & Bocola, M. Directed evolution of enantioselective enzymes: iterative cycles of CASTing for probing protein-sequence space. Angew Chem Int Ed 45, 1236–1241, doi:10.1002/anie.200502746 (2006).

2 Reetz, M. T. et al. Expanding the substrate scope of enzymes: combining mutations obtained by CASTing. Chemistry 12, 6031–6038, doi:10.1002/chem.200600459 (2006).

3 Reetz, M. T. & Carballeira, J. D. Iterative saturation mutagenesis (ISM) for rapid directed evolution of functional enzymes. Nat Protoc 2, 891–903, doi:10.1038/nprot.2007.72 (2007).

4 Acevedo-Rocha, C. G., Hoebenreich, S. & Reetz, M. T. Iterative saturation mutagenesis: a powerful approach to engineer proteins by systematically simulating Darwinian evolution. Methods Mol Biol 1179, 103–128, doi:10.1007/978-1-4939-1053-3_7 (2014).

5 Chiang, L. W., Kovari, I. & Howe, M. M. Mutagenic oligonucleotide-directed PCR amplification (Mod-PCR): an efficient method for generating random base substitution mutations in a DNA sequence element. PCR Methods Appl 2, 210–217 (1993).

6 Kegler-Ebo, D. M., Docktor, C. M. & DiMaio, D. Codon cassette mutagenesis: a general method to insert or replace individual codons by using universal mutagenic cassettes. Nucleic Acids Res 22, 1593–1599 (1994).

7 Ho, S. N., Hunt, H. D., Horton, R. M., Pullen, J. K. & Pease, L. R. Site-directed mutagenesis by overlap extension using the polymerase chain reaction. Gene 77, 51–59 (1989).

8 Tyagi, R., Lai, R. & Duggleby, R. G. A new approach to ‘megaprimer’ polymerase chain reaction mutagenesis without an intermediate gel purification step. BMC Biotechnol 4, 2, doi:10.1186/1472-6750-4-2 (2004).

9 Liu, H. & Naismith, J. H. An efficient one-step site-directed deletion, insertion, single and multiple-site plasmid mutagenesis protocol. BMC Biotechnol 8, 91, doi:10.1186/1472-6750-8-91 (2008).

10 Dennig, A., Shivange, A. V., Marienhagen, J. & Schwaneberg, U. OmniChange: the sequence independent method for simultaneous site-saturation of five codons. PLoS One 6, e26222, doi:10.1371/journal.pone.0026222 (2011).

11 Cozens, C. & Pinheiro, V. B. Darwin Assembly: fast, efficient, multi-site bespoke mutagenesis. Nucleic Acids Res 46, e51, doi:10.1093/nar/gky067 (2018).

12 Herman, A. & Tawfik, D. S. Incorporating Synthetic Oligonucleotides via Gene Reassembly (ISOR): a versatile tool for generating targeted libraries. Protein Eng Des Sel 20, 219–226, doi:10.1093/protein/gzm014 (2007).

13 Vazquez-Vilar, M. et al. Software-assisted stacking of gene modules using GoldenBraid 2.0 DNA-assembly framework. Methods Mol Biol 1284, 399–420, doi:10.1007/978-1-4939-2444-8_20 (2015).

14 Swainston, N., Currin, A., Kell, D. B. & Day, P. J. GeneGenie: optimized oligomer design for directed evolution. Nucleic Acids Res 42, W395–W400, doi:10.1093/nar/gku336 (2014).

15 Li, A. et al. Beating Bias in the Directed Evolution of Proteins: Combining High-Fidelity on-Chip Solid-Phase Gene Synthesis with Efficient Gene Assembly for Combinatorial Library Construction. ChemBioChem 19, 221–228, doi:10.1002/cbic.201700540 (2018).

16 Engler, C., Kandzia, R. & Marillonnet, S. A one pot, one step, precision cloning method with high throughput capability. PLoS One 3, e3647, doi:10.1371/journal.pone.0003647 (2008).

17 Engler, C., Gruetzner, R., Kandzia, R. & Marillonnet, S. Golden Gate Shuffling: A One-Pot DNA Shuffling Method Based on Type IIs Restriction Enzymes. PLOS ONE 4, e5553, doi:10.1371/journal.pone.0005553 (2009).

18 Werner, S., Engler, C., Weber, E., Gruetzner, R. & Marillonnet, S. Fast track assembly of multigene constructs using Golden Gate cloning and the MoClo system. Bioeng Bugs 3, 38–43, doi:10.1371/journal.pone.0016765 (2012).

19 Sarrion-Perdigones, A. et al. GoldenBraid: an iterative cloning system for standardized assembly of reusable genetic modules. PLoS One 6, e21622, doi:10.1371/journal.pone.0021622 (2011).

20 Andreou, A. I. & Nakayama, N. Mobius Assembly: A versatile Golden-Gate framework towards universal DNA assembly. PLOS ONE 13, e0189892, doi:10.1371/journal.pone.0189892 (2018).

21 Pollak, B. et al. Loop assembly: a simple and open system for recursive fabrication of DNA circuits. New Phytol 222, 628–640, doi:10.1111/nph.15625 (2019).

22 van Dolleweerd, C. J. et al. MIDAS: A Modular DNA Assembly System for Synthetic Biology. ACS Synth Biol 7, 1018–1029, doi:10.1021/acssynbio.7b00363 (2018).

23 Potapov, V. et al. Comprehensive Profiling of Four Base Overhang Ligation Fidelity by T4 DNA Ligase and Application to DNA Assembly. ACS Synth Biol 7, 2665–2674, doi:10.1021/acssynbio.8b00333 (2018).

24 Quaglia, D., Ebert, M. C., Mugford, P. F. & Pelletier, J. N. Enzyme engineering: A synthetic biology approach for more effective library generation and automated high-throughput screening. PLoS One 12, e0171741, doi:10.1371/journal.pone.0171741 (2017).

25 Yan, P., Gao, X., Shen, W., Zhou, P. & Duan, J. Parallel assembly for multiple site-directed mutagenesis of plasmids. Anal Biochem 430, 65–67, doi:10.1016/j.ab.2012.07.029 (2012).

26 Engler, C. et al. A golden gate modular cloning toolbox for plants. ACS Synth Biol 3, 839–843, doi:10.1021/sb4001504 (2014).

27 Studier, F. W. Use of bacteriophage T7 lysozyme to improve an inducible T7 expression system. J Mol Biol 219, 37–44, doi:10.1016/0022-2836(91)90855-Z (1991).

28 Weber, E., Engler, C., Gruetzner, R., Werner, S. & Marillonnet, S. A Modular Cloning System for Standardized Assembly of Multigene Constructs. PLOS ONE 6, e16765, doi:10.1371/journal.pone.0016765 (2011).

29 Bussonnier, M. et al. Binder 2.0 - Reproducible, interactive, sharable environments for science at scale. Proc Python Scien, 113–120, doi:10.25080/Majora-4af1f417-011 (2018).

30 Breslauer, K. J., Frank, R., Blocker, H. & Marky, L. A. Predicting DNA duplex stability from the base sequence. Proc Natl Acad Sci 83, 3746–3750, doi:10.1073/pnas.83.11.3746 (1986).

31 Kibbe, W. A. OligoCalc: an online oligonucleotide properties calculator. Nucleic Acids Res 35, W43–W46, doi:10.1093/nar/gkm234 (2007).

32 Acevedo-Rocha, C. G., Reetz, M. T. & Nov, Y. Economical analysis of saturation mutagenesis experiments. Sci Rep 5, 10654, doi:10.1038/srep10654 (2015).

33 Hill, J. T. et al. Poly peak parser: Method and software for identification of unknown indels using sanger sequencing of polymerase chain reaction products. Dev Dyn 243, 1632–1636, doi:10.1002/dvdy.24183 (2014).

34 Shaner, N. C. et al. Improved monomeric red, orange and yellow fluorescent proteins derived from Discosoma sp. red fluorescent protein. Nat Biotechnol 22, 1567–1572, doi:10.1038/nbt1037 (2004).

35 Weissenborn, M. J. et al. Enzyme-Catalyzed Carbonyl Olefination by the E. coli Protein YfeX in the Absence of Phosphines. ChemCatChem 8, 1636–1640, doi:doi:10.1002/cctc.201600227 (2016).

36 Hock, K. J. et al. Tryptamine Synthesis by Iron Porphyrin Catalyzed C-H Functionalization of Indoles with Diazoacetonitrile. Angew Chem Int Ed 58, 3630–3634, doi:10.1002/anie.201813631 (2019).

37 HamediRad, M., Weisberg, S., Chao, R., Lian, J. & Zhao, H. Highly Efficient Single-Pot Scarless Golden Gate Assembly. ACS Synth Biol, doi:10.1021/acssynbio.8b00480 (2019).

38 Kitagawa, M. et al. Complete set of ORF clones of Escherichia coli ASKA library (a complete set of E. coli K-12 ORF archive): unique resources for biological research. DNA Res 12, 291–299, doi:10.1093/dnares/dsi012 (2005).

39 Sutton, S. Counting colonies. PMF Newsl 12, 2–5 (2006).

