## Supplementary material for "Golden Mutagenesis: An efficient multi-site-saturation mutagenesis approach by Golden Gate cloning with automated primer design": Experimental details

#### Table SI- Primer list

Table S1. Overview of oligonucleotides used in this study. All Oligonucleotides were purchased in the purification grade “desalted” at Eurofins Genomics (Ebersberg, DE). Oligonucleotide sequences contain space characters to highlight general features of oligonucleotides: TT − type IIS restriction site – spacer nucleotides – post-cleavage 4 bp overhangs − template binding sequence (5´ to 3´ direction). Oligonucleotides, which were designed with the aid of the open source tool, are marked in italic and labeled with an asterisk (computer designed primers).

| Name | Sequence (5´ → 3´) | Purpose |
|---|---|---|
| *mcherry_I_for** | TT GAAGAC AA CTCA ATGGTGAGCAAGGGCGAGGAGG | Cloning of mCherry into pAGM9121 |
| *mcherry_I_rev** | TT GAAGAC AA CACG ATGTCCCAGGCGAAGGGCAGGG | Cloning of mCherry into pAGM9121 |
| *mcherry_II_for** | TT GAAGAC AA CGTG TCCCCTCAGTTCATGTACGGCTCC | Cloning of mCherry into pAGM9121 |
| *mcherry_II_rev** | TT GAAGAC AA TTTC TGCATTACGGGGCCGTCGGA | Cloning of mCherry into pAGM9121 |
| *mcherry_III_for** | TT GAAGAC AA GAAA AAGACGATGGGCTGGGAGGCC | Cloning of mCherry into pAGM9121 |
| *mcherry_III_rev** | TT GAAGAC AA CTCG TCACTCGAGTGCGGCCGC | Cloning of mCherry into pAGM9121 |
| *Yfex_Level 2_I_for** | TT GGTCTC A AATG TCTCAGGTTCAGAGTGGCATTTTGCC | Cloning of Yfex into pAGM22082_cRed (via level 2) |
| Yfex_Level 2_III_rev | TT GGTCTC T ACCC GCCGGGTTTTCCGTACCGTCAA | Cloning of Yfex into pAGM22082_cRed (via level 2) |
| Yfex_Level 2_IV_for | TT GGTCTC A GGGT NDTGAGACGCGTCGCGAAGTGGCGGTTATC | Cloning of Yfex into pAGM22082_cRed (via level 2) |
| *Yfex_Level 2_VI_rev** | TT GGTCTC T AAGC TTACAGCGCCATCAACTTGTCCAGC | Cloning of Yfex into pAGM22082_cRed (via level 2) |
| *Yfex_2_active site_I_rev** | TT GGTCTC T ACAA AGCCGCTCAGAHNACGCTCTTCAACCCAACGGAAGC | Cloning of Yfex into pAGM22082_cRed (via level 2) |

| | | |
|---|---|---|
| *Yfex_2_active site_II_for** | TT GGTCTC A TTGT TNDTGGTACGGAANDTCCGGCGGGTGAAGAGACGC | Cloning of Yfex into pAGM22082_cRed (via level 2) |
| *Yfex_2_active site_II_rev** | TT GGTCTC T AACA ATCTTCAGCCCTTTGCCATCTTCTTT | Cloning of Yfex into pAGM22082_cRed (via level 2) |
| *Yfex_2_active site_III_for** | TT GGTCTC A TGTT NDTCAGNDTCTGCCGTACGGCACTGCCAG | Cloning of Yfex into pAGM22082_cRed (via level 2) |
| *Yfex_Level 0_I_for** | TT GAAGAC AA CTCA AATG TCTCAGGTTCAGAGTGGCATTTTGCC | Cloning of Yfex into pAGM22082_cRed (via level 0) |
| Yfex_Level 0_III_rev | TT GAAGAC AA CTCG ACCC GCCGGGTTTTCCGTACCGTCAACAAA | Cloning of Yfex into pAGM22082_cRed (via level 0) |
| Yfex_Level 0_IV_for | TT GAAGAC AA CTCA GGGT NDTGAGACGCGTCGCGAAGTGGCGGTTATC | Cloning of Yfex into pAGM22082_cRed (via level 0) |
| *Yfex_Level 0_VI_rev** | TT GAAGAC AA CTCG AAGC TTACAGCGCCATCAACTTGTCCAGC | Cloning of Yfex into pAGM22082_cRed (via level 0) |
| *Yfex_0_active site_I_rev** | TT GAAGAC AA CTCG ACAA AGCCGCTCAGAHNACGCTCTTCAACCCAACGGAAGC | Cloning of Yfex into pAGM22082_cRed (via level 0) |
| *Yfex_0_active site_II_for** | TT GAAGAC AA CTCA TTGT TNDTGGTACGGAANDTCCGGCGGGTGAAGAGACGC | Cloning of Yfex into pAGM22082_cRed (via level 0) |
| *Yfex_0_active site_II_rev** | TT GAAGAC AA CTCG AACA ATCTTCAGCCCTTTGCCATCTTCTTT | Cloning of Yfex into pAGM22082_cRed (via level 0) |
| *Yfex_0_active site_III_for** | TT GAAGAC AA CTCA TGTTNDTCAGNDTCTGCCGTACGGCACTGCCAG | Cloning of Yfex into pAGM22082_cRed (via level 0) |
| pAGM22082_for | GAAGGAGATATACCATGGGCAGCAG | sequencing/colony PCR |
| pAGM22082_rev | CGTTTAGAGGCCCCAAGGGGTTATG | sequencing/colony PCR |
| pAGM9121_for | CCTGTCGGGTTTCGCCACCT | sequencing/colony PCR |
| pAGM9121_rev | GCCGTTACCACCGCTGCGTT | sequencing/colony PCR |

### Figures

#### Figure S1- Utilized type IIS restriction enzymes

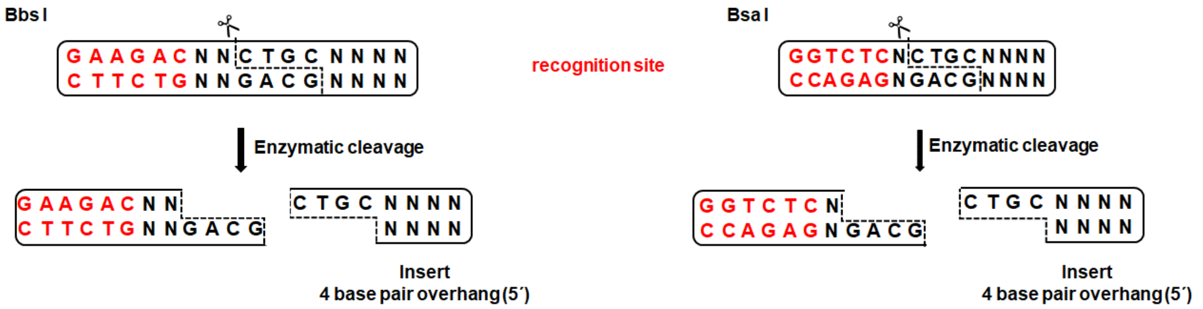


Figure S1. Recognition and cleavage pattern of type IIS restriction enzymes BbsI (from *Bacillus laterosporus*) and BsaI (from *Bacillus stearothermophilus*) used in this study. Based on the unique feature of this enzyme subclass to cleave double-stranded DNA fragments outside of their recognition site the original recognition sites are not included in the cleavage product (see above: insert). Created 4 bp can be specified through primer design to facilitate subsequent joining of neighbouring fragments in a gene reassembly reaction. Cloning, therefore, occurs in a scarless manner and preserves intact open reading frames.

**Figure S2- Cloning scheme point mutagenesis: mCherry**

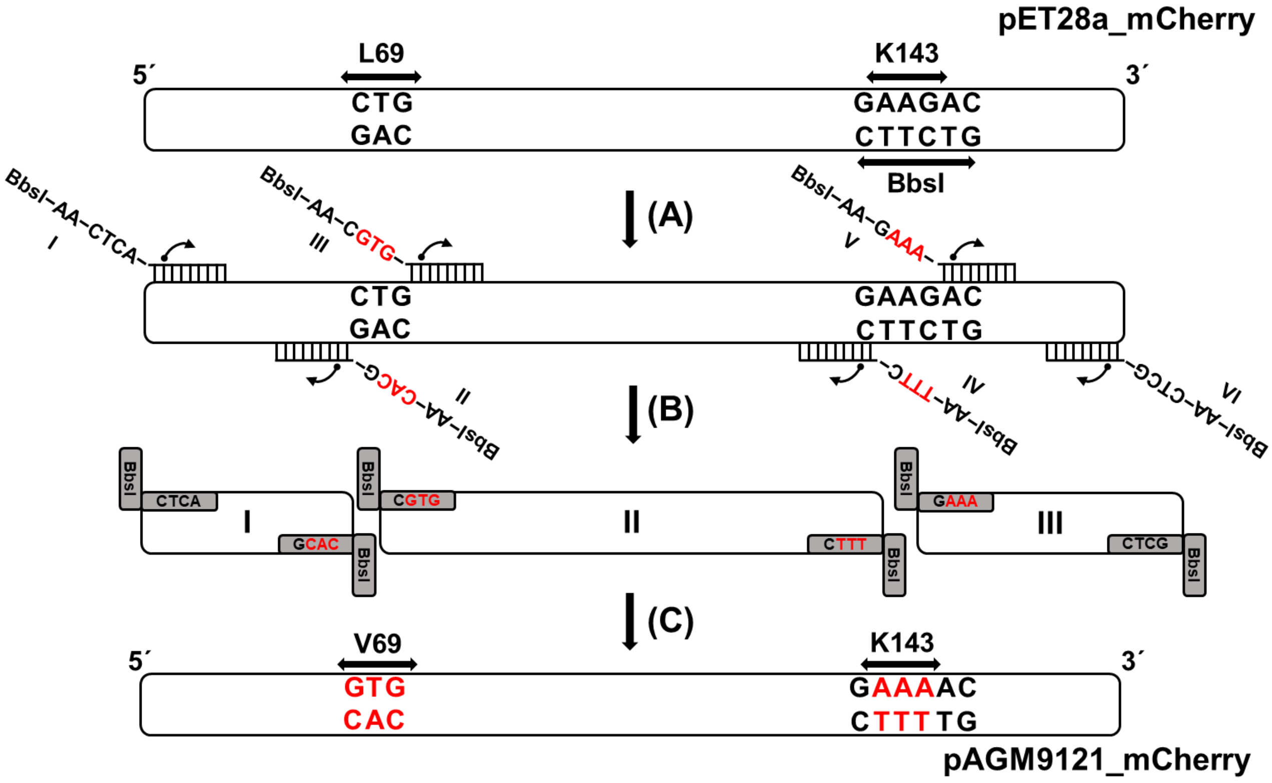


Figure S2. Cloning scheme of mCherry domestication (removal of internal type IIS restriction site) and point mutagenesis (L69V exchange) cloning into pAGM9121 (see S2 (D)). The template mCherry was cloned into a pET28a backbone and exhibiting one internal BbsI cleavage site, spanning amongst others the residue K143. An amino acid exchange of Leucine 69 (CTG) to Valine (GTG) was envisioned as well. Oligonucleotides were designed to contain terminal BbsI cleavage sites, followed by two spacer nucleotides (AA) (step A). New genetic information (displayed in red) is specified and introduced through primer design and leads to the creation of 4 bp overhangs in the course of the Golden Gate digestion ligation reaction (Figure S1). Primer as listed in table SI (I to VI: mcherry_I_for […] mcherry_III_rev). After PCR (step B) three fragments are generated with flanking BbsI recognition sites. In a subsequent Golden Gate reaction the PCR fragments are reassembled into the level 0 acceptor plasmid pAGM9121 (step C).

**Figure S3- Testing of correct assembly via colony PCR (mCherry)**

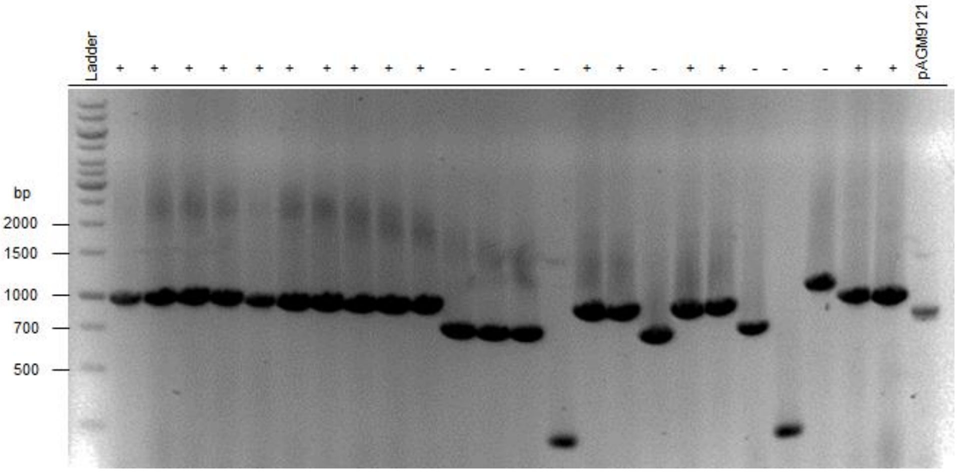


Figure S3. Analysis of correct mCherry reassembly via colony PCR using pAGM9121 plasmid primer. The expected insert size of the reassembled mCherry gene is approx. 1000 bp. In the case of 16 out of total 24 colonies the expected insert was observed (marked with a +). In the case of many negative colonies (marked with a -), slightly smaller fragments sizes (~200 bp less) were detected suggesting the correct insertion of 2 out of 3 PCR fragments and the occurrence of a competing insertion of a primer dimer fragment in one case into pAGM9121. pAGM9121 harboring the LacZ selection marker was also tested as a negative control, exhibiting a lower fragment size (approx. 850 bp). The picture was acquired using a Genoplex Imager (VWR, Darmstadt, DE) with a 300 ms exposure time. To enhance the visibility, colors were inverted and contrast/brightness adapted to an optimal value. A caption was added afterwards. The gel image was not cropped nor cut.

**Figure S4- Testing of correct assembly via colony PCR (YfeX)**

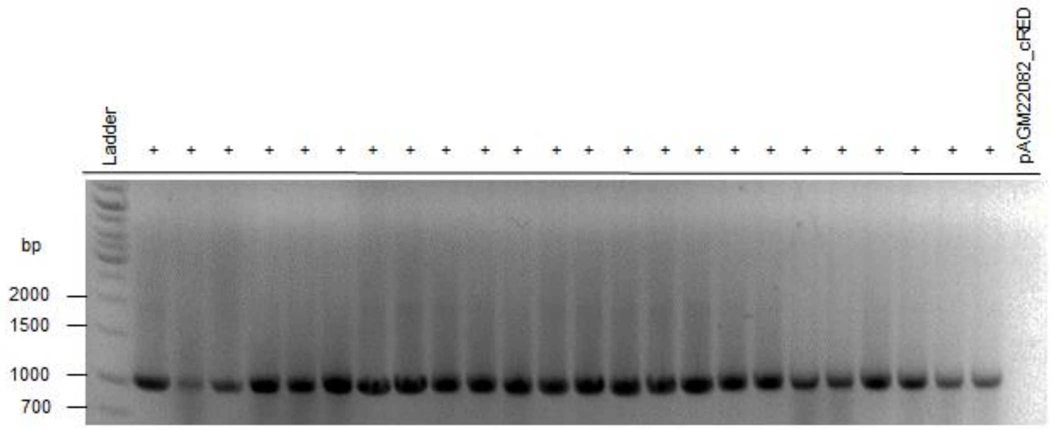


Figure S4. Analysis of correct YfeX reassembly in expression vector after a subcloning step of respective gene fragments into a cloning vector. Determination via colony PCR using Yfex_Level 0_I_for and pAGM22082_rev plasmid primer. Five amino acid residues (D137, D143, N147, R232, S234) within the active site were chosen for randomization. The expected insert size of the correctly reassembled YfeX gene is approx. 1000 bp. In the case of 24 out of total 24 colonies the expected insert was observed (marked with a +). pAGM22082_CRed was also tested as a negative control, leading to no observable amplification product. The picture was acquired using a Genoplex Imager (VWR, Darmstadt, DE) with a 300 ms exposure time. To enhance the visibility, colors were inverted and contrast/brightness adapted to an optimal value. A caption was added afterwards. The gel image was not cropped nor cut.

**Figure S5- LB-agar plate illustrating the canthaxanthin-based color formation**

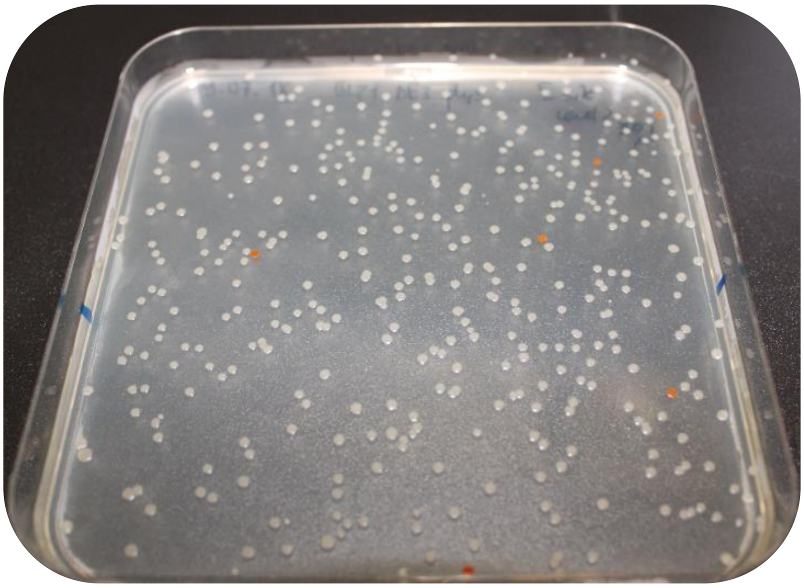

Figure S5. Exemplary picture of performed orange/white color selection on an LB-agar plate. Based on the accumulation of canthaxanthin the original plasmid (pAGM22082_CRed) can be clearly distinguished from recombined, white colonies.

**Table S2- level 0 gene parts plasmid overview**

Table S2. Overview of colony amount and blue/white distribution pattern for the created level 0 gene subcloning plasmids. Due to the creation of degenerated plasmid libraries trough PCR a minimal colony amount is necessary to preserve the desired degeneracy (e.g. one NDT site: 12 different combinations possible) within the library. In the following Table the colony amounts and blue/white ratios for all created level 0 subcloning plasmids is displayed.

| **Experiment** | **1 site via level 0** | | **active site via level 0** | | |
|---|---|---|---|---|---|
| **Level 0 plasmid** | YfeX_Part I | YfeX_Part II | YfeX_Part I | YfeX_Part II | YfeX_Part III |
| **Total amount white/blue** | 58000/350 | 30000/400 | 58000/800 | 54000/750 | 19000/44 |
| **Ratio white to blue colonies** | 99.4:0.6 | 98.7:1.3 | 98.6:1.4 | 98.6:1.4 | 99.7:0.3 |

### Gene sequences

#### mCherry (in pET28a)

ATGGTGAGCAAGGGCGAGGAGGATAACATGGCCATCATCAAGGAGTTCATGCGCTTCAAGGTGC
ACATGGAGGGCTCCGTGAACGGCCACGAGTTCGAGATCGAGGGCGAGGGCGAGGGCCGCCCC
TACGAGGGCACCCAGACCGCCAAGCTGAAGGTGACCAAGGGTGGCCCCCTGCCCTTCGCCTGG
GACATCCTGTCCCCTCAGTTCATGTACGGCTCCAAGGCCTACGTGAAGCACCCCGCCGACATCC
CCGACTACTTGAAGCTGTCCTTCCCCGAGGGCTTCAAGTGGGAGCGCGTGATGAACTTCGAGGA
CGGCGGCGTGGTGACCGTGACCCAGGACTCCTCCCTGCAGGACGGCGAGTTCATCTACAAGGT
GAAGCTGCGCGGCACCAACTTCCCCTCCGACGGCCCCGTAATGCAGAAGAAGACGATGGGCTG
GGAGGCCTCCTCCGAGCGGATGTACCCCGAGGACGGCGCCCTGAAGGGCGAGATCAAGCAGA
GGCTGAAGCTGAAGGACGGCGGCCACTACGACGCTGAGGTCAAGACCACCTACAAGGCCAAGA
AGCCCGTGCAGCTGCCCGGCGCCTACAACGTCAACATCAAGTTGGACATCACCTCCCACAACGA
GGACTACACCATCGTGGAACAGTACGAACGCGCCGAGGGCCGCCACTCCACCGGCGGCATGGA
CGAGCTGTACAAGGTCGACAAGCTTGCGGCCGCACTCGAGTGA

#### YfeX (in pCA24N)

ATGTCTCAGGTTCAGAGTGGCATTTTGCCAGAACATTGCCGCGCGGCGATTTGGATCGAAGCCA
ACGTGAAAGGGGAAGTTGACGCCCTGCGTGCGGCCAGTAAAACATTTGCCGACAAACTGGCAAC
TTTTGAAGCGAAATTCCCGGACGCGCATCTTGGTGCGGTGGTTGCCTTTGGTAACAACACCTGGC
GCGCTCTGAGCGGCGGCGTTGGGGCAGAAGAGCTGAAAGATTTTCCGGGCTACGGTAAAGGCC
TTGCGCCGACGACCCAGTTCGATGTGTTGATCCACATTCTTTCTCTGCGTCACGACGTAAACTTC
TCTGTCGCCCAGGCGGCGATGGAAGCCTTTGGTGACTGCATTGAAGTGAAAGAAGAGATCCACG
GCTTCCGTTGGGTTGAAGAGCGTGACCTGAGCGGCTTTGTTGACGGTACGGAAAACCCGGCGG
GTGAAGAGACGCGTCGCGAAGTGGCGGTTATCAAAGACGGCGTGGATGCGGGCGGCAGCTATG
TGTTTGTCCAGCGTTGGGAACACAACCTGAAGCAGCTCAACCGGATGAGCGTTCACGATCAGGA
GATGGTGATCGGGCGCACCAAAGAGGCCAACGAAGAGATCGACGGCGACGAACGTCCGGAAAC
CTCTCACCTCACCCGCGTTGATCTGAAAGAAGATGGCAAAGGGCTGAAGATTGTTCGCCAGAGC
CTGCCGTACGGCACTGCCAGTGGCACTCACGGTCTGTACTTCTGCGCCTACTGCGCGCGTCTGC
ATAACATTGAGCAGCAACTGCTGAGCATGTTTGGCGATACCGATGGTAAGCGTGATGCGATGTTG
CGTTTCACCAAACCGGTAACCGGCGGCTATTATTTCGCACCGTCGCTGGACAAGTTGATGGCGC
TGTAA

### Literature

(1) Kitagawa, M., Ara, T., Arifuzzaman, M., Ioka-Nakamichi, T., Inamoto, E., Toyonaga, H., and Mori, H. (2005) Complete set of ORF clones of Escherichia coli ASKA library (a complete set of E. coli K-12 ORF archive): unique resources for biological research, *DNA Res 12*, 291-299.
