## Supplementary material for "Golden Mutagenesis: An efficient multi-site-saturation mutagenesis approach by Golden Gate cloning with automated primer design": Computer Tool example Multi-Site Directed Mutagenesis 1

Multiple Site Saturation Mutagenesis

2018-10-10

### Multiple Site Saturation Mutagenesis – Active site

You can find the lastest version of this file at <https://github.com/ipb-halle/GoldenMutagenesis/blob/master/vignettes/MSD.md>

#### Experimental Workflow

##### Target sequence

open reading frame YfeX in pCA24N (Chloramphenicol resistance)

##### Clone into

seperate gene fragments into pAGM9121 first then reassemble into pAGM22082_cRed

##### Genomic sequence YfeX

ATGTCTCAGGTTCAGAGTGGCATTTTGCCAGAACATTGCCGCGCGGCGATTTGGATCGAAGCCAACGTGAAAGGGGAAGTTGACGCCCTGCGTGCGGCCAGTAAAACATTTGCCGACAAACTGGCAACTTTTGAAGCGAAATTCCCGGACGCGCATCTTGGTGCGGTGGTTGCCTTTGGTAACAACACCTGGCGCGCTCTGAGCGGCGGCGTTGGGGCAGAAGAGCTGAAAGATTTTCCGGGCTACGGTAAAGGCCTTGCGCCGACGACCCAGTTCGATGTGTTGATCCACATTCTTTCTCTGCGTCACGACGTAAACTTCTCTGTCGCCCAGGCGGCGATGGAAGCCTTTGGTGACTGCATTGAAGTGAAAGAAGAGATCCACGGCTTCCGTTGGGTTGAAGAGCGT**GAC**CTGAGCGGCTTTGTT**GAC**GGTACGGAA**AAC**CCGGCGGGTGAAGAGACGCGTCGCGAAGTGGCGGTTATCAAAGACGGCGTGGATGCGGGCGGCAGCTATGTGTTTGTCCAGCGTTGGGAACACAACCTGAAGCAGCTCAACCGGATGAGCGTTCACGATCAGGAGATGGTGATCGGGCGCACCAAAGAGGCCAACGAAGAGATCGACGGCGACGAACGTCCGGAAACCTCTCACCTCACCCGCGTTGATCTGAAAGAAGATGGCAAAGGGCTGAAGATTGTT**CGC**CAG**AGC**CTGCCGTACGGCACTGCCAGTGGCACTCACGGTCTGTACTTCTGCGCCTACTGCGCGCGTCTGCATAACATTGAGCAGCAACTGCTGAGCATGTTTGGCGATACCGATGGTAAGCGTGATGCGATGTTGCGTTTCACCAAACCGGTAACCGGCGGCTATTATTTCGCACCGTCGCTGGACAAGTTGATGGCGCTGTAA

##### Restriction Enzyme

###### Level 0

BbsI

Recognition site: ***GAAGAC***

###### Level 2

BsaI

Recognition site: ***GGTCTC***

##### Envisioned Mutations

Aspartic Acid - 137
Aspartic Acid - 143
Asparagine - 147
Arginine - 232
Serine - 234

Substitute for NDT

#### R Workflow

suppressWarnings(suppressMessages(library("GoldenMutagenesis")))

input_sequence<-"ATGTCTCAGGTTCAGAGTGGCATTTTGCCAGAACATTGCCGCGCGGCGATTTGGATCGAAGCCAACGTGAAAGGGGAAGTTGACGCCCTGCGTGCGGCCAGTAAAACATTTGCCGACAAACTGGCAACTTTTGAAGCGAAATTCCCGGACGCGCATCTTGGTGCGGTGGTTGCCTTTGGTAACAACACCTGGCGCGCTCTGAGCGGCGGCGTTGGGGCAGAAGAGCTGAAAGATTTTCCGGGCTACGGTAAAGGCCTTGCGCCGACGACCCAGTTCGATGTGTTGATCCACATTCTTTCTCTGCGTCACGACGTAAACTTCTCTGTCGCCCAGGCGGCGATGGAAGCCTTTGGTGACTGCATTGAAGTGAAAGAAGAGATCCACGGCTTCCGTTGGGTTGAAGAGCGTGACCTGAGCGGCTTTGTTGACGGTACGGAAAACCCGGCGGGTGAAGAGACGCGTCGCGAAGTGGCGGTTATCAAAGACGGCGTGGATGCGGGCGGCAGCTATGTGTTTGTCCAGCGTTGGGAACACAACCTGAAGCAGCTCAACCGGATGAGCGTTCACGATCAGGAGATGGTGATCGGGCGCACCAAAGAGGCCAACGAAGAGATCGACGGCGACGAACGTCCGGAAACCTCTCACCTCACCCGCGTTGATCTGAAAGAAGATGGCAAAGGGCTGAAGATTGTTCGCCAGAGCCTGCCGTACGGCACTGCCAGTGGCACTCACGGTCTGTACTTCTGCGCCTACTGCGCGCGTCTGCATAACATTGAGCAGCAACTGCTGAGCATGTTTGGCGATACCGATGGTAAGCGTGATGCGATGTTGCGTTTCACCAAACCGGTAACCGGCGGCTATTATTTCGCACCGTCGCTGGACAAGTTGATGGCGCTGTAA"
recognition_site_bbsi<-"GAAGAC"
recognition_site_bsai<-"GGTCTC"
cuf<-"e_coli_316407.csv"

The domesticate function checks for internal cleavage sites. If corresponding sites are present silent mutations are introduced to destroy the recognition sites. The functions returns a list containing the position of the choosen amino acid residue for silent mutation.

```
mutations_bbsi<-domesticate(input_sequence, recognition_site_bbsi, cuf=cuf)
## [1] "No domestication needed."
mutations_bbsi
## list()
mutations_bsai<-domesticate(input_sequence, recognition_site_bsai, cuf=cuf)
## [1] "No domestication needed."
mutations_bsai
## list()
```

The mutate_msd function designs the necessary set of primers for the desired mutations. 
The function has the following parameters:
**prefix**: Additional nucleobases in 5’ position of the recognition site [default: TT]
**restriction_enzym**: Recognition site sequence of the respective restriction enzyme [default: GGTCTC]
**codon**: The codon which should be used in the mutagenesis [default: NDT]
**suffix**: Spacer nucleotides matching the cleavage pattern of the enzyme [default: A]
**vector**: Four basepair overhangs complementary to the created overhangs in the acceptor vector [default: c(“AATG”, “AAGC”)]
**replacements**: The desired substitutions as a vector with positions OR a list containing vetors with position (char) and type of MSD mutation (char) (see MSD3.rd for an example)
**binding_min_length**: The minimal threshold value of the template binding sequence in amino acid residues [default: 4]
**primer_length**: Maximal length of the binding sequence [default: 9]
**target_temp**: Melting temperature of the binding sequence in °C [default: 60]
**replacement_range**: Maximum distance between two randomization sites to be incoporated into a single primer in amino acid residues [default: 5]
**fragment_min_size**: Minimal size of a generated gene fragment in base pairs [default 100]

It will return an object of the class Primerset.
The primers for multiple site saturation mutagenesis have an additional slot called “NDT”. This slot contains a non-binding region in which (the) NDT site(s) is/are located.

```
#If domestication is necessary follow the workflow of the Point Mutagenesis vignette
mutations<-c(137,143,147,232,234)
primers<-msd_mutate(input_sequence, prefix="TT" ,restriction_enzyme=recognition_site_bsai, suffix="A", vector=c("AATG", "AAGC"), replacements=mutations, replacement_range=5, binding_min_length=4 ,primer_length=9, target_temp=60, fragment_min_size=60 )
primers
## An object of class "Extended Primerset"
## Slot "fragments":
## [[1]]
## An object of class "Fragment"
## Slot "start":
## [1] 2
##
## Slot "stop":
## [1] 142
##
## Slot "start_mutation":
## logical(0)
##
## Slot "stop_mutation":
## [1] 137
##
##
## [[2]]
## An object of class "Fragment"
## Slot "start":
## [1] 143
##
## Slot "stop":
## [1] 231
##
## Slot "start_mutation":
## [1] 143 147
##
## Slot "stop_mutation":
## logical(0)
##
##
## [[3]]
## An object of class "Fragment"
## Slot "start":
## [1] 232
##
## Slot "stop":
## [1] 300
##
## Slot "start_mutation":
## [1] 232 234
##
## Slot "stop_mutation":
## logical(0)
##
##
##
## Slot "oldsequence":
## [1] "ATGTCTCAGGTTCAGAGTGGCATTTTGCCAGAACATTGCCGCGCGGCGATTTGGATCGAAGCCAACGTGAAAGGGGAAGTTGACGCCCTGCGTGCGGCCAGTAAAACATTTGCCGACAAACTGGCAACTTTTGAAGCGAAATTCCCGGACGCGCATCTTGGTGCGGTGGTTGCCTTTGGTAACAACACCTGGCGCGCTCTGAGCGGCGGCGTTGGGGCAGAAGAGCTGAAAGATTTTCCGGGCTACGGTAAAGGCCTTGCGCCGACGACCCAGTTCGATGTGTTGATCCACATTCTTTCTCTGCGTCACGACGTAAACTTCTCTGTCGCCCAGGCGGCGATGGAAGCCTTTGGTGACTGCATTGAAGTGAAAGAAGAGATCCACGGCTTCCGTTGGGTTGAAGAGCGTGACCTGAGCGGCTTTGTTGACGGTACGGAAAACCCGGCGGGTGAAGAGACGCGTCGCGAAGTGGCGGTTATCAAAGACGGCGTGGATGCGGGCGGCAGCTATGTGTTTGTCCAGCGTTGGGAACACAACCTGAAGCAGCTCAACCGGATGAGCGTTCACGATCAGGAGATGGTGATCGGGCGCACCAAAGAGGCCAACGAAGAGATCGACGGCGACGAACGTCCGGAAACCTCTCACCTCACCCGCGTTGATCTGAAAGAAGATGGCAAAGGGCTGAAGATTGTTCGCCAGAGCCTGCCGTACGGCACTGCCAGTGGCACTCACGGTCTGTACTTCTGCGCCTACTGCGCGCGTCTGCATAACATTGAGCAGCAACTGCTGAGCATGTTTGGCGATACCGATGGTAAGCGTGATGCGATGTTGCGTTTCACCAAACCGGTAACCGGCGGCTATTATTTCGCACCGTCGCTGGACAAGTTGATGGCGCTGTAA"
##
## Slot "primers":
## [[1]]
## [[1]][[1]]
## An object of class "Primer"
## Slot "prefix":
## [1] "TT"
##
## Slot "restriction_enzyme":
## [1] "GGTCTC"
##
## Slot "suffix":
## [1] "A"
##
## Slot "vector":
## [1] "AATG"
##
## Slot "overhang":
## [1] ""
##
## Slot "binding_sequence":
## [1] "TCTCAGGTTCAGAGTGGCATTTTGCC"
##
## Slot "temperature":
## [1] 60.36446
##
## Slot "difference":
## [1] 0.3644563
##
##
## [[1]][[2]]
## An object of class "Primer MSD"
## Slot "NDT":
## [1] "AGCCGCTCAGAHN"
##
## Slot "prefix":
## [1] "TT"
##
## Slot "restriction_enzyme":
## [1] "GGTCTC"
##
## Slot "suffix":
## [1] "T"
##
## Slot "vector":
## [1] ""
##
## Slot "overhang":
## [1] "ACAA"
##
## Slot "binding_sequence":
## [1] "ACGCTCTTCAACCCAACGGAAGC"
##
## Slot "temperature":
## [1] 59.785
##
## Slot "difference":
## [1] 0.5794564
##
##
##
## [[2]]
## [[2]][[1]]
## An object of class "Primer MSD"
## Slot "NDT":
## [1] "TNDTGGTACGGAANDT"
##
## Slot "prefix":
## [1] "TT"
##
## Slot "restriction_enzyme":
## [1] "GGTCTC"
##
## Slot "suffix":
## [1] "A"
##
## Slot "vector":
## [1] ""
##
## Slot "overhang":
## [1] "TTGT"
##
## Slot "binding_sequence":
## [1] "CCGGCGGGTGAAGAGACGC"
##
## Slot "temperature":
## [1] 59.2169
##
## Slot "difference":
## [1] 0.7831005
##
##
## [[2]][[2]]
## An object of class "Primer"
## Slot "prefix":
## [1] "TT"
##
## Slot "restriction_enzyme":
## [1] "GGTCTC"
##
## Slot "suffix":
## [1] "T"
##
## Slot "vector":
## [1] ""
##
## Slot "overhang":
## [1] "AACA"
##
## Slot "binding_sequence":
## [1] "ATCTTCAGCCCTTTGCCATCTTCTTT"
##
## Slot "temperature":
## [1] 59.04605
##
## Slot "difference":
## [1] 0.1708527
##
##
##
## [[3]]
## [[3]][[1]]
## An object of class "Primer MSD"
## Slot "NDT":
## [1] "NDTCAGNDT"
##
## Slot "prefix":
## [1] "TT"
##
## Slot "restriction_enzyme":
## [1] "GGTCTC"
##
## Slot "suffix":
## [1] "A"
##
## Slot "vector":
## [1] ""
##
## Slot "overhang":
## [1] "TGTT"
##
## Slot "binding_sequence":
## [1] "CTGCCGTACGGCACTGCCAG"
##
## Slot "temperature":
## [1] 60.52991
##
## Slot "difference":
## [1] 0.5299114
##
##
## [[3]][[2]]
## An object of class "Primer"
## Slot "prefix":
## [1] "TT"
##
## Slot "restriction_enzyme":
## [1] "GGTCTC"
##
## Slot "suffix":
## [1] "T"
##
## Slot "vector":
## [1] "AAGC"
##
## Slot "overhang":
## [1] ""
##
## Slot "binding_sequence":
## [1] "TTACAGCGCCATCAACTTGTCCAGC"
##
## Slot "temperature":
## [1] 60.90327
##
## Slot "difference":
## [1] 0.3733614
##
##
##
##
## Slot "newsequence":
## [1] "ATGTCTCAGGTTCAGAGTGGCATTTTGCCAGAACATTGCCGCGCGGCGATTTGGATCGAAGCCAACGTGAAAGGGGAAGTTGACGCCCTGCGTGCGGCCAGTAAAACATTTGCCGACAAACTGGCAACTTTTGAAGCGAAATTCCCGGACGCGCATCTTGGTGCGGTGGTTGCCTTTGGTAACAACACCTGGCGCGCTCTGAGCGGCGGCGTTGGGGCAGAAGAGCTGAAAGATTTTCCGGGCTACGGTAAAGGCCTTGCGCCGACGACCCAGTTCGATGTGTTGATCCACATTCTTTCTCTGCGTCACGACGTAAACTTCTCTGTCGCCCAGGCGGCGATGGAAGCCTTTGGTGACTGCATTGAAGTGAAAGAAGAGATCCACGGCTTCCGTTGGGTTGAAGAGCGTNDTCTGAGCGGCTTTGTTNDTGGTACGGAANDTCCGGCGGGTGAAGAGACGCGTCGCGAAGTGGCGGTTATCAAAGACGGCGTGGATGCGGGCGGCAGCTATGTGTTTGTCCAGCGTTGGGAACACAACCTGAAGCAGCTCAACCGGATGAGCGTTCACGATCAGGAGATGGTGATCGGGCGCACCAAAGAGGCCAACGAAGAGATCGACGGCGACGAACGTCCGGAAACCTCTCACCTCACCCGCGTTGATCTGAAAGAAGATGGCAAAGGGCTGAAGATTGTTNDTCAGNDTCTGCCGTACGGCACTGCCAGTGGCACTCACGGTCTGTACTTCTGCGCCTACTGCGCGCGTCTGCATAACATTGAGCAGCAACTGCTGAGCATGTTTGGCGATACCGATGGTAAGCGTGATGCGATGTTGCGTTTCACCAAACCGGTAACCGGCGGCTATTATTTCGCACCGTCGCTGGACAAGTTGATGGCGCTGTAA"
```

The primers are generated for direct cloning into the Level 2 vector.
The function primer_add_level modifies the primers for individual cloning into Level 0 vectors and subsequent assembly in Level 2.
The parameters **prefix, restriction_enzyme, suffix and vector** can be set similar to the mutate-function.

```
primers_lvl0<-primer_add_level(primers,  prefix="TT", restriction_enzyme=recognition_site_bbsi, suffix="AA", vector=c("CTCA", "CTCG"))
primers_lvl0
## An object of class "Extended Primerset"
## Slot "fragments":
## [[1]]
## An object of class "Fragment"
## Slot "start":
## [1] 2
##
## Slot "stop":
## [1] 142
##
## Slot "start_mutation":
## logical(0)
##
## Slot "stop_mutation":
## [1] 137
##
##
## [[2]]
## An object of class "Fragment"
## Slot "start":
## [1] 143
##
## Slot "stop":
## [1] 231
##
## Slot "start_mutation":
## [1] 143 147
##
## Slot "stop_mutation":
## logical(0)
##
##
## [[3]]
## An object of class "Fragment"
## Slot "start":
## [1] 232
##
## Slot "stop":
## [1] 300
##
## Slot "start_mutation":
## [1] 232 234
##
## Slot "stop_mutation":
## logical(0)
##
##
##
## Slot "oldsequence":
## [1] "ATGTCTCAGGTTCAGAGTGGCATTTTGCCAGAACATTGCCGCGCGGCGATTTGGATCGAAGCCAACGTGAAAGGGGAAGTTGACGCCCTGCGTGCGGCCAGTAAAACATTTGCCGACAAACTGGCAACTTTTGAAGCGAAATTCCCGGACGCGCATCTTGGTGCGGTGGTTGCCTTTGGTAACAACACCTGGCGCGCTCTGAGCGGCGGCGTTGGGGCAGAAGAGCTGAAAGATTTTCCGGGCTACGGTAAAGGCCTTGCGCCGACGACCCAGTTCGATGTGTTGATCCACATTCTTTCTCTGCGTCACGACGTAAACTTCTCTGTCGCCCAGGCGGCGATGGAAGCCTTTGGTGACTGCATTGAAGTGAAAGAAGAGATCCACGGCTTCCGTTGGGTTGAAGAGCGTGACCTGAGCGGCTTTGTTGACGGTACGGAAAACCCGGCGGGTGAAGAGACGCGTCGCGAAGTGGCGGTTATCAAAGACGGCGTGGATGCGGGCGGCAGCTATGTGTTTGTCCAGCGTTGGGAACACAACCTGAAGCAGCTCAACCGGATGAGCGTTCACGATCAGGAGATGGTGATCGGGCGCACCAAAGAGGCCAACGAAGAGATCGACGGCGACGAACGTCCGGAAACCTCTCACCTCACCCGCGTTGATCTGAAAGAAGATGGCAAAGGGCTGAAGATTGTTCGCCAGAGCCTGCCGTACGGCACTGCCAGTGGCACTCACGGTCTGTACTTCTGCGCCTACTGCGCGCGTCTGCATAACATTGAGCAGCAACTGCTGAGCATGTTTGGCGATACCGATGGTAAGCGTGATGCGATGTTGCGTTTCACCAAACCGGTAACCGGCGGCTATTATTTCGCACCGTCGCTGGACAAGTTGATGGCGCTGTAA"
##
## Slot "primers":
## [[1]]
## [[1]][[1]]
## An object of class "Primer"
## Slot "prefix":
## [1] "TT"
##
## Slot "restriction_enzyme":
## [1] "GAAGAC"
##
## Slot "suffix":
## [1] "AA"
##
## Slot "vector":
## [1] "CTCA"
##
## Slot "overhang":
## [1] "AATG"
##
## Slot "binding_sequence":
## [1] "TCTCAGGTTCAGAGTGGCATTTTGCC"
##
## Slot "temperature":
## [1] 60.36446
##
## Slot "difference":
## [1] 0.3644563
##
##
## [[1]][[2]]
## An object of class "Primer MSD"
## Slot "NDT":
## [1] "AGCCGCTCAGAHN"
##
## Slot "prefix":
## [1] "TT"
##
## Slot "restriction_enzyme":
## [1] "GAAGAC"
##
## Slot "suffix":
## [1] "AA"
##
## Slot "vector":
## [1] "CTCG"
##
## Slot "overhang":
## [1] "ACAA"
##
## Slot "binding_sequence":
## [1] "ACGCTCTTCAACCCAACGGAAGC"
##
## Slot "temperature":
## [1] 59.785
##
## Slot "difference":
## [1] 0.5794564
##
##
##
## [[2]]
## [[2]][[1]]
## An object of class "Primer MSD"
## Slot "NDT":
## [1] "TNDTGGTACGGAANDT"
##
## Slot "prefix":
## [1] "TT"
##
## Slot "restriction_enzyme":
## [1] "GAAGAC"
##
## Slot "suffix":
## [1] "AA"
##
## Slot "vector":
## [1] "CTCA"
##
## Slot "overhang":
## [1] "TTGT"
##
## Slot "binding_sequence":
## [1] "CCGGCGGGTGAAGAGACGC"
##
## Slot "temperature":
## [1] 59.2169
##
## Slot "difference":
## [1] 0.7831005
##
##
## [[2]][[2]]
## An object of class "Primer"
## Slot "prefix":
## [1] "TT"
##
## Slot "restriction_enzyme":
## [1] "GAAGAC"
##
## Slot "suffix":
## [1] "AA"
##
## Slot "vector":
## [1] "CTCG"
##
## Slot "overhang":
## [1] "AACA"
##
## Slot "binding_sequence":
## [1] "ATCTTCAGCCCTTTGCCATCTTCTTT"
##
## Slot "temperature":
## [1] 59.04605
##
## Slot "difference":
## [1] 0.1708527
##
##
##
## [[3]]
## [[3]][[1]]
## An object of class "Primer MSD"
## Slot "NDT":
## [1] "NDTCAGNDT"
##
## Slot "prefix":
## [1] "TT"
##
## Slot "restriction_enzyme":
## [1] "GAAGAC"
##
## Slot "suffix":
## [1] "AA"
##
## Slot "vector":
## [1] "CTCA"
##
## Slot "overhang":
## [1] "TGTT"
##
## Slot "binding_sequence":
## [1] "CTGCCGTACGGCACTGCCAG"
##
## Slot "temperature":
## [1] 60.52991
##
## Slot "difference":
## [1] 0.5299114
##
##
## [[3]][[2]]
## An object of class "Primer"
## Slot "prefix":
## [1] "TT"
##
## Slot "restriction_enzyme":
## [1] "GAAGAC"
##
## Slot "suffix":
## [1] "AA"
##
## Slot "vector":
## [1] "CTCG"
##
## Slot "overhang":
## [1] "AAGC"
##
## Slot "binding_sequence":
## [1] "TTACAGCGCCATCAACTTGTCCAGC"
##
## Slot "temperature":
## [1] 60.90327
##
## Slot "difference":
## [1] 0.3733614
##
##
##
##
## Slot "newsequence":
## [1] "ATGTCTCAGGTTCAGAGTGGCATTTTGCCAGAACATTGCCGCGCGGCGATTTGGATCGAAGCCAACGTGAAAGGGGAAGTTGACGCCCTGCGTGCGGCCAGTAAAACATTTGCCGACAAACTGGCAACTTTTGAAGCGAAATTCCCGGACGCGCATCTTGGTGCGGTGGTTGCCTTTGGTAACAACACCTGGCGCGCTCTGAGCGGCGGCGTTGGGGCAGAAGAGCTGAAAGATTTTCCGGGCTACGGTAAAGGCCTTGCGCCGACGACCCAGTTCGATGTGTTGATCCACATTCTTTCTCTGCGTCACGACGTAAACTTCTCTGTCGCCCAGGCGGCGATGGAAGCCTTTGGTGACTGCATTGAAGTGAAAGAAGAGATCCACGGCTTCCGTTGGGTTGAAGAGCGTNDTCTGAGCGGCTTTGTTNDTGGTACGGAANDTCCGGCGGGTGAAGAGACGCGTCGCGAAGTGGCGGTTATCAAAGACGGCGTGGATGCGGGCGGCAGCTATGTGTTTGTCCAGCGTTGGGAACACAACCTGAAGCAGCTCAACCGGATGAGCGTTCACGATCAGGAGATGGTGATCGGGCGCACCAAAGAGGCCAACGAAGAGATCGACGGCGACGAACGTCCGGAAACCTCTCACCTCACCCGCGTTGATCTGAAAGAAGATGGCAAAGGGCTGAAGATTGTTNDTCAGNDTCTGCCGTACGGCACTGCCAGTGGCACTCACGGTCTGTACTTCTGCGCCTACTGCGCGCGTCTGCATAACATTGAGCAGCAACTGCTGAGCATGTTTGGCGATACCGATGGTAAGCGTGATGCGATGTTGCGTTTCACCAAACCGGTAACCGGCGGCTATTATTTCGCACCGTCGCTGGACAAGTTGATGGCGCTGTAA"
```

Objects of the classes “Primer”, “Primer MSD” and “Primerset” can have a slim textual output by using the function print_primer.

```
print_primer(primers_lvl0)
## Fragment 1
## Start 2, Stop 142, Length 141
## Forward
## TTGAAGACAACTCAAATGTCTCAGGTTCAGAGTGGCATTTTGCC
## Temperature of binding site:  60.36446  °C
## Temperature difference:  0.3644563  K
## Reverse
## TTGAAGACAACTCGACAAAGCCGCTCAGAHNACGCTCTTCAACCCAACGGAAGC
## Temperature of binding site:  59.785  °C
## Temperature difference:  0.5794564  K
##
## Fragment 2
## Start 143, Stop 231, Length 89
## Forward
## TTGAAGACAACTCATTGTTNDTGGTACGGAANDTCCGGCGGGTGAAGAGACGC
## Temperature of binding site:  59.2169  °C
## Temperature difference:  0.7831005  K
## Reverse
## TTGAAGACAACTCGAACAATCTTCAGCCCTTTGCCATCTTCTTT
## Temperature of binding site:  59.04605  °C
## Temperature difference:  0.1708527  K
##
## Fragment 3
## Start 232, Stop 300, Length 69
## Forward
## TTGAAGACAACTCATGTTNDTCAGNDTCTGCCGTACGGCACTGCCAG
## Temperature of binding site:  60.52991  °C
## Temperature difference:  0.5299114  K
## Reverse
## TTGAAGACAACTCGAAGCTTACAGCGCCATCAACTTGTCCAGC
## Temperature of binding site:  60.90327  °C
## Temperature difference:  0.3733614  K
##
## Input Sequence:
##  ATGTCTCAGGTTCAGAGTGGCATTTTGCCAGAACATTGCCGCGCGGCGATTTGGATCGAAGCCAACGTGAAAGGGGAAGTTGACGCCCTGCGTGCGGCCAGTAAAACATTTGCCGACAAACTGGCAACTTTTGAAGCGAAATTCCCGGACGCGCATCTTGGTGCGGTGGTTGCCTTTGGTAACAACACCTGGCGCGCTCTGAGCGGCGGCGTTGGGGCAGAAGAGCTGAAAGATTTTCCGGGCTACGGTAAAGGCCTTGCGCCGACGACCCAGTTCGATGTGTTGATCCACATTCTTTCTCTGCGTCACGACGTAAACTTCTCTGTCGCCCAGGCGGCGATGGAAGCCTTTGGTGACTGCATTGAAGTGAAAGAAGAGATCCACGGCTTCCGTTGGGTTGAAGAGCGTGACCTGAGCGGCTTTGTTGACGGTACGGAAAACCCGGCGGGTGAAGAGACGCGTCGCGAAGTGGCGGTTATCAAAGACGGCGTGGATGCGGGCGGCAGCTATGTGTTTGTCCAGCGTTGGGAACACAACCTGAAGCAGCTCAACCGGATGAGCGTTCACGATCAGGAGATGGTGATCGGGCGCACCAAAGAGGCCAACGAAGAGATCGACGGCGACGAACGTCCGGAAACCTCTCACCTCACCCGCGTTGATCTGAAAGAAGATGGCAAAGGGCTGAAGATTGTTCGCCAGAGCCTGCCGTACGGCACTGCCAGTGGCACTCACGGTCTGTACTTCTGCGCCTACTGCGCGCGTCTGCATAACATTGAGCAGCAACTGCTGAGCATGTTTGGCGATACCGATGGTAAGCGTGATGCGATGTTGCGTTTCACCAAACCGGTAACCGGCGGCTATTATTTCGCACCGTCGCTGGACAAGTTGATGGCGCTGTAA
##
## Modified Sequence:
##  ATGTCTCAGGTTCAGAGTGGCATTTTGCCAGAACATTGCCGCGCGGCGATTTGGATCGAAGCCAACGTGAAAGGGGAAGTTGACGCCCTGCGTGCGGCCAGTAAAACATTTGCCGACAAACTGGCAACTTTTGAAGCGAAATTCCCGGACGCGCATCTTGGTGCGGTGGTTGCCTTTGGTAACAACACCTGGCGCGCTCTGAGCGGCGGCGTTGGGGCAGAAGAGCTGAAAGATTTTCCGGGCTACGGTAAAGGCCTTGCGCCGACGACCCAGTTCGATGTGTTGATCCACATTCTTTCTCTGCGTCACGACGTAAACTTCTCTGTCGCCCAGGCGGCGATGGAAGCCTTTGGTGACTGCATTGAAGTGAAAGAAGAGATCCACGGCTTCCGTTGGGTTGAAGAGCGTNDTCTGAGCGGCTTTGTTNDTGGTACGGAANDTCCGGCGGGTGAAGAGACGCGTCGCGAAGTGGCGGTTATCAAAGACGGCGTGGATGCGGGCGGCAGCTATGTGTTTGTCCAGCGTTGGGAACACAACCTGAAGCAGCTCAACCGGATGAGCGTTCACGATCAGGAGATGGTGATCGGGCGCACCAAAGAGGCCAACGAAGAGATCGACGGCGACGAACGTCCGGAAACCTCTCACCTCACCCGCGTTGATCTGAAAGAAGATGGCAAAGGGCTGAAGATTGTTNDTCAGNDTCTGCCGTACGGCACTGCCAGTGGCACTCACGGTCTGTACTTCTGCGCCTACTGCGCGCGTCTGCATAACATTGAGCAGCAACTGCTGAGCATGTTTGGCGATACCGATGGTAAGCGTGATGCGATGTTGCGTTTCACCAAACCGGTAACCGGCGGCTATTATTTCGCACCGTCGCTGGACAAGTTGATGGCGCTGTAA
```

The textual output can be printed into a file.

sink("primers.txt", append=FALSE, split=FALSE)
print_primer(primers_lvl0)
sink()
