## Supplementary material for "Golden Mutagenesis: An efficient multi-site-saturation mutagenesis approach by Golden Gate cloning with automated primer design": Computer Tool example Multi-Site Directed Mutagenesis 2

Multiple Site Saturation Mutagenesis Example 2

2018-10-10

### Multiple Site Saturation Mutagenesis

You can find the lastest version of this file at <https://github.com/ipb-halle/GoldenMutagenesis/blob/master/vignettes/MSD2.md>

#### Experimental Workflow

##### Target sequence

open reading frame YfeX in pCA24N (Chloramphenicol resistance)

##### Clone into

seperate gene fragments into pAGM9121 first then reassemble into pAGM22082_cRed

##### Genomic sequence YfeX

ATGTCTCAGGTTCAGAGTGGCATTTTGCCAGAACATTGCCGCGCGGCGATTTGGATCGAAGCCAACGTGAAAGGGGAAGTTGACGCCCTGCGTGCGGCCAGTAAAACATTTGCCGACAAACTGGCAACTTTTGAAGCGAAATTCCCGGAC**GCG**CATCTTGGTGCGGTGGTTGCCTTTGGTAACAACACCTGGCGCGCTCTGAGCGGCGGCGTTGGGGCAGAAGAGCTGAAAGATTTTCCGGGCTACGGTAAAGGCCTTGCGCCGACGACCCAGTTCGATGTGTTGATCCACATTCTTTCT**CTG**CGTCACGACGTAAACTTCTCTGTCGCCCAGGCGGCGATGGAAGCCTTTGGTGACTGCATTGAAGTGAAAGAAGAGATCCACGGCTTCCGTTGGGTTGAAGAGCGTGACCTGAGCGGCTTTGTTGACGGTACGGAAAACCCGGCGGGT**GAA**GAGACGCGTCGCGAAGTGGCGGTTATCAAAGACGGCGTGGATGCGGGCGGCAGCTATGTGTTTGTCCAGCGTTGGGAACACAACCTGAAGCAGCTCAACCGGATGAGCGTTCACGATCAGGAGATGGTGATCGGGCGCACCAAAGAGGCC**AAC**GAAGAGATCGACGGCGACGAACGTCCGGAAACCTCTCACCTCACCCGCGTTGATCTGAAAGAAGATGGCAAAGGGCTGAAGATTGTTCGCCAGAGCCTGCCGTACGGCACTGCCAGTGGCACTCACGGTCTGTACTTCTGCGCCTAC**TGC**GCGCGTCTGCATAACATTGAGCAGCAACTGCTGAGCATGTTTGGCGATACCGATGGTAAGCGTGATGCGATGTTGCGTTTCACCAAACCGGTAACCGGCGGCTATTATTTCGCACCGTCGCTGGACAAGTTGATGGCGCTGTAA

##### Restriction Enzyme

###### Level 0

BbsI

Recognition site: ***GAAGAC***

###### Level 2

BsaI

Recognition site: ***GGTCTC***

##### Envisioned Mutations

Alanine - 51
Leucine - 101
Glutamic Acid - 151
Asparagine - 202
Cysteine - 252

Substitute for NDT

#### R Workflow

suppressWarnings(suppressMessages(library("GoldenMutagenesis")))

```
#If domestication is necessary follow the workflow of the Point Mutagenesis vignette
mutations<-c(51,101,151,202,252)
primers<-msd_mutate(input_sequence, prefix="TT" ,restriction_enzyme=recognition_site_bsai, suffix="A", vector=c("AATG", "AAGC"), replacements=mutations, replacement_range=5, binding_min_length=4 ,primer_length=9, target_temp=60, fragment_min_size=60 )
primers
## An object of class "Extended Primerset"
## Slot "fragments":
## [[1]]
## An object of class "Fragment"
## Slot "start":
## [1] 2
##
## Slot "stop":
## [1] 53
##
## Slot "start_mutation":
## logical(0)
##
## Slot "stop_mutation":
## [1] 51
##
##
## [[2]]
## An object of class "Fragment"
## Slot "start":
## [1] 54
##
## Slot "stop":
## [1] 103
##
## Slot "start_mutation":
## logical(0)
##
## Slot "stop_mutation":
## [1] 101
##
##
## [[3]]
## An object of class "Fragment"
## Slot "start":
## [1] 104
##
## Slot "stop":
## [1] 153
##
## Slot "start_mutation":
## logical(0)
##
## Slot "stop_mutation":
## [1] 151
##
##
## [[4]]
## An object of class "Fragment"
## Slot "start":
## [1] 154
##
## Slot "stop":
## [1] 204
##
## Slot "start_mutation":
## logical(0)
##
## Slot "stop_mutation":
## [1] 202
##
##
## [[5]]
## An object of class "Fragment"
## Slot "start":
## [1] 205
##
## Slot "stop":
## [1] 254
##
## Slot "start_mutation":
## logical(0)
##
## Slot "stop_mutation":
## [1] 252
##
##
## [[6]]
## An object of class "Fragment"
## Slot "start":
## [1] 255
##
## Slot "stop":
## [1] 300
##
## Slot "start_mutation":
## logical(0)
##
## Slot "stop_mutation":
## logical(0)
##
##
##
## Slot "oldsequence":
## [1] "ATGTCTCAGGTTCAGAGTGGCATTTTGCCAGAACATTGCCGCGCGGCGATTTGGATCGAAGCCAACGTGAAAGGGGAAGTTGACGCCCTGCGTGCGGCCAGTAAAACATTTGCCGACAAACTGGCAACTTTTGAAGCGAAATTCCCGGACGCGCATCTTGGTGCGGTGGTTGCCTTTGGTAACAACACCTGGCGCGCTCTGAGCGGCGGCGTTGGGGCAGAAGAGCTGAAAGATTTTCCGGGCTACGGTAAAGGCCTTGCGCCGACGACCCAGTTCGATGTGTTGATCCACATTCTTTCTCTGCGTCACGACGTAAACTTCTCTGTCGCCCAGGCGGCGATGGAAGCCTTTGGTGACTGCATTGAAGTGAAAGAAGAGATCCACGGCTTCCGTTGGGTTGAAGAGCGTGACCTGAGCGGCTTTGTTGACGGTACGGAAAACCCGGCGGGTGAAGAGACGCGTCGCGAAGTGGCGGTTATCAAAGACGGCGTGGATGCGGGCGGCAGCTATGTGTTTGTCCAGCGTTGGGAACACAACCTGAAGCAGCTCAACCGGATGAGCGTTCACGATCAGGAGATGGTGATCGGGCGCACCAAAGAGGCCAACGAAGAGATCGACGGCGACGAACGTCCGGAAACCTCTCACCTCACCCGCGTTGATCTGAAAGAAGATGGCAAAGGGCTGAAGATTGTTCGCCAGAGCCTGCCGTACGGCACTGCCAGTGGCACTCACGGTCTGTACTTCTGCGCCTACTGCGCGCGTCTGCATAACATTGAGCAGCAACTGCTGAGCATGTTTGGCGATACCGATGGTAAGCGTGATGCGATGTTGCGTTTCACCAAACCGGTAACCGGCGGCTATTATTTCGCACCGTCGCTGGACAAGTTGATGGCGCTGTAA"
##
## Slot "primers":
## [[1]]
## [[1]][[1]]
## An object of class "Primer"
## Slot "prefix":
## [1] "TT"
##
## Slot "restriction_enzyme":
## [1] "GGTCTC"
##
## Slot "suffix":
## [1] "A"
##
## Slot "vector":
## [1] "AATG"
##
## Slot "overhang":
## [1] ""
##
## Slot "binding_sequence":
## [1] "TCTCAGGTTCAGAGTGGCATTTTGCC"
##
## Slot "temperature":
## [1] 60.36446
##
## Slot "difference":
## [1] 0.3644563
##
##
## [[1]][[2]]
## An object of class "Primer MSD"
## Slot "NDT":
## [1] "TGAHN"
##
## Slot "prefix":
## [1] "TT"
##
## Slot "restriction_enzyme":
## [1] "GGTCTC"
##
## Slot "suffix":
## [1] "T"
##
## Slot "vector":
## [1] ""
##
## Slot "overhang":
## [1] "AAGA"
##
## Slot "binding_sequence":
## [1] "GTCCGGGAATTTCGCTTCAAAAGTTGC"
##
## Slot "temperature":
## [1] 60.56425
##
## Slot "difference":
## [1] 0.1997931
##
##
##
## [[2]]
## [[2]][[1]]
## An object of class "Primer"
## Slot "prefix":
## [1] "TT"
##
## Slot "restriction_enzyme":
## [1] "GGTCTC"
##
## Slot "suffix":
## [1] "A"
##
## Slot "vector":
## [1] ""
##
## Slot "overhang":
## [1] "TCTT"
##
## Slot "binding_sequence":
## [1] "GGTGCGGTGGTTGCCTTTGGTAACA"
##
## Slot "temperature":
## [1] 60.1791
##
## Slot "difference":
## [1] 0.1790953
##
##
## [[2]][[2]]
## An object of class "Primer MSD"
## Slot "NDT":
## [1] "CGAHN"
##
## Slot "prefix":
## [1] "TT"
##
## Slot "restriction_enzyme":
## [1] "GGTCTC"
##
## Slot "suffix":
## [1] "T"
##
## Slot "vector":
## [1] ""
##
## Slot "overhang":
## [1] "GTGA"
##
## Slot "binding_sequence":
## [1] "AGAAAGAATGTGGATCAACACATCGAACTG"
##
## Slot "temperature":
## [1] 60.11394
##
## Slot "difference":
## [1] 0.06515983
##
##
##
## [[3]]
## [[3]][[1]]
## An object of class "Primer"
## Slot "prefix":
## [1] "TT"
##
## Slot "restriction_enzyme":
## [1] "GGTCTC"
##
## Slot "suffix":
## [1] "A"
##
## Slot "vector":
## [1] ""
##
## Slot "overhang":
## [1] "TCAC"
##
## Slot "binding_sequence":
## [1] "GACGTAAACTTCTCTGTCGCCCAGG"
##
## Slot "temperature":
## [1] 60.33165
##
## Slot "difference":
## [1] 0.3316526
##
##
## [[3]][[2]]
## An object of class "Primer MSD"
## Slot "NDT":
## [1] "TCAHN"
##
## Slot "prefix":
## [1] "TT"
##
## Slot "restriction_enzyme":
## [1] "GGTCTC"
##
## Slot "suffix":
## [1] "T"
##
## Slot "vector":
## [1] ""
##
## Slot "overhang":
## [1] "CGTC"
##
## Slot "binding_sequence":
## [1] "ACCCGCCGGGTTTTCCGTACCG"
##
## Slot "temperature":
## [1] 60.85613
##
## Slot "difference":
## [1] 0.5244756
##
##
##
## [[4]]
## [[4]][[1]]
## An object of class "Primer"
## Slot "prefix":
## [1] "TT"
##
## Slot "restriction_enzyme":
## [1] "GGTCTC"
##
## Slot "suffix":
## [1] "A"
##
## Slot "vector":
## [1] ""
##
## Slot "overhang":
## [1] "GACG"
##
## Slot "binding_sequence":
## [1] "CGTCGCGAAGTGGCGGTTATCA"
##
## Slot "temperature":
## [1] 60.61078
##
## Slot "difference":
## [1] 0.6107759
##
##
## [[4]][[2]]
## An object of class "Primer MSD"
## Slot "NDT":
## [1] "TCAHN"
##
## Slot "prefix":
## [1] "TT"
##
## Slot "restriction_enzyme":
## [1] "GGTCTC"
##
## Slot "suffix":
## [1] "T"
##
## Slot "vector":
## [1] ""
##
## Slot "overhang":
## [1] "CTCT"
##
## Slot "binding_sequence":
## [1] "GGCCTCTTTGGTGCGCCCGA"
##
## Slot "temperature":
## [1] 60.47119
##
## Slot "difference":
## [1] 0.1395836
##
##
##
## [[5]]
## [[5]][[1]]
## An object of class "Primer"
## Slot "prefix":
## [1] "TT"
##
## Slot "restriction_enzyme":
## [1] "GGTCTC"
##
## Slot "suffix":
## [1] "A"
##
## Slot "vector":
## [1] ""
##
## Slot "overhang":
## [1] "AGAG"
##
## Slot "binding_sequence":
## [1] "ATCGACGGCGACGAACGTCC"
##
## Slot "temperature":
## [1] 58.83656
##
## Slot "difference":
## [1] 1.163437
##
##
## [[5]][[2]]
## An object of class "Primer MSD"
## Slot "NDT":
## [1] "GCAHN"
##
## Slot "prefix":
## [1] "TT"
##
## Slot "restriction_enzyme":
## [1] "GGTCTC"
##
## Slot "suffix":
## [1] "T"
##
## Slot "vector":
## [1] ""
##
## Slot "overhang":
## [1] "ACGC"
##
## Slot "binding_sequence":
## [1] "GTAGGCGCAGAAGTACAGACCGT"
##
## Slot "temperature":
## [1] 59.13423
##
## Slot "difference":
## [1] 0.2976635
##
##
##
## [[6]]
## [[6]][[1]]
## An object of class "Primer"
## Slot "prefix":
## [1] "TT"
##
## Slot "restriction_enzyme":
## [1] "GGTCTC"
##
## Slot "suffix":
## [1] "A"
##
## Slot "vector":
## [1] ""
##
## Slot "overhang":
## [1] "GCGT"
##
## Slot "binding_sequence":
## [1] "CTGCATAACATTGAGCAGCAACTGC"
##
## Slot "temperature":
## [1] 60.28916
##
## Slot "difference":
## [1] 0.2891579
##
##
## [[6]][[2]]
## An object of class "Primer"
## Slot "prefix":
## [1] "TT"
##
## Slot "restriction_enzyme":
## [1] "GGTCTC"
##
## Slot "suffix":
## [1] "T"
##
## Slot "vector":
## [1] "AAGC"
##
## Slot "overhang":
## [1] ""
##
## Slot "binding_sequence":
## [1] "TTACAGCGCCATCAACTTGTCCAGC"
##
## Slot "temperature":
## [1] 60.90327
##
## Slot "difference":
## [1] 0.6141149
##
##
##
##
## Slot "newsequence":
## [1] "ATGTCTCAGGTTCAGAGTGGCATTTTGCCAGAACATTGCCGCGCGGCGATTTGGATCGAAGCCAACGTGAAAGGGGAAGTTGACGCCCTGCGTGCGGCCAGTAAAACATTTGCCGACAAACTGGCAACTTTTGAAGCGAAATTCCCGGACNDTCATCTTGGTGCGGTGGTTGCCTTTGGTAACAACACCTGGCGCGCTCTGAGCGGCGGCGTTGGGGCAGAAGAGCTGAAAGATTTTCCGGGCTACGGTAAAGGCCTTGCGCCGACGACCCAGTTCGATGTGTTGATCCACATTCTTTCTNDTCGTCACGACGTAAACTTCTCTGTCGCCCAGGCGGCGATGGAAGCCTTTGGTGACTGCATTGAAGTGAAAGAAGAGATCCACGGCTTCCGTTGGGTTGAAGAGCGTGACCTGAGCGGCTTTGTTGACGGTACGGAAAACCCGGCGGGTNDTGAGACGCGTCGCGAAGTGGCGGTTATCAAAGACGGCGTGGATGCGGGCGGCAGCTATGTGTTTGTCCAGCGTTGGGAACACAACCTGAAGCAGCTCAACCGGATGAGCGTTCACGATCAGGAGATGGTGATCGGGCGCACCAAAGAGGCCNDTGAAGAGATCGACGGCGACGAACGTCCGGAAACCTCTCACCTCACCCGCGTTGATCTGAAAGAAGATGGCAAAGGGCTGAAGATTGTTCGCCAGAGCCTGCCGTACGGCACTGCCAGTGGCACTCACGGTCTGTACTTCTGCGCCTACNDTGCGCGTCTGCATAACATTGAGCAGCAACTGCTGAGCATGTTTGGCGATACCGATGGTAAGCGTGATGCGATGTTGCGTTTCACCAAACCGGTAACCGGCGGCTATTATTTCGCACCGTCGCTGGACAAGTTGATGGCGCTGTAA"
```

The primers are generated for direct cloning into the Level 2 vector.
The function primer_add_level modifies the primers for individual cloning into Level 0 vectors and subsequent assembly in Level 2.
The parameters **prefix, restriction_enzyme, suffix and vector** can be set similar to the mutate-function.

```
primers_lvl0<-primer_add_level(primers,  prefix="TT", restriction_enzyme=recognition_site_bbsi, suffix="AA", vector=c("CTCA", "CTCG"))
primers_lvl0
## An object of class "Extended Primerset"
## Slot "fragments":
## [[1]]
## An object of class "Fragment"
## Slot "start":
## [1] 2
##
## Slot "stop":
## [1] 53
##
## Slot "start_mutation":
## logical(0)
##
## Slot "stop_mutation":
## [1] 51
##
##
## [[2]]
## An object of class "Fragment"
## Slot "start":
## [1] 54
##
## Slot "stop":
## [1] 103
##
## Slot "start_mutation":
## logical(0)
##
## Slot "stop_mutation":
## [1] 101
##
##
## [[3]]
## An object of class "Fragment"
## Slot "start":
## [1] 104
##
## Slot "stop":
## [1] 153
##
## Slot "start_mutation":
## logical(0)
##
## Slot "stop_mutation":
## [1] 151
##
##
## [[4]]
## An object of class "Fragment"
## Slot "start":
## [1] 154
##
## Slot "stop":
## [1] 204
##
## Slot "start_mutation":
## logical(0)
##
## Slot "stop_mutation":
## [1] 202
##
##
## [[5]]
## An object of class "Fragment"
## Slot "start":
## [1] 205
##
## Slot "stop":
## [1] 254
##
## Slot "start_mutation":
## logical(0)
##
## Slot "stop_mutation":
## [1] 252
##
##
## [[6]]
## An object of class "Fragment"
## Slot "start":
## [1] 255
##
## Slot "stop":
## [1] 300
##
## Slot "start_mutation":
## logical(0)
##
## Slot "stop_mutation":
## logical(0)
##
##
##
## Slot "oldsequence":
## [1] "ATGTCTCAGGTTCAGAGTGGCATTTTGCCAGAACATTGCCGCGCGGCGATTTGGATCGAAGCCAACGTGAAAGGGGAAGTTGACGCCCTGCGTGCGGCCAGTAAAACATTTGCCGACAAACTGGCAACTTTTGAAGCGAAATTCCCGGACGCGCATCTTGGTGCGGTGGTTGCCTTTGGTAACAACACCTGGCGCGCTCTGAGCGGCGGCGTTGGGGCAGAAGAGCTGAAAGATTTTCCGGGCTACGGTAAAGGCCTTGCGCCGACGACCCAGTTCGATGTGTTGATCCACATTCTTTCTCTGCGTCACGACGTAAACTTCTCTGTCGCCCAGGCGGCGATGGAAGCCTTTGGTGACTGCATTGAAGTGAAAGAAGAGATCCACGGCTTCCGTTGGGTTGAAGAGCGTGACCTGAGCGGCTTTGTTGACGGTACGGAAAACCCGGCGGGTGAAGAGACGCGTCGCGAAGTGGCGGTTATCAAAGACGGCGTGGATGCGGGCGGCAGCTATGTGTTTGTCCAGCGTTGGGAACACAACCTGAAGCAGCTCAACCGGATGAGCGTTCACGATCAGGAGATGGTGATCGGGCGCACCAAAGAGGCCAACGAAGAGATCGACGGCGACGAACGTCCGGAAACCTCTCACCTCACCCGCGTTGATCTGAAAGAAGATGGCAAAGGGCTGAAGATTGTTCGCCAGAGCCTGCCGTACGGCACTGCCAGTGGCACTCACGGTCTGTACTTCTGCGCCTACTGCGCGCGTCTGCATAACATTGAGCAGCAACTGCTGAGCATGTTTGGCGATACCGATGGTAAGCGTGATGCGATGTTGCGTTTCACCAAACCGGTAACCGGCGGCTATTATTTCGCACCGTCGCTGGACAAGTTGATGGCGCTGTAA"
##
## Slot "primers":
## [[1]]
## [[1]][[1]]
## An object of class "Primer"
## Slot "prefix":
## [1] "TT"
##
## Slot "restriction_enzyme":
## [1] "GAAGAC"
##
## Slot "suffix":
## [1] "AA"
##
## Slot "vector":
## [1] "CTCA"
##
## Slot "overhang":
## [1] "AATG"
##
## Slot "binding_sequence":
## [1] "TCTCAGGTTCAGAGTGGCATTTTGCC"
##
## Slot "temperature":
## [1] 60.36446
##
## Slot "difference":
## [1] 0.3644563
##
##
## [[1]][[2]]
## An object of class "Primer MSD"
## Slot "NDT":
## [1] "TGAHN"
##
## Slot "prefix":
## [1] "TT"
##
## Slot "restriction_enzyme":
## [1] "GAAGAC"
##
## Slot "suffix":
## [1] "AA"
##
## Slot "vector":
## [1] "CTCG"
##
## Slot "overhang":
## [1] "AAGA"
##
## Slot "binding_sequence":
## [1] "GTCCGGGAATTTCGCTTCAAAAGTTGC"
##
## Slot "temperature":
## [1] 60.56425
##
## Slot "difference":
## [1] 0.1997931
##
##
##
## [[2]]
## [[2]][[1]]
## An object of class "Primer"
## Slot "prefix":
## [1] "TT"
##
## Slot "restriction_enzyme":
## [1] "GAAGAC"
##
## Slot "suffix":
## [1] "AA"
##
## Slot "vector":
## [1] "CTCA"
##
## Slot "overhang":
## [1] "TCTT"
##
## Slot "binding_sequence":
## [1] "GGTGCGGTGGTTGCCTTTGGTAACA"
##
## Slot "temperature":
## [1] 60.1791
##
## Slot "difference":
## [1] 0.1790953
##
##
## [[2]][[2]]
## An object of class "Primer MSD"
## Slot "NDT":
## [1] "CGAHN"
##
## Slot "prefix":
## [1] "TT"
##
## Slot "restriction_enzyme":
## [1] "GAAGAC"
##
## Slot "suffix":
## [1] "AA"
##
## Slot "vector":
## [1] "CTCG"
##
## Slot "overhang":
## [1] "GTGA"
##
## Slot "binding_sequence":
## [1] "AGAAAGAATGTGGATCAACACATCGAACTG"
##
## Slot "temperature":
## [1] 60.11394
##
## Slot "difference":
## [1] 0.06515983
##
##
##
## [[3]]
## [[3]][[1]]
## An object of class "Primer"
## Slot "prefix":
## [1] "TT"
##
## Slot "restriction_enzyme":
## [1] "GAAGAC"
##
## Slot "suffix":
## [1] "AA"
##
## Slot "vector":
## [1] "CTCA"
##
## Slot "overhang":
## [1] "TCAC"
##
## Slot "binding_sequence":
## [1] "GACGTAAACTTCTCTGTCGCCCAGG"
##
## Slot "temperature":
## [1] 60.33165
##
## Slot "difference":
## [1] 0.3316526
##
##
## [[3]][[2]]
## An object of class "Primer MSD"
## Slot "NDT":
## [1] "TCAHN"
##
## Slot "prefix":
## [1] "TT"
##
## Slot "restriction_enzyme":
## [1] "GAAGAC"
##
## Slot "suffix":
## [1] "AA"
##
## Slot "vector":
## [1] "CTCG"
##
## Slot "overhang":
## [1] "CGTC"
##
## Slot "binding_sequence":
## [1] "ACCCGCCGGGTTTTCCGTACCG"
##
## Slot "temperature":
## [1] 60.85613
##
## Slot "difference":
## [1] 0.5244756
##
##
##
## [[4]]
## [[4]][[1]]
## An object of class "Primer"
## Slot "prefix":
## [1] "TT"
##
## Slot "restriction_enzyme":
## [1] "GAAGAC"
##
## Slot "suffix":
## [1] "AA"
##
## Slot "vector":
## [1] "CTCA"
##
## Slot "overhang":
## [1] "GACG"
##
## Slot "binding_sequence":
## [1] "CGTCGCGAAGTGGCGGTTATCA"
##
## Slot "temperature":
## [1] 60.61078
##
## Slot "difference":
## [1] 0.6107759
##
##
## [[4]][[2]]
## An object of class "Primer MSD"
## Slot "NDT":
## [1] "TCAHN"
##
## Slot "prefix":
## [1] "TT"
##
## Slot "restriction_enzyme":
## [1] "GAAGAC"
##
## Slot "suffix":
## [1] "AA"
##
## Slot "vector":
## [1] "CTCG"
##
## Slot "overhang":
## [1] "CTCT"
##
## Slot "binding_sequence":
## [1] "GGCCTCTTTGGTGCGCCCGA"
##
## Slot "temperature":
## [1] 60.47119
##
## Slot "difference":
## [1] 0.1395836
##
##
##
## [[5]]
## [[5]][[1]]
## An object of class "Primer"
## Slot "prefix":
## [1] "TT"
##
## Slot "restriction_enzyme":
## [1] "GAAGAC"
##
## Slot "suffix":
## [1] "AA"
##
## Slot "vector":
## [1] "CTCA"
##
## Slot "overhang":
## [1] "AGAG"
##
## Slot "binding_sequence":
## [1] "ATCGACGGCGACGAACGTCC"
##
## Slot "temperature":
## [1] 58.83656
##
## Slot "difference":
## [1] 1.163437
##
##
## [[5]][[2]]
## An object of class "Primer MSD"
## Slot "NDT":
## [1] "GCAHN"
##
## Slot "prefix":
## [1] "TT"
##
## Slot "restriction_enzyme":
## [1] "GAAGAC"
##
## Slot "suffix":
## [1] "AA"
##
## Slot "vector":
## [1] "CTCG"
##
## Slot "overhang":
## [1] "ACGC"
##
## Slot "binding_sequence":
## [1] "GTAGGCGCAGAAGTACAGACCGT"
##
## Slot "temperature":
## [1] 59.13423
##
## Slot "difference":
## [1] 0.2976635
##
##
##
## [[6]]
## [[6]][[1]]
## An object of class "Primer"
## Slot "prefix":
## [1] "TT"
##
## Slot "restriction_enzyme":
## [1] "GAAGAC"
##
## Slot "suffix":
## [1] "AA"
##
## Slot "vector":
## [1] "CTCA"
##
## Slot "overhang":
## [1] "GCGT"
##
## Slot "binding_sequence":
## [1] "CTGCATAACATTGAGCAGCAACTGC"
##
## Slot "temperature":
## [1] 60.28916
##
## Slot "difference":
## [1] 0.2891579
##
##
## [[6]][[2]]
## An object of class "Primer"
## Slot "prefix":
## [1] "TT"
##
## Slot "restriction_enzyme":
## [1] "GAAGAC"
##
## Slot "suffix":
## [1] "AA"
##
## Slot "vector":
## [1] "CTCG"
##
## Slot "overhang":
## [1] "AAGC"
##
## Slot "binding_sequence":
## [1] "TTACAGCGCCATCAACTTGTCCAGC"
##
## Slot "temperature":
## [1] 60.90327
##
## Slot "difference":
## [1] 0.6141149
##
##
##
##
## Slot "newsequence":
## [1] "ATGTCTCAGGTTCAGAGTGGCATTTTGCCAGAACATTGCCGCGCGGCGATTTGGATCGAAGCCAACGTGAAAGGGGAAGTTGACGCCCTGCGTGCGGCCAGTAAAACATTTGCCGACAAACTGGCAACTTTTGAAGCGAAATTCCCGGACNDTCATCTTGGTGCGGTGGTTGCCTTTGGTAACAACACCTGGCGCGCTCTGAGCGGCGGCGTTGGGGCAGAAGAGCTGAAAGATTTTCCGGGCTACGGTAAAGGCCTTGCGCCGACGACCCAGTTCGATGTGTTGATCCACATTCTTTCTNDTCGTCACGACGTAAACTTCTCTGTCGCCCAGGCGGCGATGGAAGCCTTTGGTGACTGCATTGAAGTGAAAGAAGAGATCCACGGCTTCCGTTGGGTTGAAGAGCGTGACCTGAGCGGCTTTGTTGACGGTACGGAAAACCCGGCGGGTNDTGAGACGCGTCGCGAAGTGGCGGTTATCAAAGACGGCGTGGATGCGGGCGGCAGCTATGTGTTTGTCCAGCGTTGGGAACACAACCTGAAGCAGCTCAACCGGATGAGCGTTCACGATCAGGAGATGGTGATCGGGCGCACCAAAGAGGCCNDTGAAGAGATCGACGGCGACGAACGTCCGGAAACCTCTCACCTCACCCGCGTTGATCTGAAAGAAGATGGCAAAGGGCTGAAGATTGTTCGCCAGAGCCTGCCGTACGGCACTGCCAGTGGCACTCACGGTCTGTACTTCTGCGCCTACNDTGCGCGTCTGCATAACATTGAGCAGCAACTGCTGAGCATGTTTGGCGATACCGATGGTAAGCGTGATGCGATGTTGCGTTTCACCAAACCGGTAACCGGCGGCTATTATTTCGCACCGTCGCTGGACAAGTTGATGGCGCTGTAA"
```

Objects of the classes “Primer”, “Primer MSD” and “Primerset” can have a slim textual output by using the function print_primer.

```
print_primer(primers_lvl0)
## Fragment 1
## Start 2, Stop 53, Length 52
## Forward
## TTGAAGACAACTCAAATGTCTCAGGTTCAGAGTGGCATTTTGCC
## Temperature of binding site:  60.36446  °C
## Temperature difference:  0.3644563  K
## Reverse
## TTGAAGACAACTCGAAGATGAHNGTCCGGGAATTTCGCTTCAAAAGTTGC
## Temperature of binding site:  60.56425  °C
## Temperature difference:  0.1997931  K
##
## Fragment 2
## Start 54, Stop 103, Length 50
## Forward
## TTGAAGACAACTCATCTTGGTGCGGTGGTTGCCTTTGGTAACA
## Temperature of binding site:  60.1791  °C
## Temperature difference:  0.1790953  K
## Reverse
## TTGAAGACAACTCGGTGACGAHNAGAAAGAATGTGGATCAACACATCGAACTG
## Temperature of binding site:  60.11394  °C
## Temperature difference:  0.06515983  K
##
## Fragment 3
## Start 104, Stop 153, Length 50
## Forward
## TTGAAGACAACTCATCACGACGTAAACTTCTCTGTCGCCCAGG
## Temperature of binding site:  60.33165  °C
## Temperature difference:  0.3316526  K
## Reverse
## TTGAAGACAACTCGCGTCTCAHNACCCGCCGGGTTTTCCGTACCG
## Temperature of binding site:  60.85613  °C
## Temperature difference:  0.5244756  K
##
## Fragment 4
## Start 154, Stop 204, Length 51
## Forward
## TTGAAGACAACTCAGACGCGTCGCGAAGTGGCGGTTATCA
## Temperature of binding site:  60.61078  °C
## Temperature difference:  0.6107759  K
## Reverse
## TTGAAGACAACTCGCTCTTCAHNGGCCTCTTTGGTGCGCCCGA
## Temperature of binding site:  60.47119  °C
## Temperature difference:  0.1395836  K
##
## Fragment 5
## Start 205, Stop 254, Length 50
## Forward
## TTGAAGACAACTCAAGAGATCGACGGCGACGAACGTCC
## Temperature of binding site:  58.83656  °C
## Temperature difference:  1.163437  K
## Reverse
## TTGAAGACAACTCGACGCGCAHNGTAGGCGCAGAAGTACAGACCGT
## Temperature of binding site:  59.13423  °C
## Temperature difference:  0.2976635  K
##
## Fragment 6
## Start 255, Stop 300, Length 46
## Forward
## TTGAAGACAACTCAGCGTCTGCATAACATTGAGCAGCAACTGC
## Temperature of binding site:  60.28916  °C
## Temperature difference:  0.2891579  K
## Reverse
## TTGAAGACAACTCGAAGCTTACAGCGCCATCAACTTGTCCAGC
## Temperature of binding site:  60.90327  °C
## Temperature difference:  0.6141149  K
##
## Input Sequence:
##  ATGTCTCAGGTTCAGAGTGGCATTTTGCCAGAACATTGCCGCGCGGCGATTTGGATCGAAGCCAACGTGAAAGGGGAAGTTGACGCCCTGCGTGCGGCCAGTAAAACATTTGCCGACAAACTGGCAACTTTTGAAGCGAAATTCCCGGACGCGCATCTTGGTGCGGTGGTTGCCTTTGGTAACAACACCTGGCGCGCTCTGAGCGGCGGCGTTGGGGCAGAAGAGCTGAAAGATTTTCCGGGCTACGGTAAAGGCCTTGCGCCGACGACCCAGTTCGATGTGTTGATCCACATTCTTTCTCTGCGTCACGACGTAAACTTCTCTGTCGCCCAGGCGGCGATGGAAGCCTTTGGTGACTGCATTGAAGTGAAAGAAGAGATCCACGGCTTCCGTTGGGTTGAAGAGCGTGACCTGAGCGGCTTTGTTGACGGTACGGAAAACCCGGCGGGTGAAGAGACGCGTCGCGAAGTGGCGGTTATCAAAGACGGCGTGGATGCGGGCGGCAGCTATGTGTTTGTCCAGCGTTGGGAACACAACCTGAAGCAGCTCAACCGGATGAGCGTTCACGATCAGGAGATGGTGATCGGGCGCACCAAAGAGGCCAACGAAGAGATCGACGGCGACGAACGTCCGGAAACCTCTCACCTCACCCGCGTTGATCTGAAAGAAGATGGCAAAGGGCTGAAGATTGTTCGCCAGAGCCTGCCGTACGGCACTGCCAGTGGCACTCACGGTCTGTACTTCTGCGCCTACTGCGCGCGTCTGCATAACATTGAGCAGCAACTGCTGAGCATGTTTGGCGATACCGATGGTAAGCGTGATGCGATGTTGCGTTTCACCAAACCGGTAACCGGCGGCTATTATTTCGCACCGTCGCTGGACAAGTTGATGGCGCTGTAA
##
## Modified Sequence:
##  ATGTCTCAGGTTCAGAGTGGCATTTTGCCAGAACATTGCCGCGCGGCGATTTGGATCGAAGCCAACGTGAAAGGGGAAGTTGACGCCCTGCGTGCGGCCAGTAAAACATTTGCCGACAAACTGGCAACTTTTGAAGCGAAATTCCCGGACNDTCATCTTGGTGCGGTGGTTGCCTTTGGTAACAACACCTGGCGCGCTCTGAGCGGCGGCGTTGGGGCAGAAGAGCTGAAAGATTTTCCGGGCTACGGTAAAGGCCTTGCGCCGACGACCCAGTTCGATGTGTTGATCCACATTCTTTCTNDTCGTCACGACGTAAACTTCTCTGTCGCCCAGGCGGCGATGGAAGCCTTTGGTGACTGCATTGAAGTGAAAGAAGAGATCCACGGCTTCCGTTGGGTTGAAGAGCGTGACCTGAGCGGCTTTGTTGACGGTACGGAAAACCCGGCGGGTNDTGAGACGCGTCGCGAAGTGGCGGTTATCAAAGACGGCGTGGATGCGGGCGGCAGCTATGTGTTTGTCCAGCGTTGGGAACACAACCTGAAGCAGCTCAACCGGATGAGCGTTCACGATCAGGAGATGGTGATCGGGCGCACCAAAGAGGCCNDTGAAGAGATCGACGGCGACGAACGTCCGGAAACCTCTCACCTCACCCGCGTTGATCTGAAAGAAGATGGCAAAGGGCTGAAGATTGTTCGCCAGAGCCTGCCGTACGGCACTGCCAGTGGCACTCACGGTCTGTACTTCTGCGCCTACNDTGCGCGTCTGCATAACATTGAGCAGCAACTGCTGAGCATGTTTGGCGATACCGATGGTAAGCGTGATGCGATGTTGCGTTTCACCAAACCGGTAACCGGCGGCTATTATTTCGCACCGTCGCTGGACAAGTTGATGGCGCTGTAA
```

The textual output can be printed into a file.

sink("primers.txt", append=FALSE, split=FALSE)
print_primer(primers_lvl0)
sink()
