## Supplementary material for "Golden Mutagenesis: An efficient multi-site-saturation mutagenesis approach by Golden Gate cloning with automated primer design": Computer Tool example Multi-Site Directed Mutagenesis 3

Multiple Site Saturation Mutagenesis Example 3

2018-10-10

### Multiple Site Saturation Mutagenesis – Active site

You can find the lastest version of this file at <https://github.com/ipb-halle/GoldenMutagenesis/blob/master/vignettes/MSD3.md>

#### Experimental Workflow

##### Target sequence

open reading frame YfeX in pCA24N (Chloramphenicol resistance)

##### Clone into

seperate gene fragments into pAGM9121 first then reassemble into pAGM22082_cRed

##### Genomic sequence YfeX

ATGTCTCAGGTTCAGAGTGGCATTTTGCCAGAACATTGCCGCGCGGCGATTTGGATCGAAGCCAACGTGAAAGGGGAAGTTGACGCCCTGCGTGCGGCCAGTAAAACATTTGCCGACAAACTGGCAACTTTTGAAGCGAAATTCCCGGACGCGCATCTTGGTGCGGTGGTTGCCTTTGGTAACAACACCTGGCGCGCTCTGAGCGGCGGCGTTGGGGCAGAAGAGCTGAAAGATTTTCCGGGCTACGGTAAAGGCCTTGCGCCGACGACCCAGTTCGATGTGTTGATCCACATTCTTTCTCTGCGTCACGACGTAAACTTCTCTGTCGCCCAGGCGGCGATGGAAGCCTTTGGTGACTGCATTGAAGTGAAAGAAGAGATCCACGGCTTCCGTTGGGTTGAAGAGCGT**GAC**CTGAGCGGCTTTGTT**GAC**GGTACGGAA**AAC**CCGGCGGGTGAAGAGACGCGTCGCGAAGTGGCGGTTATCAAAGACGGCGTGGATGCGGGCGGCAGCTATGTGTTTGTCCAGCGTTGGGAACACAACCTGAAGCAGCTCAACCGGATGAGCGTTCACGATCAGGAGATGGTGATCGGGCGCACCAAAGAGGCCAACGAAGAGATCGACGGCGACGAACGTCCGGAAACCTCTCACCTCACCCGCGTTGATCTGAAAGAAGATGGCAAAGGGCTGAAGATTGTT**CGC**CAG**AGC**CTGCCGTACGGCACTGCCAGTGGCACTCACGGTCTGTACTTCTGCGCCTACTGCGCGCGTCTGCATAACATTGAGCAGCAACTGCTGAGCATGTTTGGCGATACCGATGGTAAGCGTGATGCGATGTTGCGTTTCACCAAACCGGTAACCGGCGGCTATTATTTCGCACCGTCGCTGGACAAGTTGATGGCGCTGTAA

##### Restriction Enzyme

###### Level 0

BbsI

Recognition site: ***GAAGAC***

###### Level 2

BsaI

Recognition site: ***GGTCTC***

##### Envisioned Mutations

Aspartic Acid - 137 - NDT
Aspartic Acid - 143 - NNN
Asparagine - 147 - DBK
Arginine - 232 - NDT
Serine - 234 - NDT

```
#If domestication is necessary follow the workflow of the Point Mutagenesis vignette
#We will use a structure similar to the Point Mutagenesis workflow to insert different kinds of Multiple Site Saturations here.
mutations<-list(c(137,"NDT"), c(143,"NNN"), c( 147, "DBK"),c(232, "NDT"), c(234, "NDT"))
primers<-msd_mutate(input_sequence, prefix="TT" ,restriction_enzyme=recognition_site_bsai, suffix="A", vector=c("AATG", "AAGC"), replacements=mutations, replacement_range=5, binding_min_length=4 ,primer_length=9, target_temp=60, fragment_min_size=60 )
primers
## An object of class "Extended Primerset"
## Slot "fragments":
## [[1]]
## An object of class "Fragment"
## Slot "start":
## [1] 2
##
## Slot "stop":
## [1] 142
##
## Slot "start_mutation":
## logical(0)
##
## Slot "stop_mutation":
## [1] 137
##
##
## [[2]]
## An object of class "Fragment"
## Slot "start":
## [1] 143
##
## Slot "stop":
## [1] 231
##
## Slot "start_mutation":
## [1] 143 147
##
## Slot "stop_mutation":
## logical(0)
##
##
## [[3]]
## An object of class "Fragment"
## Slot "start":
## [1] 232
##
## Slot "stop":
## [1] 300
##
## Slot "start_mutation":
## [1] 232 234
##
## Slot "stop_mutation":
## logical(0)
##
##
##
## Slot "oldsequence":
## [1] "ATGTCTCAGGTTCAGAGTGGCATTTTGCCAGAACATTGCCGCGCGGCGATTTGGATCGAAGCCAACGTGAAAGGGGAAGTTGACGCCCTGCGTGCGGCCAGTAAAACATTTGCCGACAAACTGGCAACTTTTGAAGCGAAATTCCCGGACGCGCATCTTGGTGCGGTGGTTGCCTTTGGTAACAACACCTGGCGCGCTCTGAGCGGCGGCGTTGGGGCAGAAGAGCTGAAAGATTTTCCGGGCTACGGTAAAGGCCTTGCGCCGACGACCCAGTTCGATGTGTTGATCCACATTCTTTCTCTGCGTCACGACGTAAACTTCTCTGTCGCCCAGGCGGCGATGGAAGCCTTTGGTGACTGCATTGAAGTGAAAGAAGAGATCCACGGCTTCCGTTGGGTTGAAGAGCGTGACCTGAGCGGCTTTGTTGACGGTACGGAAAACCCGGCGGGTGAAGAGACGCGTCGCGAAGTGGCGGTTATCAAAGACGGCGTGGATGCGGGCGGCAGCTATGTGTTTGTCCAGCGTTGGGAACACAACCTGAAGCAGCTCAACCGGATGAGCGTTCACGATCAGGAGATGGTGATCGGGCGCACCAAAGAGGCCAACGAAGAGATCGACGGCGACGAACGTCCGGAAACCTCTCACCTCACCCGCGTTGATCTGAAAGAAGATGGCAAAGGGCTGAAGATTGTTCGCCAGAGCCTGCCGTACGGCACTGCCAGTGGCACTCACGGTCTGTACTTCTGCGCCTACTGCGCGCGTCTGCATAACATTGAGCAGCAACTGCTGAGCATGTTTGGCGATACCGATGGTAAGCGTGATGCGATGTTGCGTTTCACCAAACCGGTAACCGGCGGCTATTATTTCGCACCGTCGCTGGACAAGTTGATGGCGCTGTAA"
##
## Slot "primers":
## [[1]]
## [[1]][[1]]
## An object of class "Primer"
## Slot "prefix":
## [1] "TT"
##
## Slot "restriction_enzyme":
## [1] "GGTCTC"
##
## Slot "suffix":
## [1] "A"
##
## Slot "vector":
## [1] "AATG"
##
## Slot "overhang":
## [1] ""
##
## Slot "binding_sequence":
## [1] "TCTCAGGTTCAGAGTGGCATTTTGCC"
##
## Slot "temperature":
## [1] 60.36446
##
## Slot "difference":
## [1] 0.3644563
##
##
## [[1]][[2]]
## An object of class "Primer MSD"
## Slot "NDT":
## [1] "AGCCGCTCAGAHN"
##
## Slot "prefix":
## [1] "TT"
##
## Slot "restriction_enzyme":
## [1] "GGTCTC"
##
## Slot "suffix":
## [1] "T"
##
## Slot "vector":
## [1] ""
##
## Slot "overhang":
## [1] "ACAA"
##
## Slot "binding_sequence":
## [1] "ACGCTCTTCAACCCAACGGAAGC"
##
## Slot "temperature":
## [1] 59.785
##
## Slot "difference":
## [1] 0.5794564
##
##
##
## [[2]]
## [[2]][[1]]
## An object of class "Primer MSD"
## Slot "NDT":
## [1] "TNNNGGTACGGAADBK"
##
## Slot "prefix":
## [1] "TT"
##
## Slot "restriction_enzyme":
## [1] "GGTCTC"
##
## Slot "suffix":
## [1] "A"
##
## Slot "vector":
## [1] ""
##
## Slot "overhang":
## [1] "TTGT"
##
## Slot "binding_sequence":
## [1] "CCGGCGGGTGAAGAGACGC"
##
## Slot "temperature":
## [1] 59.2169
##
## Slot "difference":
## [1] 0.7831005
##
##
## [[2]][[2]]
## An object of class "Primer"
## Slot "prefix":
## [1] "TT"
##
## Slot "restriction_enzyme":
## [1] "GGTCTC"
##
## Slot "suffix":
## [1] "T"
##
## Slot "vector":
## [1] ""
##
## Slot "overhang":
## [1] "AACA"
##
## Slot "binding_sequence":
## [1] "ATCTTCAGCCCTTTGCCATCTTCTTT"
##
## Slot "temperature":
## [1] 59.04605
##
## Slot "difference":
## [1] 0.1708527
##
##
##
## [[3]]
## [[3]][[1]]
## An object of class "Primer MSD"
## Slot "NDT":
## [1] "NDTCAGNDT"
##
## Slot "prefix":
## [1] "TT"
##
## Slot "restriction_enzyme":
## [1] "GGTCTC"
##
## Slot "suffix":
## [1] "A"
##
## Slot "vector":
## [1] ""
##
## Slot "overhang":
## [1] "TGTT"
##
## Slot "binding_sequence":
## [1] "CTGCCGTACGGCACTGCCAG"
##
## Slot "temperature":
## [1] 60.52991
##
## Slot "difference":
## [1] 0.5299114
##
##
## [[3]][[2]]
## An object of class "Primer"
## Slot "prefix":
## [1] "TT"
##
## Slot "restriction_enzyme":
## [1] "GGTCTC"
##
## Slot "suffix":
## [1] "T"
##
## Slot "vector":
## [1] "AAGC"
##
## Slot "overhang":
## [1] ""
##
## Slot "binding_sequence":
## [1] "TTACAGCGCCATCAACTTGTCCAGC"
##
## Slot "temperature":
## [1] 60.90327
##
## Slot "difference":
## [1] 0.3733614
##
##
##
##
## Slot "newsequence":
## [1] "ATGTCTCAGGTTCAGAGTGGCATTTTGCCAGAACATTGCCGCGCGGCGATTTGGATCGAAGCCAACGTGAAAGGGGAAGTTGACGCCCTGCGTGCGGCCAGTAAAACATTTGCCGACAAACTGGCAACTTTTGAAGCGAAATTCCCGGACGCGCATCTTGGTGCGGTGGTTGCCTTTGGTAACAACACCTGGCGCGCTCTGAGCGGCGGCGTTGGGGCAGAAGAGCTGAAAGATTTTCCGGGCTACGGTAAAGGCCTTGCGCCGACGACCCAGTTCGATGTGTTGATCCACATTCTTTCTCTGCGTCACGACGTAAACTTCTCTGTCGCCCAGGCGGCGATGGAAGCCTTTGGTGACTGCATTGAAGTGAAAGAAGAGATCCACGGCTTCCGTTGGGTTGAAGAGCGTNDTCTGAGCGGCTTTGTTNNNGGTACGGAADBKCCGGCGGGTGAAGAGACGCGTCGCGAAGTGGCGGTTATCAAAGACGGCGTGGATGCGGGCGGCAGCTATGTGTTTGTCCAGCGTTGGGAACACAACCTGAAGCAGCTCAACCGGATGAGCGTTCACGATCAGGAGATGGTGATCGGGCGCACCAAAGAGGCCAACGAAGAGATCGACGGCGACGAACGTCCGGAAACCTCTCACCTCACCCGCGTTGATCTGAAAGAAGATGGCAAAGGGCTGAAGATTGTTNDTCAGNDTCTGCCGTACGGCACTGCCAGTGGCACTCACGGTCTGTACTTCTGCGCCTACTGCGCGCGTCTGCATAACATTGAGCAGCAACTGCTGAGCATGTTTGGCGATACCGATGGTAAGCGTGATGCGATGTTGCGTTTCACCAAACCGGTAACCGGCGGCTATTATTTCGCACCGTCGCTGGACAAGTTGATGGCGCTGTAA"
```

The primers are generated for direct cloning into the Level 2 vector.
The function primer_add_level modifies the primers for individual cloning into Level 0 vectors and subsequent assembly in Level 2.
The parameters **prefix, restriction_enzyme, suffix and vector** can be set similar to the mutate-function.

```
primers_lvl0<-primer_add_level(primers,  prefix="TT", restriction_enzyme=recognition_site_bbsi, suffix="AA", vector=c("CTCA", "CTCG"))
primers_lvl0
## An object of class "Extended Primerset"
## Slot "fragments":
## [[1]]
## An object of class "Fragment"
## Slot "start":
## [1] 2
##
## Slot "stop":
## [1] 142
##
## Slot "start_mutation":
## logical(0)
##
## Slot "stop_mutation":
## [1] 137
##
##
## [[2]]
## An object of class "Fragment"
## Slot "start":
## [1] 143
##
## Slot "stop":
## [1] 231
##
## Slot "start_mutation":
## [1] 143 147
##
## Slot "stop_mutation":
## logical(0)
##
##
## [[3]]
## An object of class "Fragment"
## Slot "start":
## [1] 232
##
## Slot "stop":
## [1] 300
##
## Slot "start_mutation":
## [1] 232 234
##
## Slot "stop_mutation":
## logical(0)
##
##
##
## Slot "oldsequence":
## [1] "ATGTCTCAGGTTCAGAGTGGCATTTTGCCAGAACATTGCCGCGCGGCGATTTGGATCGAAGCCAACGTGAAAGGGGAAGTTGACGCCCTGCGTGCGGCCAGTAAAACATTTGCCGACAAACTGGCAACTTTTGAAGCGAAATTCCCGGACGCGCATCTTGGTGCGGTGGTTGCCTTTGGTAACAACACCTGGCGCGCTCTGAGCGGCGGCGTTGGGGCAGAAGAGCTGAAAGATTTTCCGGGCTACGGTAAAGGCCTTGCGCCGACGACCCAGTTCGATGTGTTGATCCACATTCTTTCTCTGCGTCACGACGTAAACTTCTCTGTCGCCCAGGCGGCGATGGAAGCCTTTGGTGACTGCATTGAAGTGAAAGAAGAGATCCACGGCTTCCGTTGGGTTGAAGAGCGTGACCTGAGCGGCTTTGTTGACGGTACGGAAAACCCGGCGGGTGAAGAGACGCGTCGCGAAGTGGCGGTTATCAAAGACGGCGTGGATGCGGGCGGCAGCTATGTGTTTGTCCAGCGTTGGGAACACAACCTGAAGCAGCTCAACCGGATGAGCGTTCACGATCAGGAGATGGTGATCGGGCGCACCAAAGAGGCCAACGAAGAGATCGACGGCGACGAACGTCCGGAAACCTCTCACCTCACCCGCGTTGATCTGAAAGAAGATGGCAAAGGGCTGAAGATTGTTCGCCAGAGCCTGCCGTACGGCACTGCCAGTGGCACTCACGGTCTGTACTTCTGCGCCTACTGCGCGCGTCTGCATAACATTGAGCAGCAACTGCTGAGCATGTTTGGCGATACCGATGGTAAGCGTGATGCGATGTTGCGTTTCACCAAACCGGTAACCGGCGGCTATTATTTCGCACCGTCGCTGGACAAGTTGATGGCGCTGTAA"
##
## Slot "primers":
## [[1]]
## [[1]][[1]]
## An object of class "Primer"
## Slot "prefix":
## [1] "TT"
##
## Slot "restriction_enzyme":
## [1] "GAAGAC"
##
## Slot "suffix":
## [1] "AA"
##
## Slot "vector":
## [1] "CTCA"
##
## Slot "overhang":
## [1] "AATG"
##
## Slot "binding_sequence":
## [1] "TCTCAGGTTCAGAGTGGCATTTTGCC"
##
## Slot "temperature":
## [1] 60.36446
##
## Slot "difference":
## [1] 0.3644563
##
##
## [[1]][[2]]
## An object of class "Primer MSD"
## Slot "NDT":
## [1] "AGCCGCTCAGAHN"
##
## Slot "prefix":
## [1] "TT"
##
## Slot "restriction_enzyme":
## [1] "GAAGAC"
##
## Slot "suffix":
## [1] "AA"
##
## Slot "vector":
## [1] "CTCG"
##
## Slot "overhang":
## [1] "ACAA"
##
## Slot "binding_sequence":
## [1] "ACGCTCTTCAACCCAACGGAAGC"
##
## Slot "temperature":
## [1] 59.785
##
## Slot "difference":
## [1] 0.5794564
##
##
##
## [[2]]
## [[2]][[1]]
## An object of class "Primer MSD"
## Slot "NDT":
## [1] "TNNNGGTACGGAADBK"
##
## Slot "prefix":
## [1] "TT"
##
## Slot "restriction_enzyme":
## [1] "GAAGAC"
##
## Slot "suffix":
## [1] "AA"
##
## Slot "vector":
## [1] "CTCA"
##
## Slot "overhang":
## [1] "TTGT"
##
## Slot "binding_sequence":
## [1] "CCGGCGGGTGAAGAGACGC"
##
## Slot "temperature":
## [1] 59.2169
##
## Slot "difference":
## [1] 0.7831005
##
##
## [[2]][[2]]
## An object of class "Primer"
## Slot "prefix":
## [1] "TT"
##
## Slot "restriction_enzyme":
## [1] "GAAGAC"
##
## Slot "suffix":
## [1] "AA"
##
## Slot "vector":
## [1] "CTCG"
##
## Slot "overhang":
## [1] "AACA"
##
## Slot "binding_sequence":
## [1] "ATCTTCAGCCCTTTGCCATCTTCTTT"
##
## Slot "temperature":
## [1] 59.04605
##
## Slot "difference":
## [1] 0.1708527
##
##
##
## [[3]]
## [[3]][[1]]
## An object of class "Primer MSD"
## Slot "NDT":
## [1] "NDTCAGNDT"
##
## Slot "prefix":
## [1] "TT"
##
## Slot "restriction_enzyme":
## [1] "GAAGAC"
##
## Slot "suffix":
## [1] "AA"
##
## Slot "vector":
## [1] "CTCA"
##
## Slot "overhang":
## [1] "TGTT"
##
## Slot "binding_sequence":
## [1] "CTGCCGTACGGCACTGCCAG"
##
## Slot "temperature":
## [1] 60.52991
##
## Slot "difference":
## [1] 0.5299114
##
##
## [[3]][[2]]
## An object of class "Primer"
## Slot "prefix":
## [1] "TT"
##
## Slot "restriction_enzyme":
## [1] "GAAGAC"
##
## Slot "suffix":
## [1] "AA"
##
## Slot "vector":
## [1] "CTCG"
##
## Slot "overhang":
## [1] "AAGC"
##
## Slot "binding_sequence":
## [1] "TTACAGCGCCATCAACTTGTCCAGC"
##
## Slot "temperature":
## [1] 60.90327
##
## Slot "difference":
## [1] 0.3733614
##
##
##
##
## Slot "newsequence":
## [1] "ATGTCTCAGGTTCAGAGTGGCATTTTGCCAGAACATTGCCGCGCGGCGATTTGGATCGAAGCCAACGTGAAAGGGGAAGTTGACGCCCTGCGTGCGGCCAGTAAAACATTTGCCGACAAACTGGCAACTTTTGAAGCGAAATTCCCGGACGCGCATCTTGGTGCGGTGGTTGCCTTTGGTAACAACACCTGGCGCGCTCTGAGCGGCGGCGTTGGGGCAGAAGAGCTGAAAGATTTTCCGGGCTACGGTAAAGGCCTTGCGCCGACGACCCAGTTCGATGTGTTGATCCACATTCTTTCTCTGCGTCACGACGTAAACTTCTCTGTCGCCCAGGCGGCGATGGAAGCCTTTGGTGACTGCATTGAAGTGAAAGAAGAGATCCACGGCTTCCGTTGGGTTGAAGAGCGTNDTCTGAGCGGCTTTGTTNNNGGTACGGAADBKCCGGCGGGTGAAGAGACGCGTCGCGAAGTGGCGGTTATCAAAGACGGCGTGGATGCGGGCGGCAGCTATGTGTTTGTCCAGCGTTGGGAACACAACCTGAAGCAGCTCAACCGGATGAGCGTTCACGATCAGGAGATGGTGATCGGGCGCACCAAAGAGGCCAACGAAGAGATCGACGGCGACGAACGTCCGGAAACCTCTCACCTCACCCGCGTTGATCTGAAAGAAGATGGCAAAGGGCTGAAGATTGTTNDTCAGNDTCTGCCGTACGGCACTGCCAGTGGCACTCACGGTCTGTACTTCTGCGCCTACTGCGCGCGTCTGCATAACATTGAGCAGCAACTGCTGAGCATGTTTGGCGATACCGATGGTAAGCGTGATGCGATGTTGCGTTTCACCAAACCGGTAACCGGCGGCTATTATTTCGCACCGTCGCTGGACAAGTTGATGGCGCTGTAA"
```

Objects of the classes “Primer”, “Primer MSD” and “Primerset” can have a slim textual output by using the function print_primer.

```
print_primer(primers_lvl0)
## Fragment 1
## Start 2, Stop 142, Length 141
## Forward
## TTGAAGACAACTCAAATGTCTCAGGTTCAGAGTGGCATTTTGCC
## Temperature of binding site:  60.36446  °C
## Temperature difference:  0.3644563  K
## Reverse
## TTGAAGACAACTCGACAAAGCCGCTCAGAHNACGCTCTTCAACCCAACGGAAGC
## Temperature of binding site:  59.785  °C
## Temperature difference:  0.5794564  K
##
## Fragment 2
## Start 143, Stop 231, Length 89
## Forward
## TTGAAGACAACTCATTGTTNNNGGTACGGAADBKCCGGCGGGTGAAGAGACGC
## Temperature of binding site:  59.2169  °C
## Temperature difference:  0.7831005  K
## Reverse
## TTGAAGACAACTCGAACAATCTTCAGCCCTTTGCCATCTTCTTT
## Temperature of binding site:  59.04605  °C
## Temperature difference:  0.1708527  K
##
## Fragment 3
## Start 232, Stop 300, Length 69
## Forward
## TTGAAGACAACTCATGTTNDTCAGNDTCTGCCGTACGGCACTGCCAG
## Temperature of binding site:  60.52991  °C
## Temperature difference:  0.5299114  K
## Reverse
## TTGAAGACAACTCGAAGCTTACAGCGCCATCAACTTGTCCAGC
## Temperature of binding site:  60.90327  °C
## Temperature difference:  0.3733614  K
##
## Input Sequence:
##  ATGTCTCAGGTTCAGAGTGGCATTTTGCCAGAACATTGCCGCGCGGCGATTTGGATCGAAGCCAACGTGAAAGGGGAAGTTGACGCCCTGCGTGCGGCCAGTAAAACATTTGCCGACAAACTGGCAACTTTTGAAGCGAAATTCCCGGACGCGCATCTTGGTGCGGTGGTTGCCTTTGGTAACAACACCTGGCGCGCTCTGAGCGGCGGCGTTGGGGCAGAAGAGCTGAAAGATTTTCCGGGCTACGGTAAAGGCCTTGCGCCGACGACCCAGTTCGATGTGTTGATCCACATTCTTTCTCTGCGTCACGACGTAAACTTCTCTGTCGCCCAGGCGGCGATGGAAGCCTTTGGTGACTGCATTGAAGTGAAAGAAGAGATCCACGGCTTCCGTTGGGTTGAAGAGCGTGACCTGAGCGGCTTTGTTGACGGTACGGAAAACCCGGCGGGTGAAGAGACGCGTCGCGAAGTGGCGGTTATCAAAGACGGCGTGGATGCGGGCGGCAGCTATGTGTTTGTCCAGCGTTGGGAACACAACCTGAAGCAGCTCAACCGGATGAGCGTTCACGATCAGGAGATGGTGATCGGGCGCACCAAAGAGGCCAACGAAGAGATCGACGGCGACGAACGTCCGGAAACCTCTCACCTCACCCGCGTTGATCTGAAAGAAGATGGCAAAGGGCTGAAGATTGTTCGCCAGAGCCTGCCGTACGGCACTGCCAGTGGCACTCACGGTCTGTACTTCTGCGCCTACTGCGCGCGTCTGCATAACATTGAGCAGCAACTGCTGAGCATGTTTGGCGATACCGATGGTAAGCGTGATGCGATGTTGCGTTTCACCAAACCGGTAACCGGCGGCTATTATTTCGCACCGTCGCTGGACAAGTTGATGGCGCTGTAA
##
## Modified Sequence:
##  ATGTCTCAGGTTCAGAGTGGCATTTTGCCAGAACATTGCCGCGCGGCGATTTGGATCGAAGCCAACGTGAAAGGGGAAGTTGACGCCCTGCGTGCGGCCAGTAAAACATTTGCCGACAAACTGGCAACTTTTGAAGCGAAATTCCCGGACGCGCATCTTGGTGCGGTGGTTGCCTTTGGTAACAACACCTGGCGCGCTCTGAGCGGCGGCGTTGGGGCAGAAGAGCTGAAAGATTTTCCGGGCTACGGTAAAGGCCTTGCGCCGACGACCCAGTTCGATGTGTTGATCCACATTCTTTCTCTGCGTCACGACGTAAACTTCTCTGTCGCCCAGGCGGCGATGGAAGCCTTTGGTGACTGCATTGAAGTGAAAGAAGAGATCCACGGCTTCCGTTGGGTTGAAGAGCGTNDTCTGAGCGGCTTTGTTNNNGGTACGGAADBKCCGGCGGGTGAAGAGACGCGTCGCGAAGTGGCGGTTATCAAAGACGGCGTGGATGCGGGCGGCAGCTATGTGTTTGTCCAGCGTTGGGAACACAACCTGAAGCAGCTCAACCGGATGAGCGTTCACGATCAGGAGATGGTGATCGGGCGCACCAAAGAGGCCAACGAAGAGATCGACGGCGACGAACGTCCGGAAACCTCTCACCTCACCCGCGTTGATCTGAAAGAAGATGGCAAAGGGCTGAAGATTGTTNDTCAGNDTCTGCCGTACGGCACTGCCAGTGGCACTCACGGTCTGTACTTCTGCGCCTACTGCGCGCGTCTGCATAACATTGAGCAGCAACTGCTGAGCATGTTTGGCGATACCGATGGTAAGCGTGATGCGATGTTGCGTTTCACCAAACCGGTAACCGGCGGCTATTATTTCGCACCGTCGCTGGACAAGTTGATGGCGCTGTAA
```

The textual output can be printed into a file.

sink("primers.txt", append=FALSE, split=FALSE)
print_primer(primers_lvl0)
sink()
