## Supplementary material for "Golden Mutagenesis: An efficient multi-site-saturation mutagenesis approach by Golden Gate cloning with automated primer design": Computer Tool example point mutagenesis and domestication

2018-10-10

### Point Mutagenesis

You can find the lastest version of this file at <https://github.com/ipb-halle/GoldenMutagenesis/blob/master/vignettes/Point_Mutagenesis.md>

#### Experimental Workflow

##### Target sequence

open reading frame mCherry in pet28a (Kanamycin resistance)

##### Clone into

pAGM9121

##### Genomic sequence mCherry

ATGGTGAGCAAGGGCGAGGAGGATAACATGGCCATCATCAAGGAGTTCATGCGCTTCAAGGTGCACATGGAGGGCTCCG TGAACGGCCACGAGTTCGAGATCGAGGGCGAGGGCGAGGGCCGCCCCTACGAGGGCACCCAGACCGCCAAGCTGAAGGT GACCAAGGGTGGCCCCCTGCCCTTCGCCTGGGACATC**CTG**TCCCCTCAGTTCATGTACGGCTCCAAGGCCTACGT GAAGCACCCCGCCGACATCCCCGACTACTTGAAGCTGTCCTTCCCCGAGGGCTTCAAGTGGGAGCGCGTGATGAACTTC GAGGACGGCGGCGTGGTGACCGTGACCCAGGACTCCTCCCTGCAGGACGGCGAGTTCATCTACAAGGTGAAGCTGCGCG GCACCAACTTCCCCTCCGACGGCCCCGTAATGCAGAA***GAAGAC***GATGGGCTGGGAGGCCTCCTCCGAGCGGATGT ACCCCGAGGACGGCGCCCTGAAGGGCGAGATCAAGCAGAGGCTGAAGCTGAAGGACGGCGGCCACTACGACGCTGAGGT CAAGACCACCTACAAGGCCAAGAAGCCCGTGCAGCTGCCCGGCGCCTACAACGTCAACATCAAGTTGGACATCACCTCC CACAACGAGGACTACACCATCGTGGAACAGTACGAACGCGCCGAGGGCCGCCACTCCACCGGCGGCATGGACGAGCTGT ACAAGGTCGACAAGCTTGCGGCCGCACTCGAGTGA

##### Restriction Enzyme

BbsI

Recognition site: ***GAAGAC***

##### Envisioned Mutation

Leucine66 to Valine (point mutation)

**CTG** -> **GTG**

#### R Workflow

suppressWarnings(suppressMessages(library("GoldenMutagenesis")))

input_sequence<-"ATGGTGAGCAAGGGCGAGGAGGATAACATGGCCATCATCAAGGAGTTCATGCGCTTCAAGGTGCACATGGAGGGCTCCGTGAACGGCCACGAGTTCGAGATCGAGGGCGAGGGCGAGGGCCGCCCCTACGAGGGCACCCAGACCGCCAAGCTGAAGGTGACCAAGGGTGGCCCCCTGCCCTTCGCCTGGGACATCCTGTCCCCTCAGTTCATGTACGGCTCCAAGGCCTACGTGAAGCACCCCGCCGACATCCCCGACTACTTGAAGCTGTCCTTCCCCGAGGGCTTCAAGTGGGAGCGCGTGATGAACTTCGAGGACGGCGGCGTGGTGACCGTGACCCAGGACTCCTCCCTGCAGGACGGCGAGTTCATCTACAAGGTGAAGCTGCGCGGCACCAACTTCCCCTCCGACGGCCCCGTAATGCAGAAGAAGACGATGGGCTGGGAGGCCTCCTCCGAGCGGATGTACCCCGAGGACGGCGCCCTGAAGGGCGAGATCAAGCAGAGGCTGAAGCTGAAGGACGGCGGCCACTACGACGCTGAGGTCAAGACCACCTACAAGGCCAAGAAGCCCGTGCAGCTGCCCGGCGCCTACAACGTCAACATCAAGTTGGACATCACCTCCCACAACGAGGACTACACCATCGTGGAACAGTACGAACGCGCCGAGGGCCGCCACTCCACCGGCGGCATGGACGAGCTGTACAAGGTCGACAAGCTTGCGGCCGCACTCGAGTGA"
recognition_site_bbsi<-"GAAGAC"
recognition_site_bsai<-"GGTCTC"
cuf<-"e_coli_316407.csv"

The domesticate function checks for internal cleavage sites. If corresponding sites are present silent mutations are introduced to destroy the recognition sites. The functions returns a list containing the position of the choosen amino acid residue for silent mutation.

```
mutations_bbsi<-domesticate(input_sequence, recognition_site_bbsi, cuf)
mutations_bbsi
## [[1]]
## [1] "143" "K"
mutations_bsai<-domesticate(input_sequence, recognition_site_bsai, cuf)
## [1] "No domestication needed."
mutations_bsai
## list()
```

The mutate function designs the necessary set of primers for the desired mutations. 
The function has the following parameters:
**prefix**: Additional nucleobases in 5’ position of the recognition site [default: TT]
**restriction_enzyme**: Recognition site sequence of the respective restriction enzyme [default: GGTCTC]
**suffix**: Spacer nucleotides matching the cleavage pattern of the enzyme [default: A]
**vector**: Four basepair overhangs complementary to the created overhangs in the acceptor vector [default: c(“AATG”, “AAGC”)]
**replacements**: The desired substitutions
**primer_min_length**: The minimal threshold value of the template binding sequence [default: 4]
**primer_length**: Maximal length of the binding sequence [default: 9]
**target_temp**: Melting temperature of the binding sequence in °C [default: 60]
**cuf**: The Codon Usage Table which is being used to select the codon for an exchanged amino acid. [default: e_coli_316407.csv]

It will return an object of the class primer_set.

```
mutations<-c(list(c(66, "V")), mutations_bbsi)
primers<-mutate(input_sequence, prefix="TT", restriction_enzyme = recognition_site_bbsi, suffix = "AA", vector=c("CTCA", "CTCG"), replacements = mutations, binding_min_length=4 ,primer_length=9, target_temp=60, cuf=cuf)
primers
## An object of class "Primerset"
## Slot "oldsequence":
## [1] "ATGGTGAGCAAGGGCGAGGAGGATAACATGGCCATCATCAAGGAGTTCATGCGCTTCAAGGTGCACATGGAGGGCTCCGTGAACGGCCACGAGTTCGAGATCGAGGGCGAGGGCGAGGGCCGCCCCTACGAGGGCACCCAGACCGCCAAGCTGAAGGTGACCAAGGGTGGCCCCCTGCCCTTCGCCTGGGACATCCTGTCCCCTCAGTTCATGTACGGCTCCAAGGCCTACGTGAAGCACCCCGCCGACATCCCCGACTACTTGAAGCTGTCCTTCCCCGAGGGCTTCAAGTGGGAGCGCGTGATGAACTTCGAGGACGGCGGCGTGGTGACCGTGACCCAGGACTCCTCCCTGCAGGACGGCGAGTTCATCTACAAGGTGAAGCTGCGCGGCACCAACTTCCCCTCCGACGGCCCCGTAATGCAGAAGAAGACGATGGGCTGGGAGGCCTCCTCCGAGCGGATGTACCCCGAGGACGGCGCCCTGAAGGGCGAGATCAAGCAGAGGCTGAAGCTGAAGGACGGCGGCCACTACGACGCTGAGGTCAAGACCACCTACAAGGCCAAGAAGCCCGTGCAGCTGCCCGGCGCCTACAACGTCAACATCAAGTTGGACATCACCTCCCACAACGAGGACTACACCATCGTGGAACAGTACGAACGCGCCGAGGGCCGCCACTCCACCGGCGGCATGGACGAGCTGTACAAGGTCGACAAGCTTGCGGCCGCACTCGAGTGA"
##
## Slot "primers":
## [[1]]
## [[1]][[1]]
## An object of class "Primer"
## Slot "prefix":
## [1] "TT"
##
## Slot "restriction_enzyme":
## [1] "GAAGAC"
##
## Slot "suffix":
## [1] "AA"
##
## Slot "vector":
## [1] "CTCA"
##
## Slot "overhang":
## [1] ""
##
## Slot "binding_sequence":
## [1] "ATGGTGAGCAAGGGCGAGGAGG"
##
## Slot "temperature":
## [1] 60.67248
##
## Slot "difference":
## [1] 0.6724812
##
##
## [[1]][[2]]
## An object of class "Primer"
## Slot "prefix":
## [1] "TT"
##
## Slot "restriction_enzyme":
## [1] "GAAGAC"
##
## Slot "suffix":
## [1] "AA"
##
## Slot "vector":
## [1] ""
##
## Slot "overhang":
## [1] "CACG"
##
## Slot "binding_sequence":
## [1] "ATGTCCCAGGCGAAGGGCAGGG"
##
## Slot "temperature":
## [1] 61.41454
##
## Slot "difference":
## [1] 0.7420589
##
##
##
## [[2]]
## [[2]][[1]]
## An object of class "Primer"
## Slot "prefix":
## [1] "TT"
##
## Slot "restriction_enzyme":
## [1] "GAAGAC"
##
## Slot "suffix":
## [1] "AA"
##
## Slot "vector":
## [1] ""
##
## Slot "overhang":
## [1] "CGTG"
##
## Slot "binding_sequence":
## [1] "TCCCCTCAGTTCATGTACGGCTCC"
##
## Slot "temperature":
## [1] 59.45263
##
## Slot "difference":
## [1] 0.5473745
##
##
## [[2]][[2]]
## An object of class "Primer"
## Slot "prefix":
## [1] "TT"
##
## Slot "restriction_enzyme":
## [1] "GAAGAC"
##
## Slot "suffix":
## [1] "AA"
##
## Slot "vector":
## [1] ""
##
## Slot "overhang":
## [1] "TTTC"
##
## Slot "binding_sequence":
## [1] "TGCATTACGGGGCCGTCGGA"
##
## Slot "temperature":
## [1] 58.9345
##
## Slot "difference":
## [1] 0.5181218
##
##
##
## [[3]]
## [[3]][[1]]
## An object of class "Primer"
## Slot "prefix":
## [1] "TT"
##
## Slot "restriction_enzyme":
## [1] "GAAGAC"
##
## Slot "suffix":
## [1] "AA"
##
## Slot "vector":
## [1] ""
##
## Slot "overhang":
## [1] "GAAA"
##
## Slot "binding_sequence":
## [1] "AAGACGATGGGCTGGGAGGCC"
##
## Slot "temperature":
## [1] 59.2929
##
## Slot "difference":
## [1] 0.7071014
##
##
## [[3]][[2]]
## An object of class "Primer"
## Slot "prefix":
## [1] "TT"
##
## Slot "restriction_enzyme":
## [1] "GAAGAC"
##
## Slot "suffix":
## [1] "AA"
##
## Slot "vector":
## [1] "CTCG"
##
## Slot "overhang":
## [1] ""
##
## Slot "binding_sequence":
## [1] "TCACTCGAGTGCGGCCGC"
##
## Slot "temperature":
## [1] 59.15354
##
## Slot "difference":
## [1] 0.1393632
##
##
##
##
## Slot "newsequence":
## [1] "ATGGTGAGCAAGGGCGAGGAGGATAACATGGCCATCATCAAGGAGTTCATGCGCTTCAAGGTGCACATGGAGGGCTCCGTGAACGGCCACGAGTTCGAGATCGAGGGCGAGGGCGAGGGCCGCCCCTACGAGGGCACCCAGACCGCCAAGCTGAAGGTGACCAAGGGTGGCCCCCTGCCCTTCGCCTGGGACATCGTGTCCCCTCAGTTCATGTACGGCTCCAAGGCCTACGTGAAGCACCCCGCCGACATCCCCGACTACTTGAAGCTGTCCTTCCCCGAGGGCTTCAAGTGGGAGCGCGTGATGAACTTCGAGGACGGCGGCGTGGTGACCGTGACCCAGGACTCCTCCCTGCAGGACGGCGAGTTCATCTACAAGGTGAAGCTGCGCGGCACCAACTTCCCCTCCGACGGCCCCGTAATGCAGAAAAAGACGATGGGCTGGGAGGCCTCCTCCGAGCGGATGTACCCCGAGGACGGCGCCCTGAAGGGCGAGATCAAGCAGAGGCTGAAGCTGAAGGACGGCGGCCACTACGACGCTGAGGTCAAGACCACCTACAAGGCCAAGAAGCCCGTGCAGCTGCCCGGCGCCTACAACGTCAACATCAAGTTGGACATCACCTCCCACAACGAGGACTACACCATCGTGGAACAGTACGAACGCGCCGAGGGCCGCCACTCCACCGGCGGCATGGACGAGCTGTACAAGGTCGACAAGCTTGCGGCCGCACTCGAGTGA"
```

Objects of the classes “primer”, “primer_msd” and “primer_set” can have a slim textual output by using the function print_primer.

```
print_primer(primers)
## Fragment 1
## Forward
## TTGAAGACAACTCAATGGTGAGCAAGGGCGAGGAGG
## Temperature of binding site:  60.67248  °C
## Temperature difference:  0.6724812  K
## Reverse
## TTGAAGACAACACGATGTCCCAGGCGAAGGGCAGGG
## Temperature of binding site:  61.41454  °C
## Temperature difference:  0.7420589  K
##
## Fragment 2
## Forward
## TTGAAGACAACGTGTCCCCTCAGTTCATGTACGGCTCC
## Temperature of binding site:  59.45263  °C
## Temperature difference:  0.5473745  K
## Reverse
## TTGAAGACAATTTCTGCATTACGGGGCCGTCGGA
## Temperature of binding site:  58.9345  °C
## Temperature difference:  0.5181218  K
##
## Fragment 3
## Forward
## TTGAAGACAAGAAAAAGACGATGGGCTGGGAGGCC
## Temperature of binding site:  59.2929  °C
## Temperature difference:  0.7071014  K
## Reverse
## TTGAAGACAACTCGTCACTCGAGTGCGGCCGC
## Temperature of binding site:  59.15354  °C
## Temperature difference:  0.1393632  K
##
## Input Sequence:
##  ATGGTGAGCAAGGGCGAGGAGGATAACATGGCCATCATCAAGGAGTTCATGCGCTTCAAGGTGCACATGGAGGGCTCCGTGAACGGCCACGAGTTCGAGATCGAGGGCGAGGGCGAGGGCCGCCCCTACGAGGGCACCCAGACCGCCAAGCTGAAGGTGACCAAGGGTGGCCCCCTGCCCTTCGCCTGGGACATCCTGTCCCCTCAGTTCATGTACGGCTCCAAGGCCTACGTGAAGCACCCCGCCGACATCCCCGACTACTTGAAGCTGTCCTTCCCCGAGGGCTTCAAGTGGGAGCGCGTGATGAACTTCGAGGACGGCGGCGTGGTGACCGTGACCCAGGACTCCTCCCTGCAGGACGGCGAGTTCATCTACAAGGTGAAGCTGCGCGGCACCAACTTCCCCTCCGACGGCCCCGTAATGCAGAAGAAGACGATGGGCTGGGAGGCCTCCTCCGAGCGGATGTACCCCGAGGACGGCGCCCTGAAGGGCGAGATCAAGCAGAGGCTGAAGCTGAAGGACGGCGGCCACTACGACGCTGAGGTCAAGACCACCTACAAGGCCAAGAAGCCCGTGCAGCTGCCCGGCGCCTACAACGTCAACATCAAGTTGGACATCACCTCCCACAACGAGGACTACACCATCGTGGAACAGTACGAACGCGCCGAGGGCCGCCACTCCACCGGCGGCATGGACGAGCTGTACAAGGTCGACAAGCTTGCGGCCGCACTCGAGTGA
##
## Modified Sequence:
##  ATGGTGAGCAAGGGCGAGGAGGATAACATGGCCATCATCAAGGAGTTCATGCGCTTCAAGGTGCACATGGAGGGCTCCGTGAACGGCCACGAGTTCGAGATCGAGGGCGAGGGCGAGGGCCGCCCCTACGAGGGCACCCAGACCGCCAAGCTGAAGGTGACCAAGGGTGGCCCCCTGCCCTTCGCCTGGGACATCGTGTCCCCTCAGTTCATGTACGGCTCCAAGGCCTACGTGAAGCACCCCGCCGACATCCCCGACTACTTGAAGCTGTCCTTCCCCGAGGGCTTCAAGTGGGAGCGCGTGATGAACTTCGAGGACGGCGGCGTGGTGACCGTGACCCAGGACTCCTCCCTGCAGGACGGCGAGTTCATCTACAAGGTGAAGCTGCGCGGCACCAACTTCCCCTCCGACGGCCCCGTAATGCAGAAAAAGACGATGGGCTGGGAGGCCTCCTCCGAGCGGATGTACCCCGAGGACGGCGCCCTGAAGGGCGAGATCAAGCAGAGGCTGAAGCTGAAGGACGGCGGCCACTACGACGCTGAGGTCAAGACCACCTACAAGGCCAAGAAGCCCGTGCAGCTGCCCGGCGCCTACAACGTCAACATCAAGTTGGACATCACCTCCCACAACGAGGACTACACCATCGTGGAACAGTACGAACGCGCCGAGGGCCGCCACTCCACCGGCGGCATGGACGAGCTGTACAAGGTCGACAAGCTTGCGGCCGCACTCGAGTGA
```

The textual output can be printed into a file.

sink("primers.txt", append=FALSE, split=FALSE)
print_primer(primers)
sink()
