## Supplementary material for "Golden Mutagenesis: An efficient multi-site-saturation mutagenesis approach by Golden Gate cloning with automated primer design": Computer Tool example Quick Quality Control

2018-10-10

### Quick Quality Control

You can find the lastest version of this file at <https://github.com/ipb-halle/GoldenMutagenesis/blob/master/vignettes/QQC.md>

#### Experimental Workflow

##### Target sequence

open reading frame YfeX in pCA24N (Chloramphenicol resistance)

##### Clone into

seperate gene fragments into pAGM9121 first then reassemble into pAGM22082_cRed

mutations<-c(137,143,147,232,234)

##### Quality Control

The functions aligns the obtained sequencing results to the target gene sequence. It also tries to align the reverse complement of the obtained sequence. Afterwards it checks for mismatches between the sequences. Mismatches are likely to be sucessfully mutated nucleotides. Positions regarded as mismatches are displayed as pie charts. The shown distributions are based on the signal intensities of the four nucleobases at the mismatch positions. You can compare the pie charts with expected pattern of randomization, therefore validating the quality of the created library.

###### Forward Sequencing

abfile<-"sequences/Yfex_0activesite_for_EF01147142.ab1")
base_distribution(input_sequence=input_sequence, ab1file=abfile, replacements=mutations)

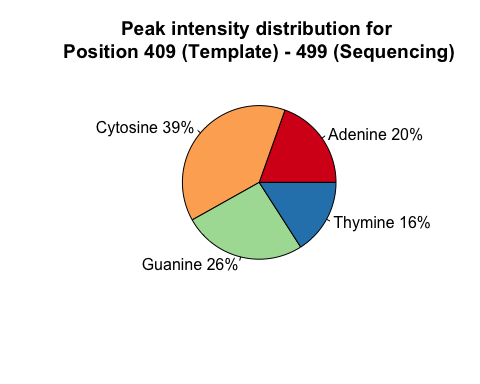

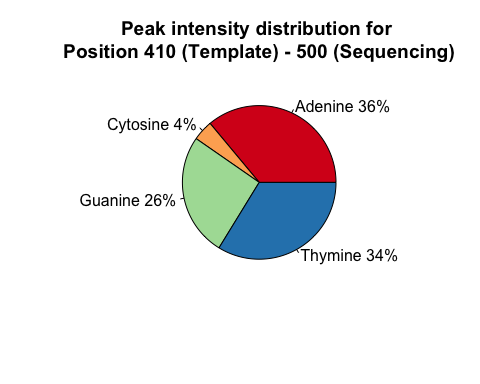

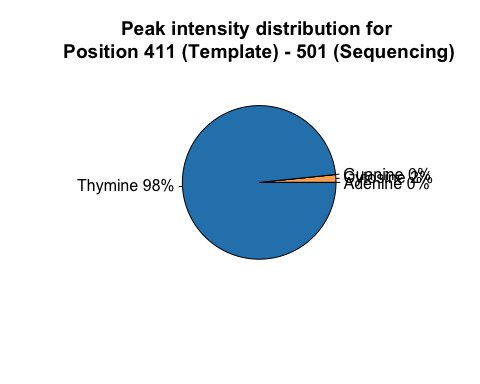

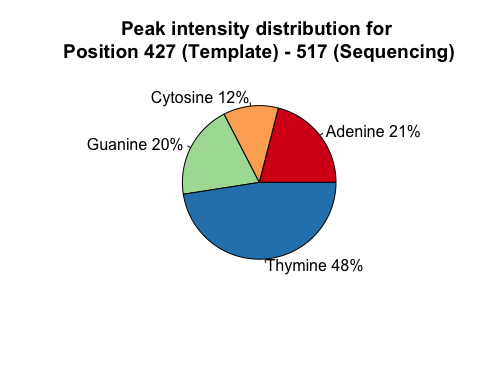

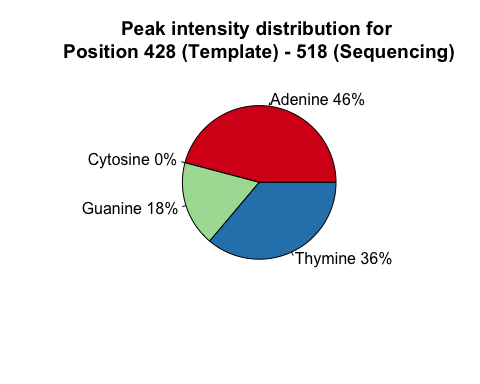

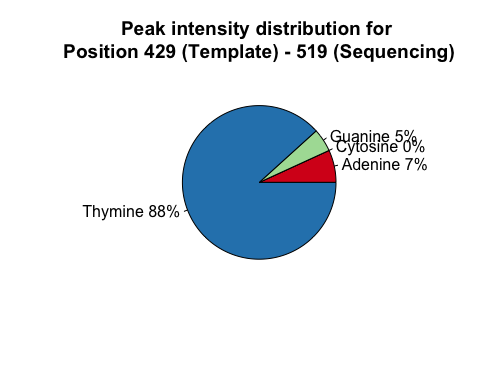

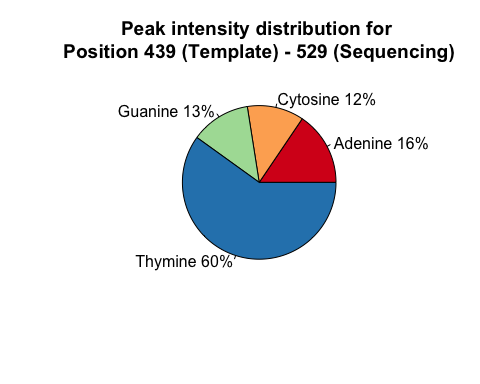

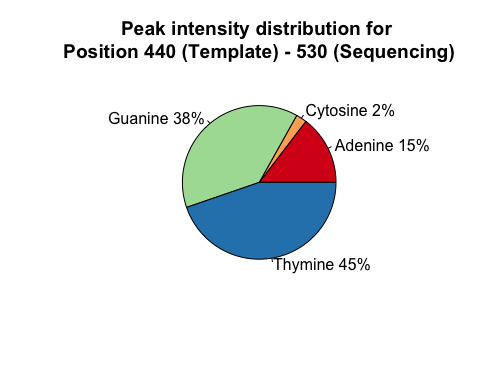

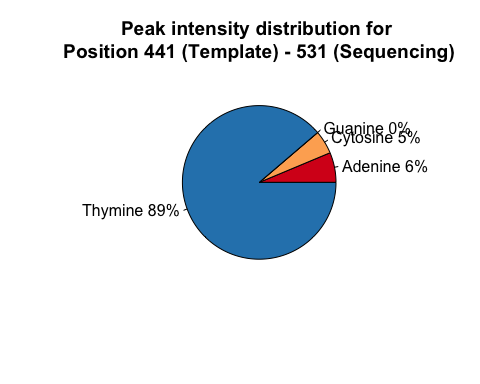

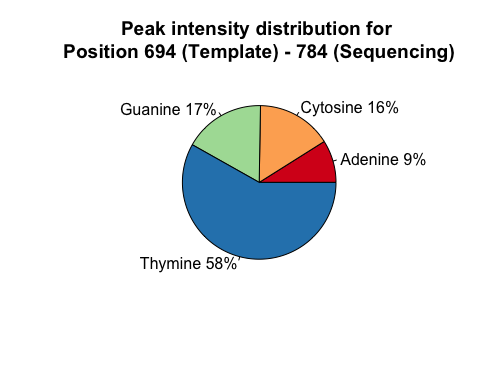

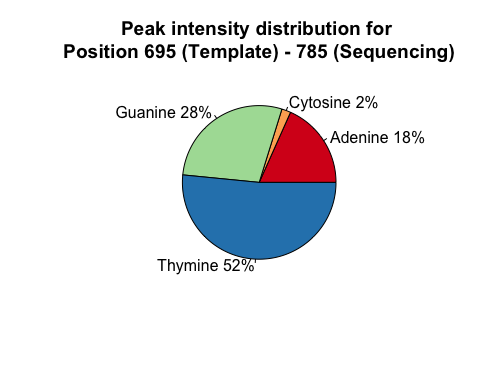

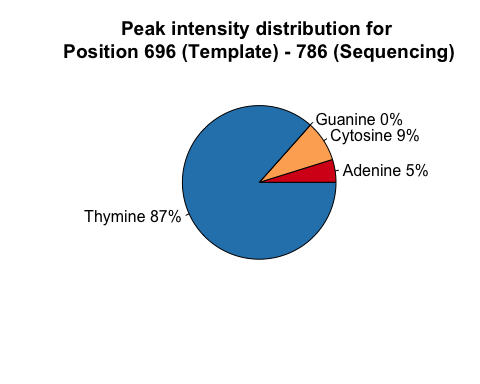

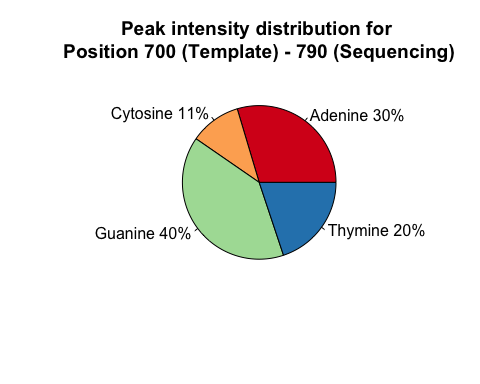

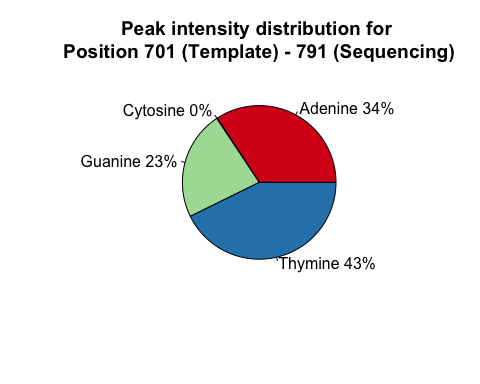

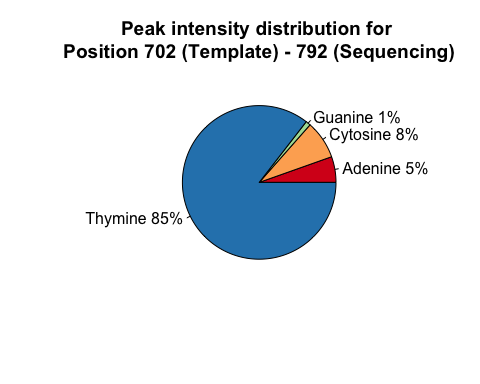

###### Reverse Sequencing

```
abfile<-"sequences/Yfex_activesite_rev_EF01147143.ab1")
base_distribution(input_sequence=input_sequence, ab1file=abfile, replacements=mutations)
## [1] "Reverse sequence detected!"
```

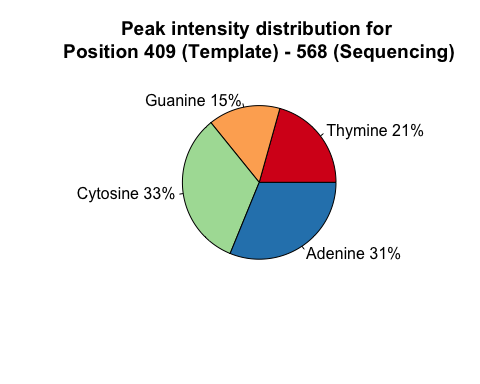

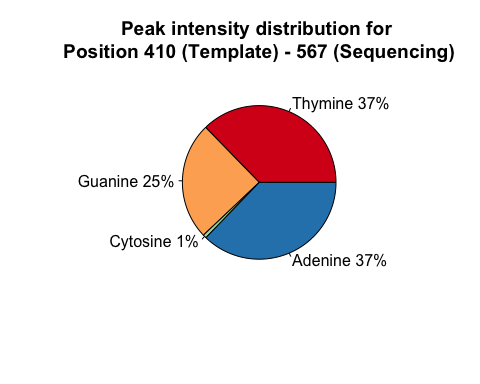

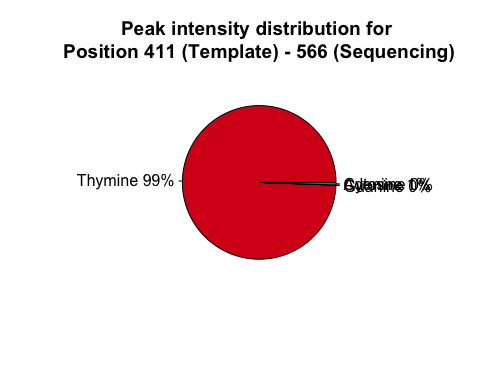

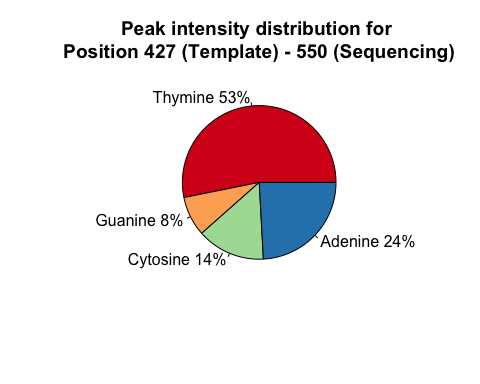

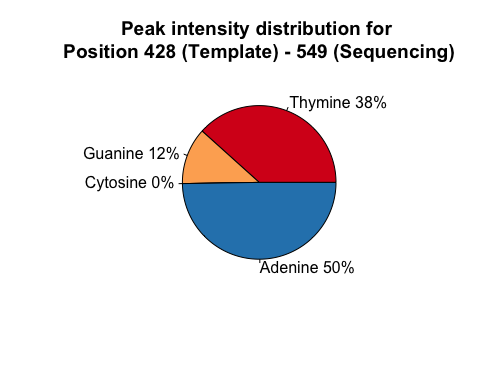

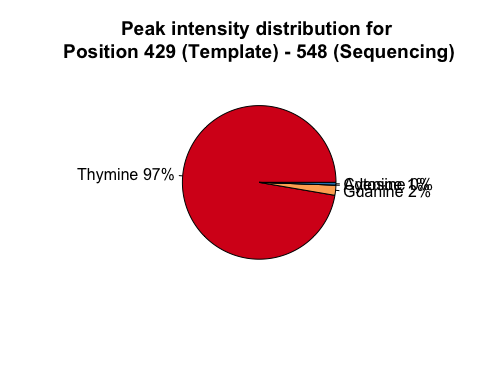

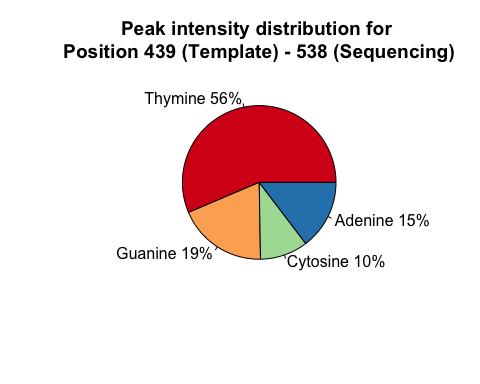

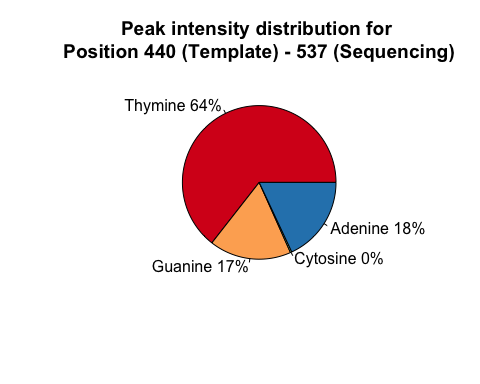

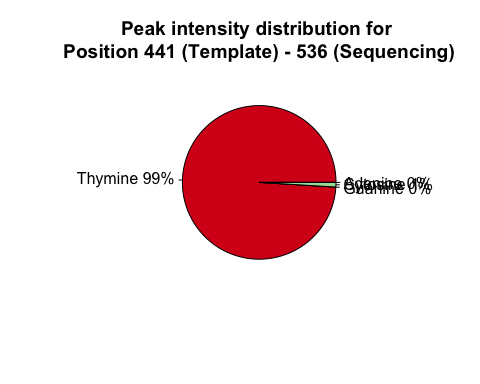

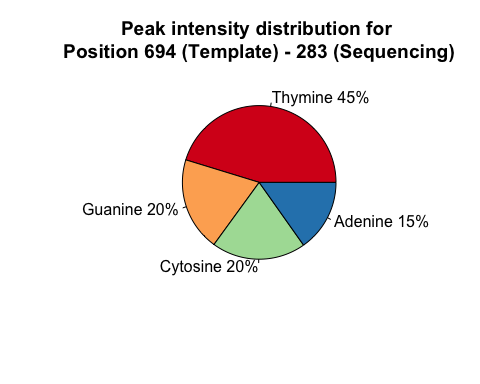

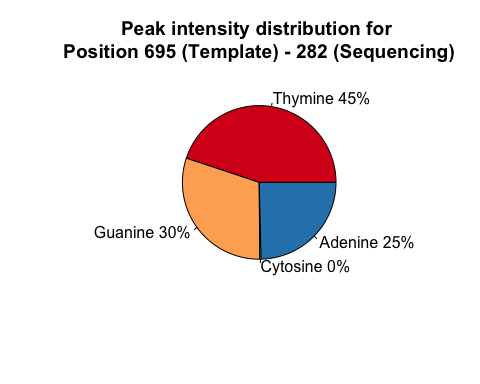

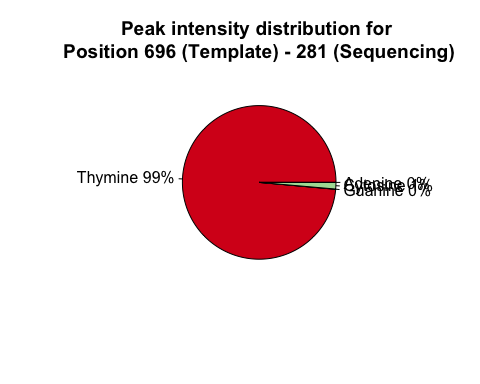

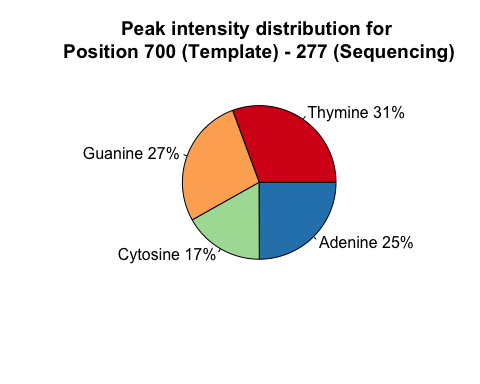

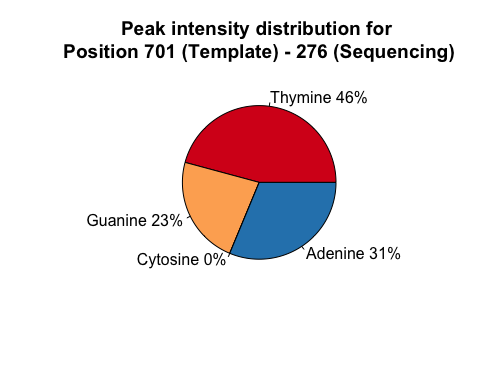

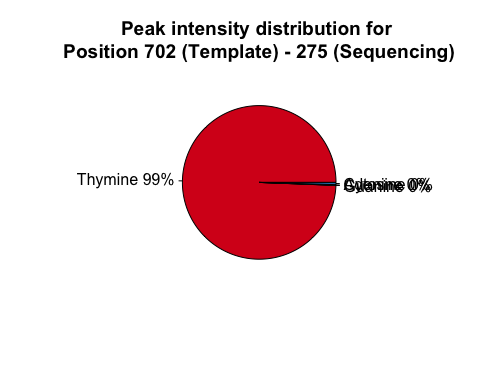
